# Cross-species comparative connectomics reveals the evolution of an olfactory circuit

**DOI:** 10.1101/2025.06.11.659158

**Authors:** Ruairí J.V. Roberts, Christoph Giez, Serene Dhawan, Song Pang, Nadine Randel, Zhiyuan Lu, Hui Gong, Léonard Dekens, James Di Frisco, C. Shan Xu, Harald F. Hess, Marta Zlatic, Albert Cardona, Lucia L. Prieto-Godino

## Abstract

Animal behavioural diversity ultimately stems from variation in neural circuitry, yet how central neural circuits evolve remains poorly understood. Studies of neural circuit evolution often focus on a few elements within a network. However, addressing fundamental questions in evolutionary neuroscience, such as whether some elements are more evolvable than others, requires a more global and unbiased approach. Here, we used synapse-level comparative connectomics to examine how an entire olfactory circuit evolves. We compared the full antennal lobe connectome of the larvae of two closely related *Drosophila* species, *D. melanogaster* and *D. erecta*, which differ in their ecological niches and odour-driven behaviours. We found that evolutionary change is unevenly distributed across the network. Some features, including neuron types, neuron numbers and interneuron-to-interneuron connectivity, are highly conserved. These conserved elements delineate a core circuit blueprint presumably required for fundamental olfactory processing. Superimposed on this scaffold, we find rewiring changes that mirror each species’ ecologies, including a systematic shift in the excitation-to-inhibition balance in the feedforward pathways. We further show that some neurons have changed more than others, and that even within individual neurons some synaptic elements remain conserved while others display major species-specific changes, suggesting evolutionary hot-spots within the circuit. Our findings reveal constrained and adaptable elements within olfactory networks, and establish a framework for identifying general principles in the evolution of neural circuits underlying behaviour.

## Introduction

The astonishing diversity of animals’ sensory driven behaviours is a direct consequence of evolution shaping neural circuits. However, in contrast to our rapidly increasing understanding of how sensory information is processed in the brain, we know far less about how neural circuits evolve to generate species-specific behaviours. Some progress has been made by focusing on specific neurons within sensory circuits^1–8^. However, this strategy rarely offers the global perspective required to address fundamental questions about the way neural circuits evolve, for example, whether some elements are more evolvable than others.

An ideal way to gain such a systematic outlook is to examine complete circuits at synaptic resolution in species with different behaviours by using comparative connectomics. This approach was pioneered by comparing the neural circuits of two distantly related nematode species^9–11^, providing important insights into neuronal evolution. However, it remains unclear how broadly these findings apply across the diverse nervous system architectures found in other animals. Additionally, given the large phylogenetic distance between the two nematode species examined, the question of how neural circuits evolve over shorter evolutionary timescales remains open.

The olfactory system of the *Drosophila* larva is an attractive model because it shares core organisational features with olfactory circuits of other clades, including vertebrates, yet its smaller size unlocks the potential for a complete, synaptic level characterisation of a complex glomerular olfactory network through connectomics^12^.

A key feature of glomerular olfactory systems, whether in vertebrates or invertebrates, is that the axons of olfactory sensory neurons (OSNs) that express the same receptors project to a common region within the first olfactory relay centre: the antennal lobe in insects and olfactory bulb in vertebrates. These regions, defined by the molecular identity of the OSNs, are called glomeruli and are the functional units of olfactory circuits^13–15^. As with most neural networks, the information from these structurally organised inputs is processed by local interneurons and sent to other brain regions via output projections neurons (Fig. 1a). Some of the computations performed by local neurons are common across phyla and networks, such as divisive normalisation^12,16,17^. Therefore, studying olfactory circuits can provide insights widely applicable to other neural networks. At the same time, their organisation into clearly defined functional units – the glomeruli – facilitates the interpretation of function out of structure, making them ideal systems for connectomics.

**Figure 1.**
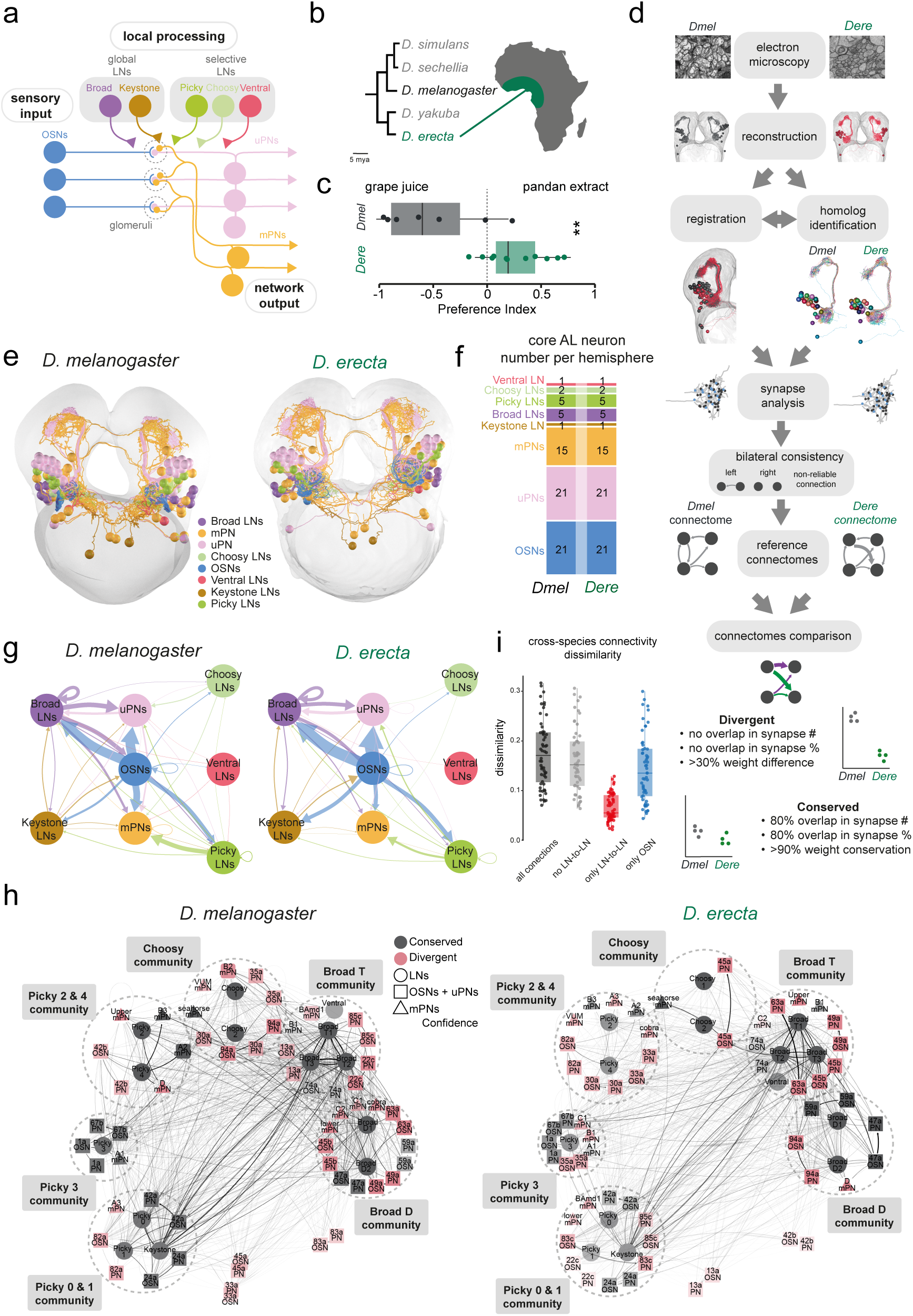
Comparative connectomics between *D. melanogaster* and *D. erecta*. **a,** Schematic of the olfactory system of the Drosophilid larva indicating each of the main neuronal classes. Olfactory sensory neurons (OSN, blue), uniglomerular projection neurons (uPN, pink), multiglomerular projection neurons (mPNs, yellow), local inhibitory neurons can be classified into global and selective LNs. Global LNs: Broad LNs (purple) and Keystone LNs (brown), Selective LNs: Choosy LNs (light green), Picky LNs (dark green) and Ventral LNs (dark pink). **b,** Evolutionary tree showing the placement of *D. melanogaster* – cosmopolitan generalist – and *D. erecta* – endemic to West Africa and specialist in pandan fruits (top). **c,** Behavioural response of the first instar larvae of *D. melanogaster* and *D. erecta* in a two choice assay with grape juice and pandan extract. Asterisks indelicate statistical significance (p < 0.01, Wilcoxon test). **d,** Schematic of the working flow. Starting from electron microscopy volumes for *D. melanogaster* ^32^, and *D. erecta* (this work), we manually traced neurons and synapses of the olfactory circuit. Reconstructed circuits were then registered onto a common template, and homologous neurons identified within and across species. Next, synapses were analysed without any filtering, following by the identification of reliable connections as those found across the two hemispheres of each dataset in at least 2 synapses or more. This defined the reference *D. melanogaster* and *D. erecta* connectomes. Next, we compared the connectomes across species, defining divergent and conserved connections as shown in figure and methods. **e,** Neuronal reconstructions of all the main elements of the antennal lobe circuit: OSNs (blue), uPNs (pink), mPNs (yellow), and the five classes of LNs – broad (magenta), choosy (light green), ventral (fuscia), keystone (brown) and picky (dark green). **f,** Quantification of the number of core AL neurons across species. **g,** General connectivity between the main classes of neurons within the AL is very similar in both species. **h,** Communities of neurons identified by the Louvain method of modularity maximization^66^ when run on the networks formed by AL core neurons (see Methods). The algorithm independently identified the same six communities in both species (delineated by dashed lines), defined by the intra-LN connectivity: Broad D, Broad T, Choosy, Picky 2 & 4, Picky 3 and Picky 0 & 1 communities. Two additional communities in each species with two neurons each were also identified. The neurons belonging to these communities are different across species. In black are those neurons that belong to the same community across both species, in red are those that in each species has been assigned to a different community. The algorithm was run 1000 times. The opacity of each neuron indicates confidence of community assignment, or the number of times in the 1,000 runs that a given neuron was assigned to the community shown in the figure. LNs are indicated with circles, OSNs and uPNs with squares and mPNs with triangles. **i,** Quantification of the dissimilarity across species for each of the neurons in the core AL network (each of the dots in the plot is the comparison of each of the identified one to one homologs of each species) as measured by Euclidean distance. This calculation was done for the real connectomes, as well as connectomes where we experimentally removed *in-silico* either all LN-to-LN connections, all connections but LN-to-LN, or all connections but those coming from OSNs.

Although the organisational principles of olfactory circuits are similar across large evolutionary distances, the degree to which the network changes at the synaptic level across species as these adapt to different ecological niches remains largely unknown. Drosophilids are a powerful model clade to investigate the evolution of neural circuits because of the divergent behaviours displayed by closely related species^18^ and the insights that can be gained by comparing them to the well-characterised circuits of *D. melanogaster*^1–4,7,8,19–23^. Although most studies have focused on adult flies, *Drosophila* larvae have a complex olfactory-guided behavioural repertoire that allows them to navigate the heterogeneous ever-changing environments of decaying fruits^15,24–27^. Importantly, the larvae of *D. erecta*, a close relative of *D. melanogaster*, has divergent odour-guided behaviours^28^ (Fig.1b-c), akin to differences reported for adults^29,30^. *D. erecta*, like its relative *D. melanogaster*, originated in Africa; while *D. melanogaster* has since become a cosmopolitan generalist – feeding and breeding on a wide range of fruits – *D. erecta* is endemic to West Africa and is specialised to exclusively feed and breed on the green and woody fruits of the shrub *Pandanus*^29,30^.

To address questions about how neural circuits evolve in an unbiased and integrated way, we here provide, to our knowledge, the first synaptic resolution cross-species comparative connectomic analysis of a complete glomerularly organised olfactory network. We compare the full antennal lobe circuitry of the larvae of two closely related *Drosophila* species with divergent ecologies and odour-guided behaviours. We find that both species share a common circuit blueprint, including an identical complement of homologous neurons. Accordingly, direct comparison of how the *same* neurons connect in *new* ways, enables us to investigate the evolution of neuronal connectivity at the level of individually identified neurons.

We find that evolutionary change is unevenly distributed across the network. Some connectivity features, such as the interneuron-to-interneuron connectivity, are strikingly conserved, pointing to their key role in odour processing across species. These “stable” circuits are embedded in a suite of changes, including a systematic shift in the balance of excitation to inhibition in the feedforward network in a way that matches each species’ ecologies. Remarkably, we uncover that not all circuit elements have evolved their connectivity to the same extent, and that even within single neurons some synaptic elements remain conserved while others display major species-specific changes. Our work highlights how evolution can differentially impinge on the elements of a neural circuit with synaptic precision and suggests that some elements might act as hot-spots for behavioural change.

## Results

### Comparing olfactory circuits across species at synaptic resolution

We acquired, using electron microscopy, a synaptic resolution volume of the central nervous system of a first instar *D. erecta* larva that encompasses the whole brain and part of the ventral nerve cord (Methods). We bilaterally reconstructed all olfactory sensory neurons (OSNs) and their synaptic partners and compared these reconstructions to a corresponding *D. melanogaster* connectome^12,31^ (Fig. 1d).

We find that *D. erecta* has the exact same neuronal complement as *D. melanogaster^12^,* including 21 OSNs connecting one to one with each of the 21 uniglomerular projection neurons (uPNs), constituting the feedforward uniglomerular system. In addition, information is carried to higher brain centres by a second set of 15 multiglomerular projections neurons (mPNs). Olfactory information is processed in both species by five distinct classes of local inhibitory interneurons (LNs), classified as: global LNs – encompassing the classes Broad (5 neurons per hemisphere) and keystone (2 medial bilaterally projecting neurons) –, and selective LNs – consisting of the classes Choosy (2 neurons per hemisphere), Picky (5 per hemisphere) and ventral (1 per hemisphere)^12^ (Fig. 1a,e-f). This conservation points at changes in connectivity, rather than neuronal cell types, as the key substrate for behavioural change across closely related species.

To examine differences in connectivity with cellular and synaptic resolution, we next established homologies for each of the 21 uPNs (Methods), thus defining 21 homologous glomeruli across species (Fig. 1d and Fig. S1). Employing similar methods, we established homologies for all the neurons in the circuit (Fig. 1d-g). In the following sections we analyse their connectivity. Features conserved across species likely represent elements required for core computations in olfactory processing, while species-specific differences might reflect adaptations of each species to their respective ecological niche. Throughout the manuscript we have applied a stringent framework to define species-specific features, favouring a conservative approach that robustly identifies both, conserved connections as well as cross-species changes (Methods). Briefly, based on previous work, we use bilateral symmetry to estimate within species variability. Connections represented by more than 2 synapses in each hemisphere were shown in the *D. melanogaster* larva to be species-characteristic connections that can be found reliably between the same homologous identifiable neurons across individuals^32^, and furthermore those representing more than 1% of input/output connectivity are considered to be strong and conserved within a species ^31,33,34^. In addition, we attach a divergence confidence score to every cross-species difference, and identify strongly conserved connections (Fig. 1d, Methods). Therefore, for each cross-species change in connectivity we report the magnitude of the difference and its confidence.

### A common circuit blueprint

We clustered all LNs of both species based on their total connectivity and found that homologous neurons clustered together as expected, closely matching their clustering based on morphological similarity (Fig. S2). This result confirms our assignment of homologs across species (which was agnostic to neuronal connectivity) and establishes a conserved circuit blueprint of the AL across species. OSNs provide input to all circuit elements and receive input mostly from Broad and Keystone LNs. The two output channels of the AL, uPNs and mPNs, receive their main inputs from OSNs followed by Broad LNs (for uPNs) and Picky LNs (for mPNs) (Fig. 1g).

To uncover general features of this common circuit blueprint, we first determined the selectivity of each of the LN classes by calculating their cumulative glomerular synapse score^35^ (see Methods, Fig. S2b). We further characterise each LN class by measuring its input/output characteristics (Fig. S2c) and its influence in the information flow through the network (Fig, S2d-e). The later calculated with two metrics: out-degree – a measure of how much a neuron influences its postsynaptic partners, and local reaching centrality – a measure of how far the activity of a neuron spreads through the network (see Methods). We found that these quantitative features delineate each LN class and are conserved across species, with some subtle differences (Fig. S2).

Next, we investigated how the network architecture of the AL has evolved by using modularity analysis to identify groups of neurons with high connectivity within groups and lower connectivity across groups. Although the network is not strongly modular, this approach independently identified the same 6 communities across both species (Fig. 1h, see Methods). These communities are delineated by the interneuron to interneuron (LN-to-LN) connectivity, highlighting strong conservation in the local interneuron network. Interestingly, OSN-uPN pairs always cluster together in the same community but their association to specific communities has changed across species (Supplementary table 1). These observations suggest that the interneuron to interneuron (LN-to-LN) connectivity has remained relatively conserved while the way in which each of the olfactory channels plug into this backbone network has changed to a greater extent. We confirmed these conclusions by calculating the connectivity dissimilarity for each neuron in the core AL network and then *in-silico* removing either all LN-to-LN connectivity, all connectivity but LN-to-LN, or all connectivity but those originating from OSN (Fig. 1i). This revealed that LN-to-LN connectivity is more similar across species than both, the overall connectivity of the AL and the connectivity from OSNs (Fig. 1i). This uncovers a first principle in the evolution of olfactory circuits: conservation of the general blueprint of the local inhibitory network with divergence of the connectivity from individual olfactory channels.

In the following sections we delve deeper into the conserved and divergent connectivity features, tackling each element of the circuit in turn. First, we examine the uniglomerular circuit, OSNs and uPNs, followed by each of the LNs and the mPNs.

### Evolution of the excitatory versus inhibitory input balance at the first synapse of olfaction

The shortest feedforward pathway of an olfactory circuit is the connection from OSNs to their glomerular partner uPNs (Fig. 2a). We calculated the anatomical transfer function^34^ of this synapse per glomerulus and surprisingly found that in each species a different set of OSNs are either strongly or weakly connected to their partner uPN (Fig. 2b). This cannot be explained solely by compensatory mechanisms adjusting OSN outputs to maintain a constant synaptic density on uPNs with larger dendrites^36^, because in that case we would expect these patterns to disappear when normalising by uPN dendritic cable length (Fig. 2c and S3a-b). Importantly, whether looking at OSN synaptic number, or normalising by uPN inputs or cable length, three OSNs – all predicted to detect leafy odours (82a, 63a and 22c, Methods and Supplementary table 2) – are more strongly connected to their corresponding uPNs in *D. erecta* (Fig.2b-c and S3a-b). This matches *D. erecta’s* ecology, given that pandan is a green, leafy, woody fruit. Furthermore, of these, 82a OSNs encode with high sensitivity the presence of geranyl acetate, a major component in the volatiles of pandan fruit^37^. Our results, therefore, indicate specific increases in *D. erecta’s* OSN-to-uPN transfer function, potentially increasing the reliability of the signal for ecologically key odours^36,38^.

**Figure 2.**
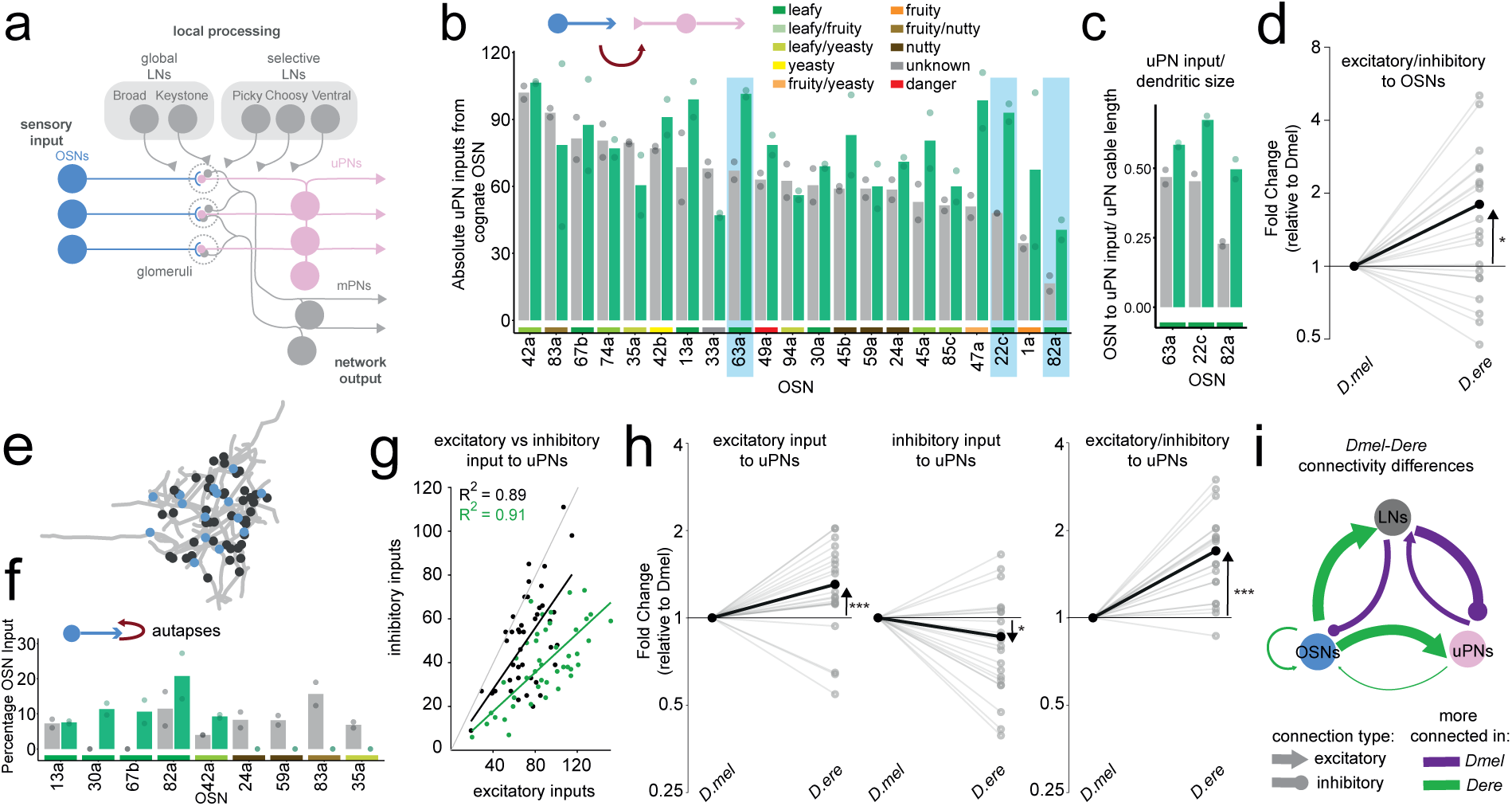
Evolution of excitatory vs inhibitory balance in the uniglomerular circuit. **a,** Schematic of the olfactory system of the Drosophilid larva indicating each of the main neuronal classes, highlighting OSNs (blue) and uPNs (pink). **b,** Number of synapses onto each uPN from its cognate OSN in *D. melanogaster* (black) and *D. erecta* (green). Bars are the average between the two circles that represent left and right hemispheres of each species. **c,** uPN inputs from its cognate OSN normalised uPN cable length for the indicated uPNs. **d,** Evolution of the ratio of excitation to inhibition onto OSNs across species. Fold change of excitatory vs inhibitory inputs in *D. erecta* when compared to *D. melanogaster*. Each grey dot is an OSN, black represents the mean. The asterisk represents significance p = 0.048. Wilcoxon Rank test. **e,** *D. erecta* 82a OSN axonal terminals (black) with circles labelling autapses (blue). **f,** percentage of OSN autapses of total input for the indicated OSNs. Legend as in b. **g,** Correlation of excitatory and inhibitory inputs onto each uPN per species. D. melanogaster, black R^2^ = 0.897, D. erecta, green R^2^= 0.914. **h,** Ratio of excitation to inhibition onto uPNs across species. Fold change of excitatory inputs, inhibitory inputs and their ratio onto uPNs when compared to *D. melanogaster*. Each grey dot is a uPN, black represents the mean. The asterisks represent significance *** p = 0.045 x10^-3^, * p = 0.02, *** p = 0.017 x 10^-8^. Wilcoxon Rank test. **i,** Network diagram of the difference in connectivity between OSNs, uPNs and LNs. Thickness of the arrows indicates the difference in number of synaptic connections between the two species. Green means more connected in *D. erecta* and magenta more connected in *D. melanogaster*.

In sensory circuits, the feedforward signal from sensory neurons is typically shaped through presynaptic inhibition by local interneurons, in the larval olfactory circuit this function is largely carried out by the Broad LNs^12,39^ (Fig. S3c). We found that *D. erecta* OSNs receive less Broad LN inputs than *D. melanogaster*, despite the former having slightly longer cable length and synaptic output (Fig. S3d). This results in lower presynaptic inhibition (Fig. S3e) and a net increase in the excitation vs inhibition ratio in the terminals of OSNs in *D. erecta* when compared to *D. melanogaster* (Fig. 2d, Fig. S3e). This could be an adaptation that in the specialist, *D. erecta,* increases signal sensitivity by reducing noise introduced by local inhibitory neurons^40,41^ – facilitating navigation within a single fruit -, while in the generalist, *D. melanogaster,* favours signal discriminability^17,42^ across multiple substrates.

Surprisingly, we found that some OSNs reliably synapse onto themselves – and occasionally other OSNs. Such autapses are also present in *D. melanogaster* larval OSNs, although they have not previously been reported (Fig. 2e-f, Fig. S3g-h, see Methods). These synapses represent a small fraction of total OSN input (on average 9.8% and 12.2% for *Dmel* and *Dere* respectively) but can be high for some OSNs, up to 27% for 82a OSN in *D. erecta*. Furthermore, this connectivity motif has also evolved in a way that correlates with each species’ ecology: In *D. erecta,* autapses are restricted to OSNs detecting leafy and fruity odours, while in *D. melanogaster* are more evenly distributed, mostly happening in yeasty and nutty OSNs (Fig. 2f). Excitatory autapses can implement positive feedback loops that sustain activity beyond the time a neuron would be active based on its input activation alone^43–45^. Furthermore, in the adult *Drosophila* antennal lobe, it has been shown that OSN-to-OSN excitatory synapses can mediate decorrelation of odour responses and increase odour discriminability^46^. Therefore, the differences in OSN autapses might serve as a mechanism to amplify signals of ecologically relevant channels, leading to increased decorrelation and discriminability of these odour representations. Furthermore, this axo-axonic connectivity establishes larval OSNs as the first processing layer of olfactory information, even before information is transferred to the local interneuron network.

### Increased excitation vs inhibition in the specialist *D. erecta* is a feature of the uniglomerular circuit

We next examined the second neuronal class of the uniglomerular circuit, the uPNs, main downstream partners of OSNs and one of the two output channels of the AL (Fig. 2a). The dendrites of uPNs integrate excitatory input from OSNs and lateral inhibition from LNs, mostly Broad LNs followed by Choosy LNs (Fig. S4a). We found that excitatory and inhibitory inputs onto uPN dendrites are correlated within each species (Fig. 2g), indicating that the dendrites of each uPN keep a specific balance. In both species excitation dominates over inhibition, a pattern that is significantly stronger for *D. erecta* (Fig. 2g). This is due to a combination of both, increased excitation and decreased inhibition onto uPNs in the specialist, leading to a greater overall excitatory to inhibitory (E/I) ratio (Fig. 2h).

Therefore, both uniglomerular circuit elements – OSNs and uPNs – display a higher E/I input ratio in *D. erecta*. The different levels of lateral inhibition in this pathway might have evolved to match the needs of each species’ ecologies, with the generalist maximising gain control and discriminability and the specialist optimising sensitivity.

Furthermore, uPN dendrites also make output synapses, including towards OSNs, with a higher proportion in *D. erecta* (Fig. 2i and S4b). This is expected to create a positive feedback loop that is stronger in the specialist, which further contributes to the overall increased excitatory drive of their uniglomerular pathway, potentially contributing to improved sensitivity.

### Global inhibitory network conservation with evolution of synaptic weight distributions

Given the observed differences in the balance of excitatory vs inhibitory input to OSNs and uPNs, we next examined the connectivity of the main global inhibitory interneurons of the AL, Broad LNs (Fig. 3a). These interneurons innervate all or most glomeruli without distinguishable dendritic/axonal compartments, as revealed by their glomerular synapse scores, which are the highest of all LNs for both species, and their low segregation index (Fig. S2b-c).

**Figure 3.**
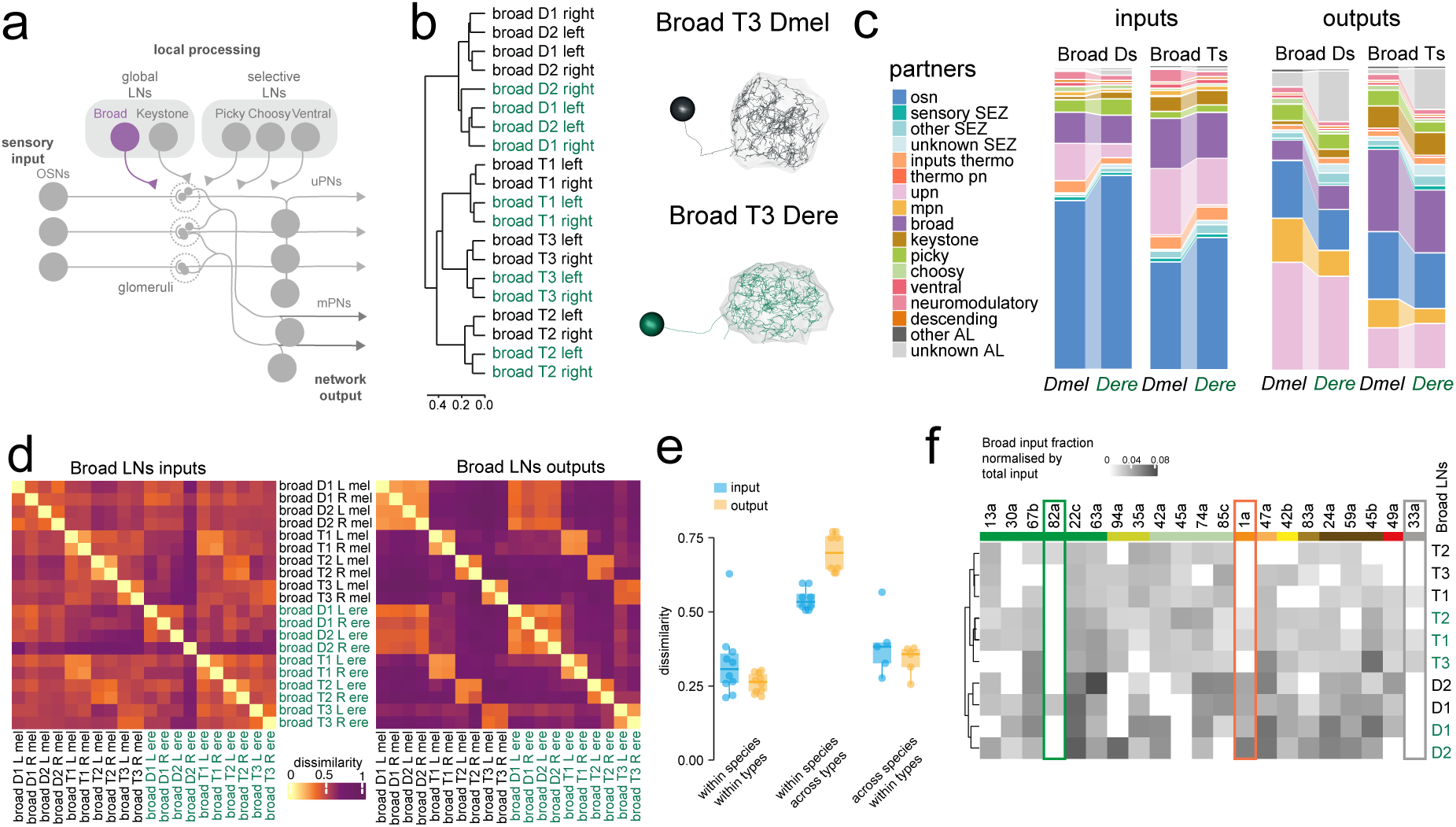
The main global inhibitory circuit: Broad LNs. **a,** Schematic of the olfactory system of the Drosophilid larva indicating each of the main neuronal classes, highlighting Broad LNs (purple). **b,** Clustering of Broad LNs of both species based on total connectivity (homologous neurons only) after filtering for connections consisting of 2 or more synapses in both hemispheres of a species. (left) Illustration of Broad LNs morphology, Broad T3 left in *D. melanogaster* (black) and *D. erecta* (green). **c,** Fraction of total Broad LN inputs (left) and outputs (right) to/from the indicated neuronal classes for each species. **d,** Dissimilarity matrix of Broad LNs based on the Euclidean distances of the vectors formed by the normalised inputs (right) or outputs (left) onto/from Broad LNs from/to all neurons for which homologs were identified across both species. **e,** Quantification of the dissimilarity matrix, showing the dissimilarity values when comparing Broad LNs as indicated: within types, across types, or within types across species. Connectivity is similar within types and across species, but different across types, particularly in the outputs, due to the intra Broad T connectivity (see Fig. S9a). **f,** Heatmap of Broad LN input fraction of total connectivity coming from each of the OSNs. Only connections that constitute at least 1% of total input connectivity in both hemispheres of a species are shown. Three glomeruli that are discussed in the main text are highlighted.

We found that general connectivity features of Broad LNs are well conserved. This includes a clear separation of this class into two types: Broad Ds and Broad Ts (Fig. 3b)^12^. The main synaptic partners of both types are OSNs and uPNs. However, the synaptic weight contribution for each type is different. Broad Ds receive most of their inputs from OSNs and provide feedforward postsynaptic inhibition mainly to uPNs, a circuit motif that shapes uPN responses to the derivative of OSN activation^12,42^ (Fig. 3c and Fig. S5-S7). Broad Ts receive a relatively stronger input contribution from uPNs and provide stronger presynaptic inhibition to OSNs, implementing gain control^12,47^ (Fig. 3c and Fig. S7a). Both operations contribute to the decorrelation glomerular responses and increase odour discriminability^17,42^. The key function in odour signal processing of these connectivity motifs likely underlies their conservation, while evolution has tuned their levels in a species-specific fashion (Fig. 2 and Fig 3c). In particular, *D. erecta* Broad LNs generally receive more inputs from OSNs and fewer from uPNs than *D. melanogaster* (Fig. 2i and 3c and Fig. S7). The functional consequences of this change are unclear, but it could confer the *D. erecta* Broad LN circuit faster odour dynamics. On the output side, as shown before, Broad LNs in *D. erecta* send on average fewer outputs to OSNs than in *D. melanogaster* (Fig. 2i and 3c, and Fig. S7-S8), a change we speculate could reflect different evolutionary trade-offs in sensitivity and discriminability between the specialist and the generalist species.

To examine the evolution of synaptic connectivity at a finer scale beyond neuronal classes, we calculated the input and output dissimilarity matrices (Fig. 3d and S9-S10). Broad LNs share many inputs within types and across species, with only slightly more differences when comparing across types (i.e. Broad Ds vs Ts) (Fig. 3d-e and S9). In contrast, the output dissimilarity matrix has more structure, revealing common outputs for Broad Ds, and conserved characteristic connectivity for each Broad T LN type, leading to comparable dissimilarity within types and species but larger across type dissimilarity (Fig. 3d-e). This pattern is driven by reciprocal inhibition among Broad Ts in both species, and removing intra-Broad T connectivity breaks the pattern and reduces across types dissimilarity (Fig. 3e and Fig. S8a), suggesting that depending on the combination of activated glomeruli, one of the three Broad Ts might dominate presynaptic inhibition in the AL.

Superimposed on these general patterns, we found that although these local interneurons sample across all glomeruli (Fig. S5, S7 and S9), the strength of the connectivity with specific OSNs and uPNs is heterogeneous, with some strongly conserved connections and some species-specific weight changes (Fig. 3f and Fig. S7-S8). Most prominently, glomerulus 82a – whose OSNs express a high sensitivity receptor for a pandan volatile –seems to operate relatively independently of the Broad LN circuit in *D. erecta*. In this species Broad LNs do not receive any strong inputs from either 82a OSN or uPN and only send weak outputs to 82a OSN but not 82a uPN (Fig. 3f and S7-S8). This might be an adaptation to reduce the noise and increase the reliability of this important ecological signal^40^. Similarly, in *D. melanogaster*, Broad LNs are less connected to the 1a glomerulus, which is narrowly tuned to fruity scents.

Interestingly, a third presumably narrowly tuned glomerulus, the orphan 33a^48^, has low input and output synapses to and from Broad LNs across both species. These results suggest that the Broad LN inhibitory circuits have likely been under purifying selection to maintain a core computational function, while at the same time some aspects of their connectivity have evolved to facilitate navigation in each species-specific environment.

### Conserved co-regulation of two global inhibitory circuits

Together with Broad LNs, Keystone LNs constitute the second component of the AL global inhibitory network^12^ (Fig. 4a). These bilateral interneurons receive input from most glomeruli but, unlike Broad LNs, they provide inhibition in a more heterogeneous way across the AL, as reflected by their lower output glomerular scores (Fig. 4b-c and Fig. S2b). A conserved feature across species is that even though Keystone LNs have a lower number of downstream partners than Broad LNs (i.e. a lower outdegree), the strength of their influence over the AL network dynamics might be similar, as reflected in their comparable local reaching centrality scores (Fig. 4c and Fig. S2d-e).

**Figure 4.**
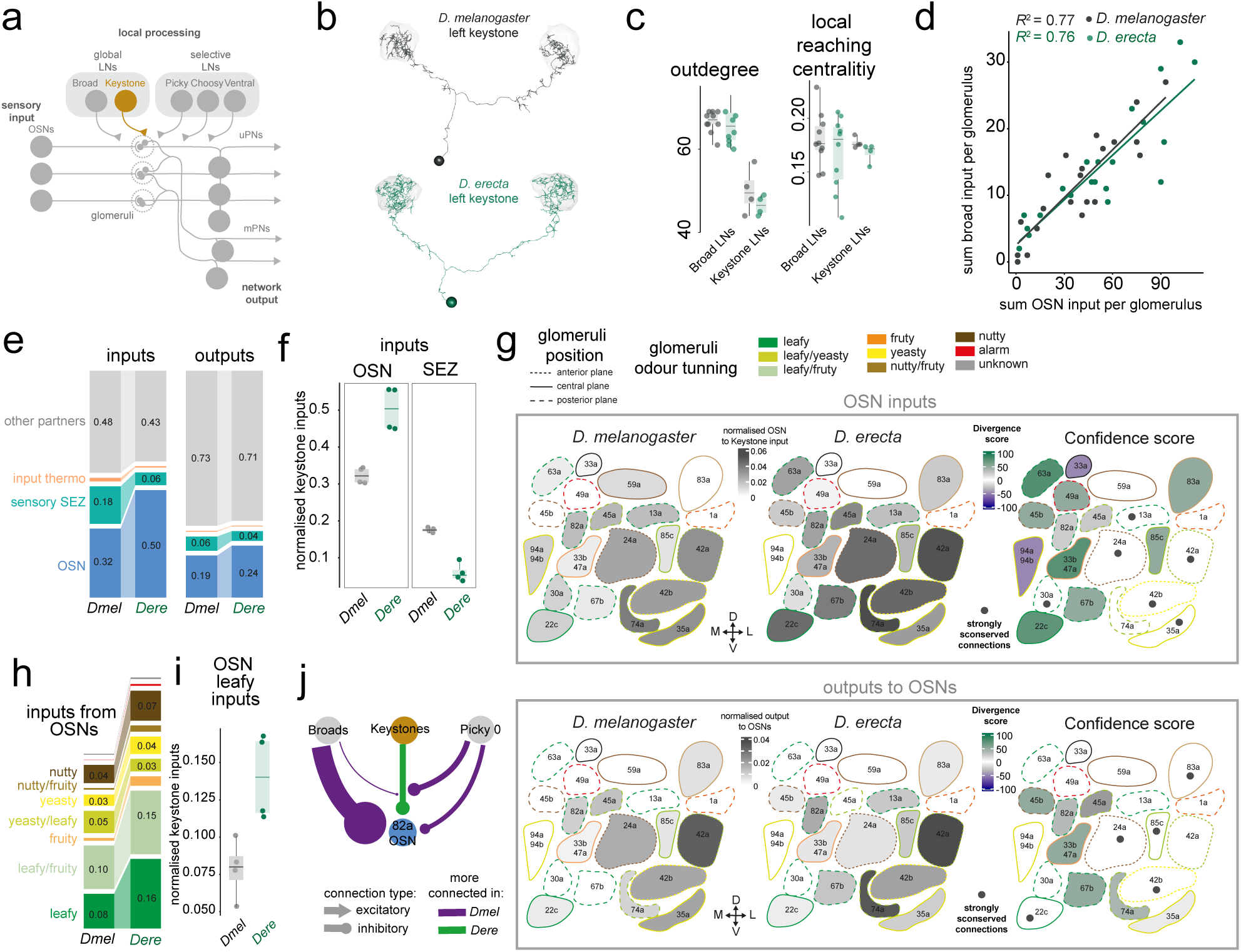
The second global inhibitory network: Keystone LNs. **a,** Schematic of the olfactory system of the Drosophilid larva indicating each of the main neuronal classes, highlighting Keystone LNs (brown). **b,** Keystone morphology in both species. **c,** Calculation of outdegree (left) and local reaching centrality (LRC, right) for Broad LNs and Keystone LNs. Neurons with high out-degree (strong output connections) have strong effects on the network, neurons with high LRC can reach many nodes in the network. **d,** Correlation between the OSN input and the Broad LN input received by Keystone LNs in each glomerulus. Each dot represents the sum of synapses to both Keystone LNs in the left and right glomeruli. The correlation coefficient R^2^ = 0.7 and 0.72 for *D. melanogaster* and *D. erecta* respectively. **e,** Fraction of total Keystone LN inputs (left) and outputs (right) to/from the indicated neuronal classes for each species. **f,** Individual datapoints (each a branch of a Keystone LN) showing the faction of Keystone LN inputs from OSNs and SEZ sensory inputs. **g,** Fraction of total Keystone inputs (top) and outputs (bottom) from/to each OSN. Only connections consisting of 2 or more synapses in both hemispheres of a species are shown. The connectivity of the two Keystones has been averaged. To the left in both plots the divergence confidence score (see methods) of the differences in OSN input to Keystone LNs between the two species, indicated as dots are those connections that are strongly conserved (see methods). The glomerular map indicates the relative position of each glomerulus. The Anterior (A) – Posterior (P) and Dorso (D) – Ventral (V) axis are indicated at the top left inset. Those with solid lines are located in the centre of the AL, those with small dashed lines are located anteriorly and those with larger dashed lines are located posteriorly. The colour of the outline depicts the ecological odour tuning of the OSN innervating each glomeruli (Table S2). **h,** Fraction of OSN inputs (left) and outputs (right) to/from Keystone LNs grouped by the ecological significance of the chemicals the OSNs detect (see methods). **i,** Individual datapoints (each a branch of a Keystone LN) showing the faction of Keystone LN inputs from leafy OSNs. **j,** Network diagram of the difference in input connectivity from Broad LNs and Picky LN 0 into Keystone LNs and 82a OSNs within the 82a glomerulus. The thickness of the connections indicates the difference in total number of synapses, but is scaled differently for the input to each neuron. Magenta means more connected in *D. melanogaster* and green means more connected in *D. erecta*.

In both species, Keystone LNs and Broad LNs inhibit each other (Fig. 1g, S5-S6, and S11-S13), a connectivity motif that had been suggested to regulate whether presynaptic inhibition is dominated by one or the other^12^. However, two lines of evidence argue against this interpretation. First, Keystone LN neurites are mixed input/output elements (Fig. S2c), and recent functional data has shown that OSN activation leads to local Keystone LN depolarisation, suggesting that neurites within glomeruli can act independently^49^. Second, we found that Broad LNs inputs onto Keystone LNs are not homogenous (Fig. S13a-b) and scale with OSN inputs onto Keystones LNs in each glomerulus (Fig. 4d and Fig. S13d). This suggests a strongly conserved and previously undescribed computation for olfactory processing, whereby Broad LNs implement divisive normalisation of Keystone LNs responses as a function of total OSN drive per glomerulus. This structured feedback between the two components of the global inhibitory network could scale lateral inhibition over a range of odour concentrations.

### Evolution of multisensory integration in a global inhibitory network

Unlike the conservation of general connectivity features in Broad LNs (Fig. 3c), we found that the types of inputs Keystone LNs receive have changed, while their output structure has remained conserved (Fig. 4e-f and Fig. S13c). In *D. erecta*, sensory input to these interneurons is dominated by OSNs, while in *D. melanogaster* they receive a larger contribution from non-olfactory sensory neurons from the suboesophageal zone (SEZ), predominantly gustatory, thermo- and hygro-sensory neurons (Fig. 4e-f and Fig. S11 and S13). This evolutionary switch in the multi-sensory integration of Keystone LNs suggest that, in *D. erecta,* Keystone LNs are more driven by ecologically relevant olfactory input (Fig. 4e-i), presumably leading to keystone-dependent processing in the presence of pandan.

To further explore this pattern, we analysed the connectivity of Keystone LNs with OSNs, taking into account the relative anatomical position of each glomerulus. This is functionally important because OSN activation generates Keystone LN neurites depolarisation over a few nearby glomeruli^49^. Despite the larger input contribution from OSNs in *D. erecta*, the relative connectivity pattern to Keystones LNs is overall conserved with a few cross-species differences (Fig. 4g). In both species there is a cluster of antero-ventral glomeruli strongly connected to Keystone LNs likely strongly influencing their activation (Fig. 4g). In addition, *D. erecta* Keystone LNs are more strongly connected to a posterior cluster of glomeruli tuned to leafy odours, and in *D. melanogaster* the connectivity to yeasty/leafy glomeruli is stronger (Fig. 4g-h). The proportion of outputs to OSNs is more similar between the two species, with common targets that include the above-mentioned common input cluster, and a few – mostly leafy glomeruli – more strongly targeted in *D. erecta* (Fig. 4g). These connectivity patterns suggest that Keystone LNs are activated in the presence of host cues and evolution has tweaked their tuning, biasing it towards each species niche – leafy olfactory in *D. erecta* and mixed olfactory and gustatory for *D. melanogaster*. Their activation, in turn, leads to presynaptic inhibition of select OSNs, a computation that produces lateral inhibition potentially enhancing contrast among key odour channels.

Moreover, Keystone LN activity is also regulated by Picky LN 0, which inhibits them in a glomerular-selective fashion, more so in *D. melanogaster* than in *D. erecta*. This contributes further to the heterogeneous inhibition that Keystone LNs exert on OSNs. For example, 82a OSNs are under slightly stronger Keystone LN inhibition in *D. erecta*, a difference that is amplified through the *D. melanogaster*-specific inhibition of Keystone LNs in this glomerulus (Fig. 4j and S13e). Together with the results above, this implies that 82a is under stronger inhibition from Broad LNs in *D. melanogaster*, and from Keystone LNs in *D. erecta* (Fig. 4j and Fig. S13b and e).

### Evolution of modality processing in a potential ripeness circuit

In addition to the two global inhibitory circuits described above, the larval AL has a network of selective inhibitory interneurons whose function remains largely unknown. These circuits are composed of three neuronal classes: Choosy LNs, Ventral LNs and Picky LNs.

There are two Choosy LNs per AL with very similar connectivity to each other (Fig. 5a-c). In both species they get half of their input from sensory neurons, mostly OSNs, followed by the suboesophageal zone (SEZ) sensory afferents (mainly taste and mechanosensory). However, the non-sensory input contributions to Choosy LNs have evolved. In *D. melanogaster* the other half of the input comes from processed olfactory information, but in *D. erecta* they receive a larger fraction from processed SEZ information, indicating that these neurons integrate a more complex multimodal stimuli, including olfactory, taste, thermal and mechanosensory(Fig. 5d and Fig. S14). This pattern is mirrored in their outputs, where *D.erecta* Choosy LNs talk more to the SEZ, while in *D. melanogaster* they primarily target uPNs and mPNs in the AL (Fig. 5d and Fig. S14). Therefore, Choosy LNs seem to have evolved a stronger role as integrators of multimodal information in the specialist, in addition to their role in olfaction as providers of postsynaptic inhibition to projection neurons.

**Figure 5.**
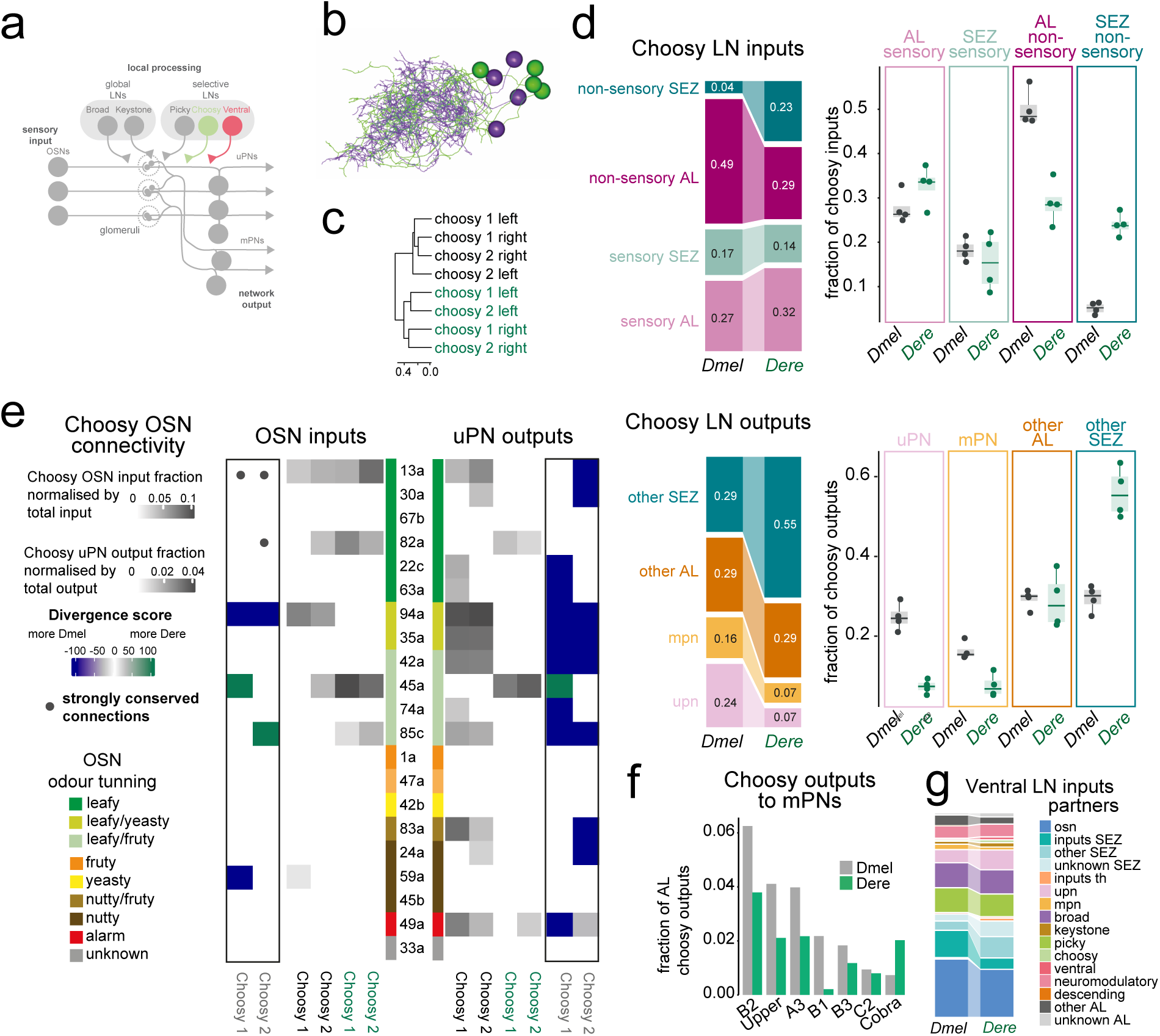
Evolution of modality processing in a potential ripeness circuit. **a,** Schematic of the olfactory system of the Drosophilid larva indicating each of the main neuronal classes, highlighting Choosy LNs (light green) and Ventral LNs (dark pink). **b,** Morphology of Choosy LNs in D. melanogaster (magenta) and D. erecta (green), registered on the same AL space. **c,** Clustering of Choosy LNs of both species based on total connectivity (homologous neurons only) after filtering for connections consisting of 2 or more synapses. **d,** Fraction of total Choosy LN inputs (top) and outputs (bottom) to/from the indicated neuronal classes for each species. To the right, boxplots display the same data as the barplot but show the variability across neurons, indicating robust differences in the non-sensory AL and SEZ input, as well as PN and SEZ outputs. **e,** Fraction of total Choosy LN inputs (right) and outputs (left) from OSNs (right). Only connections that constitute at least 1% of total input connectivity in both hemispheres of a species are shown. To the side, divergence confidence score (see methods) of the differences in OSN input/outputs to/from Choosy LNs between the two species. Indicated as dots are strongly conserved connections (see methods). **f,** Fraction of Choosy LN outputs within the AL onto each of the mPNs. Only connections consisting of 2 or more synapses were used for this plot. **g,** Fraction of total Ventral LN inputs from the indicated neuronal classes for each species.

To gain insights into the putative – still unknown – function of Choosy LNs, we looked at their main input partners, OSNs. In both species, these interneurons get input almost exclusively from OSNs with leafy descriptors (Fig. 5e and Fig. S14a), with 82a and 45a – the high and low sensitivity OSNs for geranyl acetate, a pandan volatile – more connected in *D. erecta*. Interestingly, 13a is the only OSN strongly synapsing with all Choosy LNs in both species. This OSN had been described to respond mostly to leafy odours, but some of its major ligands have also been identified as products of the degradation of fruit pigments by bacteria^48^ and as products of yeast fermentation (Table S2). It is possible that Choosy LNs integrate olfactory channels and modalities to compute fruit “ripeness”; 13a would indicate fruit degradation by microorganisms, while the other leafy “green” channels could inform ripeness progression – in particular for *D. erecta* through the levels of geranyl acetate in pandan -, taste inputs carry information on bitter/sweet qualities of the substrate which decrease/increase as fruits ripen, while mechanosensory information informs fruit hardness. This suggest a previously unappreciated possibility, that some olfactory LNs might compute stimulus features that are ecologically relevant, such as “ripeness”, analogous to, what has been suggested for some amacrine cells in the visual system^50,51^. Under such scenario, Choosy LN input connectivity might have evolved to best represent the ripeness features of each species fruit hosts.

This ripeness feature representation in Choosy LNs could be used to shape the response of key projection neurons, thus leading to appropriate behaviours. As major providers of inhibition to uPNs and mPNs Choosy LNs are well positioned to carry out this function (Fig. 5d). Choosy LNs provide postsynaptic inhibition to half of the uPNs in *D. melanogaster* and the two geranyl acetate sensors in *D. erecta* (Fig. 5e and Fig. S14b). This can mould the responses of these uPNs to the derivative of OSN firing ^42,52^, thus adjusting their dynamic range as a function of fruit host ripeness. At the same time, Choosy LNs shape the responses of a subset of mPNs (Fig. 5f). Although little is known about the function of these projection neurons, Cobra mPN – which is targeted more strongly by Choosy LNs in *D. erecta* (Fig. 5f) – has been described to mediate innate aversion to geranyl acetate^53^, raising the possibility that adequate pandan ripeness levels could, through the activation of Choosy LNs, counteract this aversion.

We next looked at another component of the selective inhibitory network, the Ventral LNs. These interneurons are perhaps the most enigmatic within the AL, as their function, even their neurotransmitter remains unknown^12^. Their general connectivity is very similar across species (Fig. 5g), likely indicating a conserved role. Most of their OSN input comes from 13a (Fig. S14c), hinting that they might also be tuned to fruit ripeness.

### A switchboard for olfactory evolution

A characteristic feature of Picky LNs, the last component of the selective inhibitory circuit, is that each of its five types has very distinctive morphology and connectivity. These are sufficiently conserved to drive clustering of one-to-one homologs across species (Fig. 6a-b).

**Figure 6.**
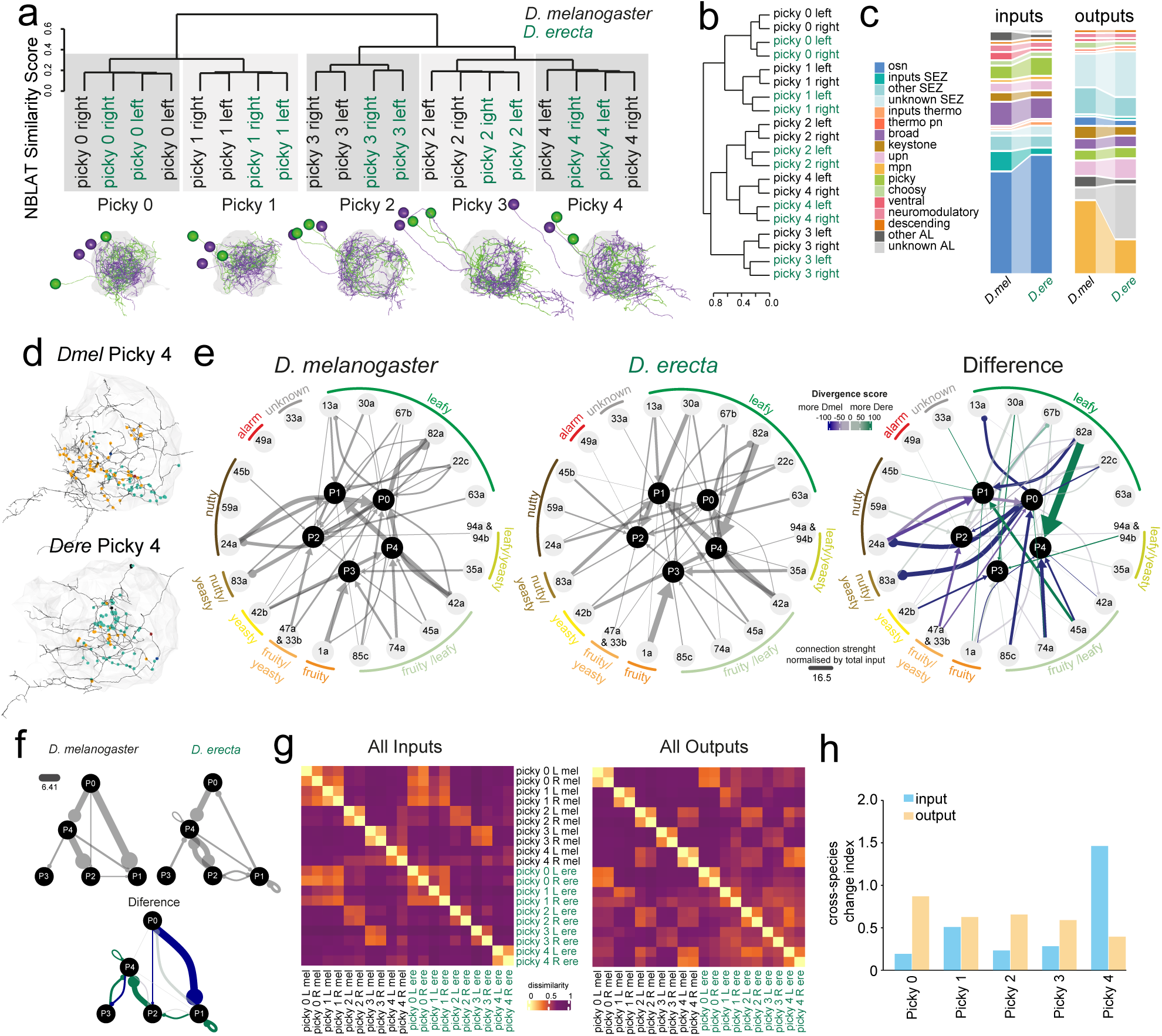
Picky LNs: A switchboard for olfactory evolution. **a,** Clustering of Picky LNs of both species based on morphological similarity (NBLAST scores) identifies clear one to one homologs for each of the Picky LNs (top). Their morphology is shown at the bottom: *D. melanogaster* neurons in magenta and *D. erecta* in green. **b,** Clustering of Picky LNs based on total connectivity (homologous neurons only) after filtering for connections consisting of 2 or more synapses in both hemispheres of a species. **c,** Fraction of total Picky LN inputs from the indicated neuronal classes for each species. **d,** Morphology and input synapses (dots coloured by ecology, green = leafy descriptors, yellow = yeasty and fruity) of the left Picky 4 LN of *D. melanogaster* (left) and *D. erecta* (right). **e,** Network plots of the connectivity between each Picky LN and each OSN in *D. melanogaster* and *D. erecta*. Connections have been filtered to include only those consisting of 2 or more synapses in both hemispheres of a species. The thickness of the arrows represents the fraction of total input onto each Picky LN and are an average of the left and right homologs of each species (left). The arrows of the plot in the right represent the difference in connectivity between *D. melanogaster* and *D. erecta*. The colour of the arrows shows the confidence score, green is more connected in *D. erecta* and magenta more connected in *D. melanogaster*. **f,** Network plots of the connectivity between each of the Picky LNs. Connections have been filtered to include only those consisting of 2 or more synapses in both hemispheres of a species. The thickness of the arrows represents the fraction of total input onto each Picky LN and are an average of the left and right homologs of each species. To the left, the plot shows the difference in connectivity between *D. melanogaster* and *D. erecta*. The colour of the arrows shows the divergence confidence score, green is more connected in *D. erecta* and magenta more connected in *D. melanogaster*. To aid visualisation, as shown in the figure legend, differences under 30% are not represented in grey but in very light shades of green or magenta to indicate cross-species differences with weak confidence support. **g,** Dissimilarity matrix of Picky LNs based on the Euclidean distances of the vectors formed by the inputs(right) or outputs (left) onto/from Picky LNs to/from all neurons for which homologs were identified across both species. **h,** Cross-species change index, which measures how much connectivity has changed across species when compared to the intra-species variability (zero indicating no change, see Methods) of the total inputs (blue) and total outputs (orange) for each Picky LN.

The general input structure to Picky LNs is well conserved, with some subtle type and species-specific differences (Fig. 6c and Fig. S15a-b). In contrast, there has been considerable divergence in the pattern of OSN inputs, with complete gain/losses of connections for every Picky LN (Fig. 6d-e and Fig. S15c). The most striking differences are in Picky LN 4, including strong and species-specific input in *D. erecta* from 82a OSNs – narrowly tuned to geranyl acetate, a pandan compound^37^, and stronger input from leafy OSN 67b, as well as input exclusively in *D. melanogaster*, but not in *D. erecta,* from OSNs with fruity descriptors 1a, 42a, 45a and 74a (Fig. 6e and Fig. S15c). We find a similar pattern for Picky LN 3, with connections in *D. erecta* but not *D. melanogaster* to leafy OSNs 30a, and leafy/yeasty 94a, while in *D. melanogaster* Picky LN 3 gets input from yeasty OSN 42b (Fig. 6e and Fig. S15c). Differences also exist for OSN input to the other Picky LN types (Fig. 6e and Fig. S15c and S16). Therefore, it appears that the stereotypically sparse OSN to Picky LN connectivity has acted as an evolutionary switchboard, rerouting information within this inhibitory LN network.

We have shown that in the AL, connectivity among interneurons is overall well conserved (Fig. 1g). However, surprisingly, the intra-picky LN connectivity has changed dramatically. In *D. melanogaster*, Picky LNs have been described to inhibit each other creating a Picky LN hierarchy^12^ with Picky LN 0 (P0) at the top. We found that *D. erecta* Picky LNs also inhibit each other but in a very different pattern to *D. melanogaster* and without a clear hierarchy (Fig. 6f). This likely generates different Picky LN dynamics between the two species and underscores Picky LN connectivity as an evolutionary hot-spot.

These data suggest that information processing by Picky LNs has substantially diverged across species. We, therefore, next examined whether their outputs have changed to the same extent. We computed the input and output dissimilarity matrices, and from these we calculated a cross-species change index, which measures how much connectivity has evolved across species when compared to intra-species variability (zero indicating no change, see Methods, Fig. 6g-h). This uncovered both, conserved Picky LN connectivity features, and patterns of evolutionary change. The dissimilarity matrices revealed that, in both species, the input connectivity of P0 and P1 is relatively similar to each other, while their outputs are more different (Fig. 6g and Fig. S16-S17), indicating that they integrate information from a similar set of neurons but then route this information to parallel channels. The opposite is true for P2 and P4, whose inputs are very different but share more common outputs (Fig. 6g). Importantly, the cross-species change index revealed that evolution has differentially impinged in the input and output connectivity of each Picky LN (Fig. 6h). This is particularly striking for P4, whose input connectivity has dramatically changed, but its output connectivity has diverged to a lesser extent, and the opposite is true for P0 (Fig. 6h).

It had previously been described that the main role of Picky LNs in olfactory processing could be to regulate the activity of mPNs^12^ (Fig. 6c). These projection neurons receive input from specific combinations of OSNs and Picky LNs, a motif that in *D. melanogaster* was hypothesised to confer them tuning to specific features of the odour space, in a way that could have been evolutionarily shaped^12^. We have shown that, indeed, the OSN to Picky LNs input seems to have evolved to match each species’ ecologies (Fig. 6e). In turn, the output of Picky LNs to mPNs seems to be overall better conserved but has changed in a Picky-specific way (Fig. S18). In the next section we investigate the evolution of the OSN-Picky-mPN motif in more detail.

### The multiglomerular output layer, a key component of the Picky LN circuitry, is also an evolutionary hub

Each of the 15 mPNs have clear homologs across species based on morphology clustering (Fig. S19). They are multimodal neurons receiving inputs both within and outside the AL, with a large variation in the input type for each mPN, a feature that is conserved across species (Fig. 7a-b, and Fig. S20a). We decided to classify them as Olfactory mPNs if they received at least 50% of their input connectivity from the AL, and these are the only mPNs we consider for subsequent analyses (Fig. 7b).

**Figure 7.**
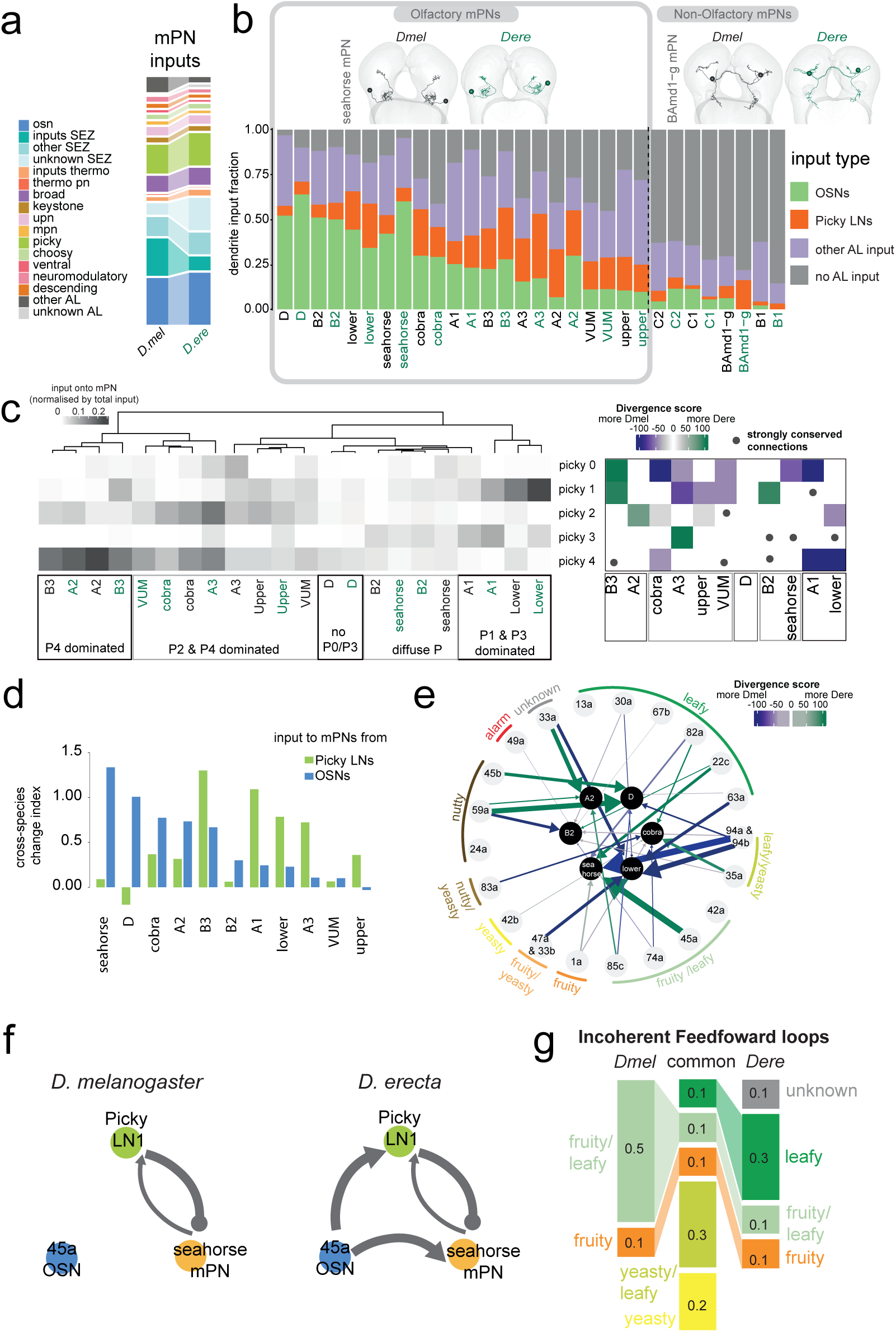
The multiglomerular output layer. **a,** Fraction of total dendritic input into mPNs from the neuronal classes indicated in the legend. **b,** Fraction of total dendritic input into mPNs from the following classes of neurons: OSNs (green), Picky LNs (orange), other AL neurons (purple), neurons outside of the AL (grey). We classified olfactory mPNs as those receive at least 50% of their inputs from AL neurons. On top, example morphology of an olfactory mPN (seahorse mPNs) and a non-olfactory mPN (BAmd1-g mPN). **c,** Input from Picky LNs onto olfactory mPNs identifies potential functional clusters. Only connections found in both hemispheres of a species are shown (left). The plot on the right shows the divergence confidence score (see methods) of the differences in Picky LN input to mPNs between the two species, dots indicate strongly conserved connections (see methods). **d,** Cross-species change index, which measures how much connectivity has changed across species when compared to the intra-species variability (zero indicating no change, see Methods) of the total inputs to mPNs from OSNs (blue) and Picky LNs (green). **e,** Network plot of the difference between *D. melanogaster* and *D. erecta* of the connectivity between the indicated mPNs and each OSN. The plot was made using only connections that constitute at least 2% of total input connectivity in both hemispheres of a species. The colour of the arrows shows the divergence confidence score, green is more connected in *D. erecta* and magenta more connected in *D. melanogaster*. To aid visualisation, as shown in the figure legend, differences under 30% are not represented in grey but in very light shades of green or magenta to indicate cross-species differences with weak confidence support. **f,** Network plot showing the connectivity between OSN 45a, Picky LN1 and seahorse mPN, as an example of an incoherent feedforward loop that is present in *D. erecta* but not in *D. melanogaster*. **g,** Proportion of the strongest feedforward loops grouped by those that are common to both species, or species-specific and classified according to the ecological significance of the chemicals the OSNs detect.

Given the suggested role of mPNs in controlling odour guided behaviours based on the input they receive from their main AL partners: Picky LNs and OSNs^53^, we focused on these two input channels. Despite some cross-species differences, mPNs can be grouped into five coarse groups based on Picky LN inputs (Fig. 7c). We have named them accordingly and speculate they might represent distinct functional groups essential for the computations carried out by the Picky LN – mPN circuit within the AL.

We next examined OSN inputs, each mPN samples from a stereotypical set of glomeruli^12^ (Fig. S20-S21). Our first question was whether this stereotyped OSN input connectivity might have evolved more than the Picky LN to mPN inputs. We computed the cross-species change index for both connectivity features and found that while, indeed, overall OSN inputs have evolved more than those from Picky LNs, the connectivity from the two neuronal classes has changed to different extents in different mPNs (Fig. 7d). For example, seahorse and D mPNs have dramatically evolved their OSN connectivity, while their Picky LN inputs have remained remarkably conserved. At the other end, A3 and B3 mPNs have undergone the opposite evolutionary trajectory, with many more changes to their Picky LNs than OSN inputs. This, again, underscores the “multiplexed” nature of evolutionary change in neuronal connectivity, whereby within the same dendritic field some input elements can diverge while other remain conserved.

To explore the evolution of OSN connectivity to mPNs, we plotted the network diagram of mPNs with at least 30% input contribution from OSNs (Fig. 7e, the connectivity matrix for all olfactory mPNs is in Fig. S20b). This highlighted some of the largest cross-species changes. For example, the strongest modification is a gain of connectivity of Or45a to seahorse mPN in *D. erecta* (Fig. 7e-f). This olfactory receptor senses the same pandan odour as Or82a (geranyl acetate) albeit with lower sensitivity^53,54^. Interestingly, Picky LN 1 also gained novel connectivity to Or45a in *D. erecta* (Fig. 6e and 7f). Furthermore, Picky LN 1 is connected to seahorse mPN (Fig. 7c and 7f). Therefore, these three neurons form, in *D. erecta,* an incoherent feedforward loop (IFLs), whereby Or45a activation leads to direct activation of seahorse mPN and its inhibition via Picky LN 1 (Fig. 7f). Intriguingly, another new IFL in *D. erecta* involves the high sensitivity geranyl acetate sensor OSN 82a, which is connected to cobra mPN (Fig. 7e) as well as Picky LN 4 (Fig. 6e) in *D erecta* but not in *D. melanogaster* – the loop is closed by the connection of Picky LN 4 to cobra mPN (Fig. 7c and Fig. S21). To examine these OSN-Picky LN-mPN IFLs in an unbiased way we selected the top 10 for each species (Table S3, see methods). We found that 40% of these are shared between the two species and have a strong representation of OSNs with yeasty descriptors (Fig. 7g) – matching the ecological relevance of yeast fermentation for both species. *D. melanogaster* specific IFLs have a stronger contribution from OSNs with fruity descriptors, while those exclusive to *D. erecta* are enriched in OSNs with leafy descriptors – including the two IFLs involving 82a and 45a described above (Fig. 7f-g and Table S3). These patterns reflect the evolution of the combinatorial OSN excitatory and Picky LN inhibitory input onto mPNs. A motif that shapes mPN responses, likely to represent key environmental features, leading to the activation of species’ appropriate odour-guided behavioural programmes.

## Discussion

To understand how neural circuits evolve at the cellular and synaptic levels, we need to compare the brains of animals with divergent behaviours at high resolution. Furthermore, key questions in evolutionary neuroscience – such as the evolvability of different elements in the circuit – are best addressed by examining complete circuits in an unbiased way. Here, we have leveraged the small size, yet complex organisation, of the olfactory circuit of the Drosophilid larvae to carry out, to our knowledge, the first cross-species comparative connectomics of a glomerularly organised olfactory circuit. Importantly, this glomerular organisation facilitates the interpretation of function out of structure, making them an ideal system for connectomics. By comparing, at synaptic resolution, the circuits of two closely related species with divergent ecologies and odour-guided behaviours, we have uncovered evolutionary patterns that might be generalisable to other networks (Fig. 8).

**Figure 8.**
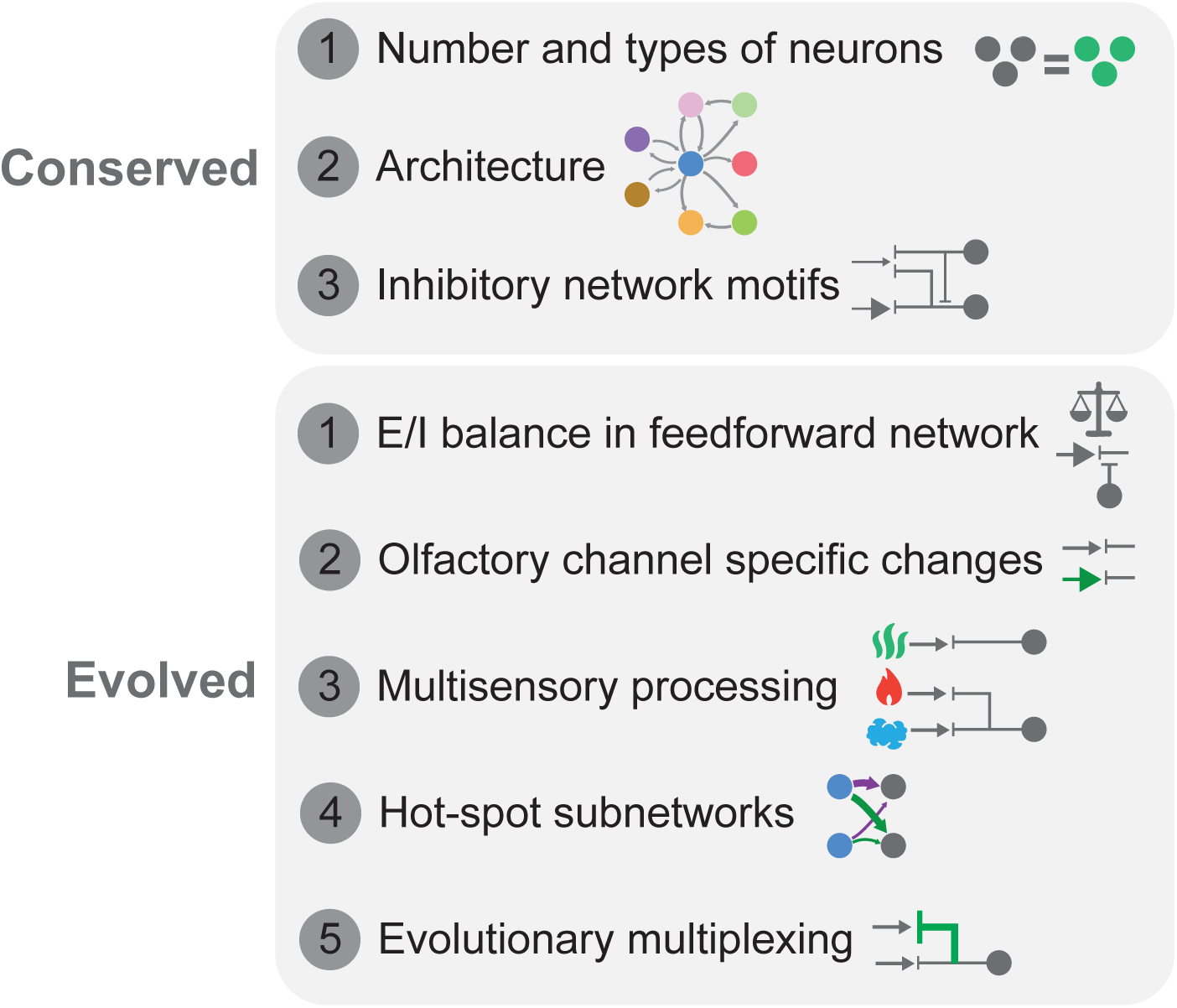
Patterns of evolution in an olfactory network. Key conserved and divergent evolved features. Circles represent neurons and lines their connectivity.

### Conserved features

Comparing circuits across species enables us to identify not only the changes, but also the conserved features that define a putative circuit blueprint. We found that both species display the same number and types of neurons, in contrast with the evolutionary divergence of cell numbers observed between distantly related nematode species^11^. This, together with previous work^3,4,7,21–23^, suggests that neuronal connectivity may be more evolvable than cellular composition.

We also found that the general circuit blueprint as defined by the connectivity among neuronal classes is well conserved. However, when looking at finer resolution, into individually identifiable neuronal elements, we found that while the interneuron-to-interneuron connectivity is conserved, the way in which each of the olfactory channels “plug” into this network has evolved to various degrees. This profiles a mode of evolution by which the circuit motifs performing core computational transformations required to process olfactory information remain conserved, but their synaptic weights and sensory channel specificity are shaped by species-specific environmental needs.

### Evolved features

A key difference between the olfactory system of the generalist versus the specialist species is a change in the excitatory-to-inhibitory ratio in the feedforward circuit, most prominently onto the dendrites of its main output channel, the uPNs. We speculate that this might be an adaptation to each species’ lifestyles. Lateral inhibition has been shown to decorrelate uPN responses^42^, increasing odour discriminability^17,42^, but it also introduces noise ^40,41^. Therefore, the observed shift would favour stimuli discriminability in the generalist – which needs to navigate varied substrates –, and sensitivity in the specialist (by reducing noise into the system) – for which accurately sensing the right ripening stage of a single fruit might be crucial. This finding contrasts with the evolutionary trajectory found when comparing subcircuits in the cortex of human and mice, whose projection neurons – the pyramidal cells – maintain a strict E/I ratio but where the interneuron-to-interneuron connections have evolved substantially^55^. The origin of this difference is unclear, given that the exact identity of the inhibitory neurons undergoing the change remains unknown, and future work on the identity and function of different types of cortical neurons might help resolve this question. This highlights the need to examine circuits in different brain regions and across a wide variety of species and evolutionary distances, as merging the knowledge across these studies can inform future work and help us gain a global understanding of how neural circuits evolve.

Another important finding is that not all elements of the system have evolved to the same extent. Generally, the interneuron-to-interneuron connectivity is well conserved while the sensory input that LNs receive has evolved to a greater extent. However, even this is variable from neuron to neuron. For example, we find that a single interneuron, Picky 4 LN, stands out as a hot-spot, having changed its input connectivity the most. Furthermore, within a single neuron, synapses with some neuronal types might remain conserved while those with other neuronal populations might change dramatically, evidencing synaptic evolutionary multiplexing, reminiscent of what has been reported in the evolution of direction selective circuits in the retinas of mice and rabbits^5^. In the antennal lobe, this is best exemplified by the inputs to mPNs, where each mPN has independently evolved its Picky LN and OSN inputs to different extents.

### Outlook and limitations

Examining full circuits at synaptic resolution is challenging given the limitations of data acquisition, both in terms of imaging and circuit reconstruction. Therefore, we have compared a single individual of each species, meaning that potentially some of the differences identified could be cross-individual rather than cross-species. We have aimed at minimising this possibility by leveraging bilateral symmetry (the lack of crossover of OSNs in the larva), and previous connectomics work, to inform a stringent framework for the identification of cross-species differences (see Methods). We attach to every connection two values: the magnitude of change and a confidence score of divergence/conservation. Changes represented by a large difference in magnitude across species and a strong divergence confidence value are most likely to be true cross-species differences, and we have focused on these throughout the manuscript. In the future, acquisition of multiple connectomes per species – aided by recent advances in automatic neuronal reconstruction^56^ – will enable to fine tune the confidence values identified here. Finally, arguably one of the most important outcomes of connectomics is the generation of specific and testable hypotheses. In this regard, our work – thanks to the tractability of Drosophilids – opens exciting future opportunities to experimentally investigate the functional implications of the identified neural circuit changes and their contribution to behavioural evolution.

## Methods

### Electron microscopy sample preparation

The central nervous system of an 8 to12 hour old *D. erecta* larva was dissected in 4°C cold Baines solution and immediately processed. The fixation protocol was adapted from^57^ with the following modifications. All steps were done with 0.1 M sodium cacodylate buffer (pH 7.4), unless stated otherwise. The sample was fixed in primary fixative of 2.5% glutaraldehyde, 2.5% formaldehyde on ice for 1 h. After washing, the sample was transferred to 0.5% OsO_4_ for 40min on ice in the dark. After removing the OsO_4_ solution, the sample was immediately incubated with 0.8% K_4_[Fe(CN)_6_ (potassium ferrocyanide) for 2 h on ice. For the rest of the protocol, sodium cacodylate buffer was exchanged with water, and the sample was thoroughly washed after each step. The washed sample was incubated with filtered 1% TCH (thiocarbohydrazide) for 15 min at room temperature in the dark, followed by 2% OsO_4_ on ice for 30 min, and 0.5% uranyl acetate for 20 min on ice. Afterwards, the sample was incubated in a preheated (60°C) Walton lead solution^58^ for 30 min at 55°C and then left at room temperature for 2 h. Dehydration with acetone follows the progressive lowering of temperature (PLT) protocol as published^57^, but without the addition of heavy metals throughout the procedure. For the last PLT step, 0.2% uranyl acetate in acetone medium was used. After the PLT steps, the sample was infiltrated with 100% acetone on ice, embedded in Durcupan epoxy resin, and polymerized at 60°C for 48 h.

### FIB-SEM sample preparation

The Durcupan embedded first instar Larva of *D. erecta* sample was prepared for the high resolution large volume imaging of the two brain lobes and partial central Ventral Nerve Cord (VNC). This sample was initially mounted to the top of a 1 mm copper post which was in contact with the metal-stained sample for better charge dissipation. A small vertical sample post was trimmed to a small block containing the Region of Interest (ROI) with a width of 145 µm perpendicular to the ion beam, and a depth of 155 µm in the direction of the ion beam, as previously described^59^. The trimming was guided by X-ray tomography data obtained by a Zeiss Versa XRM-510 and optical inspection under an ultramicrotome. Thin layers of conductive material of 10-nm gold followed by 100-nm carbon were coated on the trimmed samples using a Gatan 682 Precision Etching and Coating System. The coating parameters were 6 keV, 200 nA on both argon gas plasma sources, 10 rpm sample rotation with 45-degree tilt.

### FIB-SEM image acquisition and registration

This FIB-SEM prepared sample was imaged by a customized Zeiss Merlin FIB-SEM system previously described^60,61^. The block face was imaged by a 3 nA electron beam with 1.2 keV landing energy at 10 MHz scanning rate. The x-y pixel resolution was set at 8 nm. A subsequently applied focused Ga^+^ beam of 16 nA at 30 keV strafed across the top surface and ablated away 8 nm of the surface. The newly exposed surface was then imaged again. The ablation – imaging cycle continued about once every minute for 12 days to complete FIB-SEM imaging the two brain lobes and partial VNC. The acquired image stack formed a raw imaged volume, followed by post processing of image registration and alignment using a Scale Invariant Feature Transform (SIFT) based algorithm. The aligned stack resulted in a final isotropic volume of 135 x 145 x 120 µm^3^ at 8-nm voxel resolution. The volume was then contrast-adjusted and split into tiles at different zoom levels for import into CATMAID^62^.

### Neuronal reconstruction

Neurons and synapses in the *D. erecta* brain were manually reconstructed using CATMAID^62^ as previously described^31,32^. Briefly, the antennal lobe was identified based on stereotyped landmarks and olfactory sensory neurons (OSNs) identified by their axons leaving the brain via the antennal nerve. Reconstructions started at OSNs tracing their pre- and post-synaptic partners. Reconstructed neurons were subsequently proof-read using the “review” widget of CATMAID.

These reconstructions were compared to neurons and synapses reconstructed in CATMAID using the same methods on an existing electron microscopy volume of a *D. melanogaster* first instar larva^12,31,32^. A possible limitation of this work is that the *D. melanogaster* volume was acquired employing different EM technology (transmitted electron microscopy, rather than eFIB-SEM) and reconstructed by a different team of people, which could introduce biases. However, the stringent analysis pipeline based on bilateral symmetry that we have implemented aims at minimising technical errors – which are unlikely to be bilaterally consistent. Furthermore, our work has better resolution than published work based on automatic reconstructions, as manual reconstructions are more precise, recovering a much higher proportion of synapses and thin branches. Finally, we proof-read and corrected the OSN pre-synapses of the *D. melanogaster* volume to ensure similar tracing across both datasets at this level.

Common to both volumes (the *D. melanogaster* and the *D. erecta*), and indeed a characteristic of all Drosophilid connectomes^12,31,33,35,63^, is the presence of traced neurites that can not be connected to any neuron. Following the consensus of previous literature^35^, we have classed these “fragments” as “unknown”, and further classify them into “unknown AL” or “unknown SEZ” based on their location.

### Identification of homologous neurons

We first identified homologous OSNs and uPNs across species by employing the relative position of each glomerulus in the AL. Next, we validated these assignments by examining the morphology of uPNs – similar morphology is a hallmark of homology^23^. Because uPNs are born sequentially from different neuroblasts the position of their cell bodies reflects this developmental origin, with neurons forming a tract in the direction of the neuroblast of origin with earlier born neurons having cell bodies closer to the neuropile and later born neurons closer to the outside of the brain. Homologous neurons are expected to have common developmental origins^23^ and thus similar cell body positioning. Next, we examined the anatomy of uPNs, in particular their axonal paths and arbourisations in the lateral horn and mushroom body. This was done iteratively by several observers. This gave a high confidence identification of homologs of all OSN-uPN pairs across species. We note that the naming of the glomeruli is according to the olfactory receptor expressed by each OSN in *D. melanogaster*. We find that *D. erecta* has the same complement of olfactory receptors and targeted experiments have confirmed for a subset of them – as expected – their expression in the same OSNs as in *D. melanogaster*. Next, based on general morphology and cell body position, we identified the five types of AL LNs, and separately examined the components of each type to assign one to one homologies.

Once neurons had been assigned homologies across species through the abovementioned methods, we verified these homologies employing NBLAST^64^, which scores neuronal morphological similarity. NBLAST requires the neurons being compared to be on the same space. Therefore, we registered the left and right hemispheres of the *D. melanogaster* and the *D. erecta* datasets onto a common space through rigid and non-rigid transformations using Deformetrica^65^. NBLAST scores were then clustered by Euclidean distances to generate the trees shown in the figures.

Some neurons could only be found in the AL connectome of either *D. melanogaster* or *D. erecta*. These are predominately neurons connected only very weakly to AL neurons and in most cases the connections are not bilaterally consistent. Therefore, in most cases, and most likely all cases, these are neurons that exist in both species but that in one or the other are not connected to neurons of AL. For example, Trito neurons exist in both species, but in *D. melanogaster* only receive SEZ input while in *D. erecta* they receive a small fraction of OSN inputs. To favour a conservative approach to the identification of species differences, for some analyses we have excluded these neurons, thus only including those neurons for which we could find homologs across both species. In each case, this is indicated in the figure legends.

### Comparing connectomes across species

To compare connectomes across species we first defined what constitutes each species connectome. Because OSNs do not cross-over the midline, the right and left ALs develop independently. Therefore, we used bilateral symmetry as an indication of true species characteristic connectivity and consider asymmetries as an indication of within-species variability. This approach is based on previous connectomics work. First, work on the *D. melanogaster* larvae showed that connections between two identified neurons that are represented by 2 or more synapses in each hemisphere of the same animal are true species-characteristic connections that can be found reliably between the same homologous identifiable neurons across individuals^32^. Furthermore, analysis of whole brain connectomes of the *D. melanogaster* larval and adult revealed that connections that constitute more than 1% of either input or output connectivity are considered to be strong and conserved within a species^31,33,34^. Therefore, we use these two heuristics through the paper to describe species-characteristic connectivity. Specifically, we define species-characteristic connections as those between homologous pre- and post-synaptic partners that are observed to consist of 2 or more synapses in both hemispheres (e.g. a connection between two homologous neurons that is represented by a single synapse in one side is not considered a reliable connection, even if on the other hemisphere the connection consists of more than 2 synapses). We define strong connections using a similar criteria but with a higher threshold, considering strong species-characteristic connections as those that constitute in each hemisphere at least 1% of input or output connectivity between two connected pairs of homologous neurons. We indicate in the text and the figures which of these two thresholds is being applied. For bilateral neurons, such as Keystone LNs and some mPNs, we applied the same rule as for unilateral neurons, virtually flipping their arbours to compare left-right homologs connectivity (i.e. we avoid thresholding across the branches within the same hemisphere, and instead compare the arbour ipsilateral to cell body of each homolog with each other and the contralateral arbour to the contralateral arbour for each neuron).

To compare connectomes across species we have applied a stringent framework, favouring a conservative approach that robustly identifies cross-species changes, at the expense of overlooking potential subtle differences. This is one of the motivations behind employing two different synapse thresholds, for example, using the more stringent threshold of 1% might eliminate true synapses in one species, which might be represented only by 0.9% on one side, but not the other species, where both sides might be connected at 1%, if this is the only connectivity that is being compared, it could lead to the wrong conclusion that this is a connection present in one species but not the other, while the connection is truly present in both species, but at different strengths. Therefore, in an effort to robustly identify species differences, across the manuscript we have examined connectivity without any thresholding and with the two thresholds presented above. In addition, we attach to every species divergence a confidence score, calculated based on three parameters: 1) for any connection between two identified neurons, the number of synapses connecting them in any of the two hemispheres of one species can not overlap with that of the equivalent connectivity between homologous across any of the two hemispheres of the other species, if they do, the confidence score is drawn to zero, 2) the same criteria applies to their normalised connectivity – i.e, fraction of input or output connectivity that the connection represents. 3) we calculate for each connection the percentage of change in synapse fraction across species and, based on recent analysis of multiple connectomes of *D. melanogaster* adult brains, we assign a low confidence (grey colour) to differences below 30%^33^. Therefore, for each cross-species changes in connectivity we report the magnitude of the difference and its confidence. This means that sometimes we might find small differences, with high confidence, and conversely bigger differences with lower confidence, depending on how variable the connectivity is across hemispheres within a species. In addition, we identify robustly conserved connections as those that are represented by more than 2 synapses across both hemispheres of each species, and whereby both the raw number of synapses as well as its proportion of total synapses have at least 90% similarity, including those connections where the distributions of either raw synaptic count or proportion of synapses overlaps across species at least 80%.

### Community detection

Modularity analysis was performed using the Louvain method of modularity maximization^66^, by employing the “lovain_communities” function of the Communities subpackage of the NetworkX^67^ package for Python, with a standard resolution of 1. Briefly, given a network, methods of modularity maximization partition this network into modules or "communities" with the goal of maximizing the number of edges that fall within communities rather *than between communities*. To quantify how modular the partitions of the network are, these methods use the following modularity coefficient *Q*^68^.

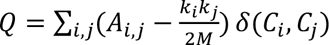

where *A*_*i*,*j*_ is the adjacency coefficient between nodes *i* and *j* (the total weight of the edges between *i* and *j*), *k*_*i*_ (*k*_*j*_) is the weighted degree of node *i* (*j*), ie. the total weight of all edges touching that node, *m* is the total weighted number of edges in the network 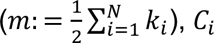 (resp. *C*_*j*_) is the community the node belongs to *i* (resp. *j*), *δ* is the Kronecker symbol (*δ*(*C*_*i*_, *C*_*j*_) = 0 if *C*_*i*_ ≠ *C*_*j*_ i.e., if *i* and *j* are in different communities, and *δ*(*C*_*i*_, *C*_*j*_) = 1 otherwise). The modularity coefficient *Q* is a measure of how many edges fall within communities (quantified by the sum of the *A*_*i*,*j*_ with *i* and *j* nodes in the same community) relative to a network in which the same number of edges are randomly distributed (with the same degrees of the nodes, quantified by 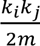). *Q* takes values between 0 (random configuration of edges) and 1, where high values of *Q* indicate partitions of the network that are strongly modular. It is important to empathise that because of how the modularity maximisation is run, it is likely that the resulting partitions of the network correspond to local, rather than absolute, modularity maxima. The mean modularity coefficient of the AL networks were: 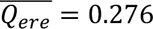 and 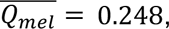 with very low variance across 1000 iterations, indicating that the network is not strongly modular^69^.

The Louvain algorithm proceeds in two phases. First, every node is in its own community. Nodes then change from belonging to their own community to the community of one of their neighbours if this increases the modularity coefficient *Q*. If not, the node stays in its own community. All nodes are considered sequentially according to this rule. When no further change of community membership at the node level leads to an increase in *Q*, the first phase ends. The second phase proceeds similarly, but on a new network in which the nodes are the communities found at the end of the first phase, thus grouping communities into communities. When no further move increases the modularity coefficient at this community level, the second phase ends. The combination of first and second phases is iterated until no further change leads to an increase in the modularity coefficient. We ran this algorithm independently for the *D. melanogaster* and *D. erecta* networks, including only AL core neurons. For each species, we ran the algorithm 1000 times independently to quantify the variation due to the order of nodes considered by the algorithm. For each species we selected the iteration with the highest repeatability and this is what is what we called the “reference” configuration, shown in Figure 1. This configuration had eight communities in each species, but of these six communities are common to both species (dashed lines in Figure 1). We use nodes’ trasparency to visualise the robustness of these configurations. More precisely, the transparency of each node is the proportion of iterations (out of all 1000 iterations) where the focal node is assigned to the same community as in the ‘reference’ configuration. For example, neurons that were assigned to the same community in all 1000 repetitions are coloured in solid colour, while those that only appeared in the ‘reference’ configuration 500 times would have a 50% transparency appearance.

### Glomerular meshes

Glomerular meshes were generated by taking the location of OSN inputs onto AL neurons and calculating a gaussian kernel density per OSN identity using the skimage script within the scikit-image python package^70^. Additionally, we generated a bounding AL mesh using olfactory inputs and outputs of Broad LNs and an alphashape fitted to the resultant points. We used these to then generate the final glomerular meshes with the ‘AL glomeruli meshes’ script from ^35^.

### Properties of Antennal lobe neurons

The segregation index was calculated using the R package catnat v0.1. Briefly, catnat makes use of the synapse flow centrality algorithm^63^ to assign dendrite and axon identity to sections of the neuronal arbour. Subsequently, it calculates a segregation index based on the distribution of pre- and post-synaptic sites in each compartment.

Local reaching centrality (LCR) and out-degree were calculated as presented in^34^. Briefly, LCR of a neuron (node) is a measure of the proportion of other nodes in the network can be reached via outgoing connections. Therefore, a high LCR is an indication that the activity of a neuron can – theoretically – propagate through a large part of the network. It was calculated using NetworkX v2.5^67^, which takes directed weighted graphs and takes into account the weight of each edge (number of synaptic connections). Out-degree is the weighted count of all outgoing connections from a neuron. Unlike LRC it measures the influence of a neuron over its immediate downstream synaptic partners.

Synaptic scores were calculated as presented in^35^. Briefly, for each morphological type we constructed a matrix where each row was a neuron and each column a glomerulus, indicating the number of input or output synapses each neuron gets/gives per glomerulus (for the input and output synapse scores respectively). For each neuron (column) the rows were normalised between 0 and 1 and sorted in descending order. From here the sum of the ranked glomeruli was calculated within each neuronal class.

### Excitatory vs inhibitory balance analysis

We calculated the bilateral averages of the number of excitatory and inhibitory input synapses for OSNs and uPNs, and calculated the ratio: excitatory divided by inhibitory. From here, we compared the number of synapses within each neuron-pair across the two species, normalised to *D. melanogaster* (=1), such that *D. erecta* data shows the corresponding fold-change. Stats are based on a 2-tailed Wilcoxon Rank test. In addition, we also used origin-restricted linear regression to quantify the relationship between the number of excitatory and inhibitory synapses per neurons in a class. These analyses were done using custom written scripts in Igor-Pro 9 (Wavemetrics).

### Cross-species change index and dissimilarity matrices

To produce the dissimilarity matrices, we generated for each neuron a vector of either its input or output connectivity normalised by total inputs or outputs from homologous identified neurons. Using these normalised vectors, we calculated the Euclidean distance between each neuron and plotted it as a dissimilarity matrix using R. The cross-species change index was then calculated as follows. We estimated intra-species dissimilarity for each neuron by averaging the dissimilarity index of the left and right homologs of each species, i.e. intra-species dissimilarity = μ ((dissimilarity left vs right in Dmel), (dissimiliary left vs right in Dere)). Then we estimated inter-species dissimilarity by averaging the dissimilarity between all the possible combinations of homologs across species, i.e. inter-species similarity = μ ((dissimilarity of right Dmel vs right Dere), (dissimilarity of right Dmel vs left Dere), (dissimilarity of left Dmel vs right Dere), (dissimilarity of left Dmel vs left Dere)). The cross-species change index was then calculated as, index = ((inter-species dissimilarity) – (intra-species dissimilarity)) / (intra-species dissimilarity).

### Connectivity per glomerulus in Keystones

To calculate the input and output that the neurites of Keystone LNs make with Broad LNs and Picky 0 LN per glomeruli, we first generated meshes for each glomerulus in the four ALs of our datasets, and then extracted the synapses of each Keystone LN branch from within the corresponding glomeruli mesh.

### Behavioural assays

Clear 10 cm square petri dishes filled with 1% agarose in distilled water were used as experimental arenas. At opposing sides of the plate, 1 cm from the edge, two stripes of parafilm were placed, and on top 10 droplets of 10ul each of either grape juice or pandan extract were pipetted onto each side. For each experiment, 10-15 newly hatched (0-2 hours after egg laying) larvae were place on the midline of the plate and left to crawl for 30 minutes and final larval positions recorded. The preference index was calculated as A-B/A+B, A and B being the number of larvae on each side. Larvae that did not move from the midline were excluded from the analysis.

### Visualisations

The *D. erecta* brain mesh was generated by segmenting a subset of frames using 3DSlicer^71^ and using the interpolation function. This generated a NRRD file that was used to compute a mesh object by employing the Marching Cubes algorithm via the skin image and trimesh modules in Python^70^. The mesh was exported in STL format and subsequently treated in MeshLab 2021.05^72^ by using its Uniform Mesh Resampling tool. Finally, this resulting mesh was solid-fied using the Make Solid tool in MeshMixer. Neuronal skeletons were either exported from CATMAID and plotted using R scripts, or imported into Blender.

## Data availability

All analyses can be reproduced by using R pack scripts, together with R-markdown documentation and the full dataset found in: https://github.com/PrietoGodinoLab/Comparative.Connectomics.Drosophila.Larvae, by using R (v 4.4.0) and the following packages: ggplot2^73^ (v3.5.1), igraph^74,75^ (v2.1.2), dplyr^76^ (v1.1.4), magrittr^77,78^ (v2.0.3), stringr^79,80^ (v1.5.1), tidyr^81^ (v1.3.1), svDialogs^82^ (v1.1.0), reshape2^83^ (v1.4.4), tcltk^84^ (v4.4.0), ComplexHeatmap^85^ (v2.20.0), grid^86^ (v4.4.0), circlize^87^ (v0.4.16), Morpho^88^ (v2.12), and natverse^89^ (v0.2.4).

## Acknowledgements

We thank Alexander Shakeel Bates for assistance in early stages of the project; Tom Baden for advice on various analyses, discussions and comments on the manuscript; Michael Winding and Antonio Torres Mendez for comments on the manuscript; The Scientific Computing Science Technology platform at The Crick for assistance with Catmaid; R.J.V.R. was supported by a Boehringer Ingelheim Fonds Ph.D. fellowship. C.G. is supported by a EMBO Postdoctoral Fellowship (ALTF1114-2024)) and HFSP Postdoctoral Fellowship (LT0036/2025-L). S.P., Z.L., C.S.X. and H.F.H. were supported by Janelia Research Campus of the Howard Hughes Medical Institute. Work in the L.L.P.-G. laboratory is supported by a European Research Council Starting Investigator Grant (802531), an Allen Distinguished Investigator Award, a Human Frontiers Science Grant (RGY0052/2022), a Vallee Scholar Award and a Chan Zuckerberg Collaborative Pairs Project (CP-2-1-Prieto-Godino). Work in the L.L.P.-G. and J.D. laboratories is also supported by the Francis Crick Institute, which receives its core funding from Cancer Research UK (CC2067 and CC2240), the UK Medical Research Council (CC2067 and CC2240) and the Wellcome Trust (CC2067 and CC2240).

## Author contributions

L..L.P.-G conceived the project; N.R. dissected the *D. erecta* brain, optimised and carried out the staining and embedding; Z.L. assisted in the *D. erecta* brain staining; S.P. performed FIB-SEM sample preparation; S.P. and C.S.X. performed FIB-SEM imaging of the *D. erecta* brain volume; H.F.H. provided funding support for FIB-SEM operations; M.Z. and A.C. coordinated the *D. erecta* brain preparation and imaging and provided funding; A.C. aligned and prepared the *D. erecta* EM volume for analysis; R.J.V.R. set-up the analysis pipeline, performed 90% of the tracing, revised the OSN synapses in the *D. melanogaster* volume and identified homologous neurons across species and hemispheres; S.D. performed 6% of the tracing; R.J.V.R, S.D, C.G and L.L.P.-G analysed the data; H.G. performed the behavioural experiment shown in Fig. 1 and generated the assignments of odour scenes to olfactory sensory neurons; L.D. performed the community detection analysis under the supervision of J.D; L.L.P.-G wrote the paper with input from all authors.

## Extended Data

**Figure S1. uPN homologies across species.** Morphology of each of the uPNs for both species.

**Figure S2. LN general connectivity features. a,** NBLAST of LNs validates homologies across species (right) and clustering of LNs based on total connectivity (homologous neurons only) recovers similar grouping as morphological clustering (left), indicating conserved general connectivity features. **b,** Input (left) and output (right) synaptic scores^35^ (see Methods) for each of the LNs in both species. In these plots, the value in the X axis where the curves asymptote represents the number of glomeruli sampled, while the height in the Y axis is a function of how many glomeruli have been sampled and how evenly they are sampled. Solid line is the mean, shading represents standard error of the mean. **c,** Segregation index (see Methods) for each of the main AL neuronal classes in both species. Neurons with more clearly distinguishable input (dendritic) and output (axonal) elements have higher segregation indices. **d,** Schematic of the network analysis^34^ (see methods). Neurons with high out-degree (strong output connections) have strong effects on the network, neurons with high local reaching centrality (LRC) can reach many nodes in the network. **e,** Results from out-degree and LRC for each of the main neuronal classes within the AL in both species.

**Figure S3. Additional connectivity features of the uniglomerular circuit. a,** Percentage of uPN inputs from its cognate OSN in *D. melanogaster* (black) and *D. erecta* (green). Bars are the average between the two circles that represent left and right hemispheres of each species. Highlighted in blue are the three uPNs that display increased OSN to uPN connectivity in *D. erecta* regardless of the metric used (total synapse number, normalised by uPN input or by uPN cable length). **b,** Number of synapses onto each uPN from its cognate OSN, normalised by each uPN dendritic cable length in *D. melanogaster* (black) and *D. erecta* (green). Bars are the average between the two circles that represent left and right hemispheres of each species. Highlighted in blue are the three uPNs that display increased OSN to uPN connectivity in *D. erecta* regardless of the metric used (total synapse number, normalised by uPN input or by uPN cable length). **c,** Fraction of OSN inputs from each of the neuronal classes as indicated in the legend. **d,** Fold change of input from Broad LNs onto OSNs in *D. erecta* when compared to *D. melanogaster*. Each grey dot is an OSN, black represents the mean. The asterisks represent significance p = 0.2 x 10^-7^. Wilcoxon Rank test. **e,** Evolution of the ratio of excitation to inhibition onto OSNs across species. Fold change of excitatory inputs, inhibitory inputs and their ratio onto OSNs when compared to *D. melanogaster*. Each grey dot is an OSN, black represents the mean. The asterisks represent significance ns p = 0.79, *** p = 0.00035, * p = 0.01. Wilcoxon Rank test. **f,** Fold change of the excitatory to Broad LN input onto uPNs when compared to *D. melanogaster*. p = 0.2 x 10^-7^. Wilcoxon Rank test. **g,** Electron microscopy images of examples of autapses. **h,** Fraction of total input connectivity that OSNs receive from other OSNs. Only connections that consist of at least 2 synapses in both hemispheres of a species are shown.

**Figure S4. General connectivity of uPN dendrites. a,** Input onto uPN dendrites to the neurons indicated in the legend. The two left most bars are the mean, left and right side neurons are shown. **b,** Output from uPN dendrites to the neurons indicated in the legend. The two left most bars are the mean, left and right side neurons are shown.

**Figure S5. Input connectivity of Broad LNs.** Fraction of total Broad LN inputs from each of the indicated neurons. Only connections with homologous neurons identified across both species and that consist of at least 2 synapses in both hemispheres of a species are shown. Broad LNs are clustered based on the displayed connectivity.

**Figure S6. Output connectivity of Broad LNs.** Fraction of total outputs from Broad LNs to each of the indicated neurons. Only connections with homologous neurons identified across both species and that consist of at least 2 synapses in both hemispheres of a species are shown. Broad LNs are clustered based on the displayed connectivity.

**Figure S7. Inputs to Broad LNs from OSNs and uPNs. a,** Fraction of total Broad LN inputs (left) and outputs (right) to/from the indicated neuronal classes for each species. Left (L)/right (R) homologs are shown for each neuron and species. **b,** Fraction of total Broad LN inputs coming from each OSN (right). Only connections that consist of at least 2 synapses in both hemispheres of a species are shown. The plot on the left shows the divergence confidence score (see methods) of the differences in OSN input to Broad LNs between the two species, indicated as dots are those connections that are strongly conserved (see methods). **b,** Fraction of total Broad LN inputs coming from each uPN (right). Only connections that consist of at least 2 synapses in both hemispheres of a species are shown. The plot on the right shows the confidence score (see methods) of the differences in uPN input to Broad LNs between the two species, indicated as dots are those connections that are robustly conserved (see methods).

**Figure S8. Outputs from Broad LNs to OSNs and uPNs. a,** Dissimilarity plot of Broad LNs outputs (left) and the same when in-silico removing all Broad T to Broad T connectivity. This demonstrates that the strong output similarity among Broad Ts is mostly driven by intra-Broad T connectivity. **b,** Dissimilarity values of Broad inputs (blue) and outputs (yellow) when comparing across Broad types (within each species) in the real connectome, and when we in silico removed intra Broad T connectivity. This demonstrates that the strong output similarity among Broad Ts is mostly driven by intra-Broad T connectivity. **c,** Fraction of total outputs from each Broad LN onto each OSN (right). Only connections that consist of at least 2 synapses in both hemispheres of a species are shown. The plot on the left shows the divergence confidence score (see methods) of the differences in Broad LN output onto OSNs between the two species, indicated as dots are those connections that are strongly conserved (see methods). **d,** Fraction of total outputs from each Broad LN onto each uPN (right). Only connections that consist of at least 2 synapses in both hemispheres of a species are shown. The plot on the left shows the divergence confidence score (see methods) of the differences in Broad LN output onto uPNs between the two species, indicated as dots are those connections that are strongly conserved (see methods).

**Figure S9. Connectivity dissimilarity plots based on OSN input connectivity.** Euclidean dissimilarity based on total OSN input connectivity. All connections from OSNs to each LN were used to generate these similarity connectivity plots.

**Figure S10. Connectivity dissimilarity plots based on total output connectivity.** Euclidean dissimilarity based on total output connectivity. All output connections from each LN to neurons with homologs identified across both species were used to generate these similarity connectivity plots.

**Figure S11. Input connectivity of Keystone LNs.** Fraction of total inputs onto Keystone LNs from each of the indicated neurons. The two branches of each Keystone LN are shown separately. Only connections with homologous neurons identified across both species and that consist of at least 2 synapses in both hemispheres of a species are shown. Keystone LNs are clustered based on the displayed connectivity.

**Figure S12. Output connectivity of Keystone LNs.** Fraction of total outputs from Keystone LNs to each of the indicated neurons. Only connections with homologous neurons identified across both species and that consist of at least 2 synapses in both hemispheres of a species are shown. Keystone LNs are clustered based on the displayed connectivity.

**Figure S13. Additional connectivity features of Keystone LNs with Broad and Picky 0 LNs. a,** Fraction of total inputs coming from Broad LNs onto Keystone LNs at each glomerulus (right). Only connections that consist of at least 2 synapses in both hemispheres of a species are shown. Connectivity from the two Keystone LNs have been averaged. The glomerular map indicates the relative position of each glomerulus. The anterior (A) – Posterior (P) and Dorso (D) – Ventral (V) axis are indicated at the top left inset. Those with solid lines are located in the centre of the AL, those with small dashed lines are located anteriorly and those with larger dashed lines are located posteriorly. The colour of the outline depicts the ecological odour tuning of the OSN innervating each glomerulus (see methods). **b,** Fraction of Keystone LN total output to Broad LNs to each glomerulus. Only connections that consist of at least 2 synapses in both hemispheres of a species are shown. Connectivity from the two Keystone LNs have been averaged. **c,** Fraction of inputs (right) and outputs (left) onto Keystone LNs from each of the neuronal classes shown in the legend, each bar is a Keystone branch. **d,** Correlation between the OSN input and the Broad LN input received by Keystone LNs in each glomerulus. Each dot represents a single Keystone LN branch in each glomerulus. The correlation coefficient R^2^ = 0.16 and 0.25 for *D. melanogaster* and *D. erecta* respectively. **e,** Fraction of total inputs coming from Picky 0 LN onto Keystone LNs at each glomerulus (right). The two branches of each keystone are shown separately. All connections between Picky LN 0 and Keystone LN are shown. The plot on the left shows the confidence score (see methods) of the differences in Picky 0 LN input to Keystone LNs between the two species.

**Figure S14. Additional connectivity features of Choosy LNs and Ventral LNs. a,** Fraction of total inputs onto each Choosy LN from each OSN (right). Only connections that consist of at least 2 synapses in both hemispheres of a species are shown. The plot on the left shows the divergence confidence score (see methods) of the differences in OSN input to Choosy LN between the two species, dots indicate strongly conserved connections (see methods). **b,** Fraction of total outputs from each Choosy LN onto each uPN (right). Only connections with that consist of at least 2 synapses in both hemispheres of a species are shown. The plot on the left shows the divergence confidence score (see methods) of the differences in Choosy LN output onto uPNs between the two species, dots indicate strongly conserved connections (see methods). **c,** Fraction of total inputs onto each Ventral LN from each OSN (right). Only connections that consist of at least 2 synapses in both hemispheres of a species are shown. The plot on the left shows the confidence score (see methods) of the differences in Ventral LN OSN inputs between the two species. **d,** Fraction of total Choosy LN inputs (left) and outputs (right) from/to the indicated neuronal classes for each species. **e,** Fraction of total Ventral LN inputs (left) and outputs (right) from/to the indicated neuronal classes for each species.

**Figure S15. OSN input to Picky LNs. a,** Fraction of total Picky LN inputs from the indicated neuronal classes for each species. The two most left bars of each plot are the average, with total number of synapses on top. **b,** Fraction of total Picky LN outputs to the indicated neuronal classes for each species. The two most left bars of each plot are the average, with total number of synapses on top. **c,** Fraction of total inputs onto each Picky LN from each OSN (right). Only connections that consist of at least 2 synapses in both hemispheres of a species are shown. The plot on the left shows the divergence confidence score (see methods) of the differences in OSN input to Picky LN between the two species, dots indicate connections that are strongly conserved.

**Figure S16. Input connectivity of Picky LNs.** Fraction of total Picky LN inputs from each of the indicated neurons. Only connections with homologous neurons identified across both species and that consist of at least 2 synapses in both hemispheres of a species are shown. Picky LNs are clustered based on the displayed connectivity.

**Figure S17. Output connectivity of Picky LNs.** Fraction of total outputs from Picky LNs to each of the indicated neurons. Only connections with homologous neurons identified across both species and that consist of at least 2 synapses in both hemispheres of a species are shown. Picky LNs are clustered based on the displayed connectivity.

**Figure S18. Connectivity from Picky LNs to mPNs.** Fraction of mPN outputs from Picky LNs to each of the mPNs. The plot on the left shows the divergence confidence score (see methods) of the differences in Picky LN to mPN outputs between the two species, dots indicate strongly conserved connections (see methods).

**Figure S19. mPN homologies across species.** Clustering of mPNs of both species based on morphological similarity (NBLAST scores) identifies one to one homologs for each of the mPNs.

**Figure S20. mPNs connectivity features. a,** Fraction of total dendritic input onto olfactory mPNs from the neuronal classes indicated on the legend, left and right neuronal homologs are shown. **b,** Fraction of total mPN inputs coming from each OSN (right). Only connections present in both hemispheres of a species are shown. The plot on the left shows the divergence confidence score (see methods) of the differences in OSN input to mPNs between the two species, dots indicate strongly conserved connections (see methods).

**Figure S21. Dendritic input connectivity of mPNs.** Fraction of total inputs onto mPNs from each of the indicated neurons. Only connections with homologous neurons identified across both species and that are present in both hemispheres of a species are shown.

## Supplementary tables

**Supplementary table 1.** Results from the community analysis presented in Figure 1g.

**Supplementary table 2.** Classification of OSNs according to their described ecological significance

**Supplementary table 3.** Top 10 OSN-Picky LN-mPN IFL motifs for each species.

