## Supplementary Figures for "Cross-species comparative connectomics reveals the evolution of an olfactory circuit"

### Figure S1

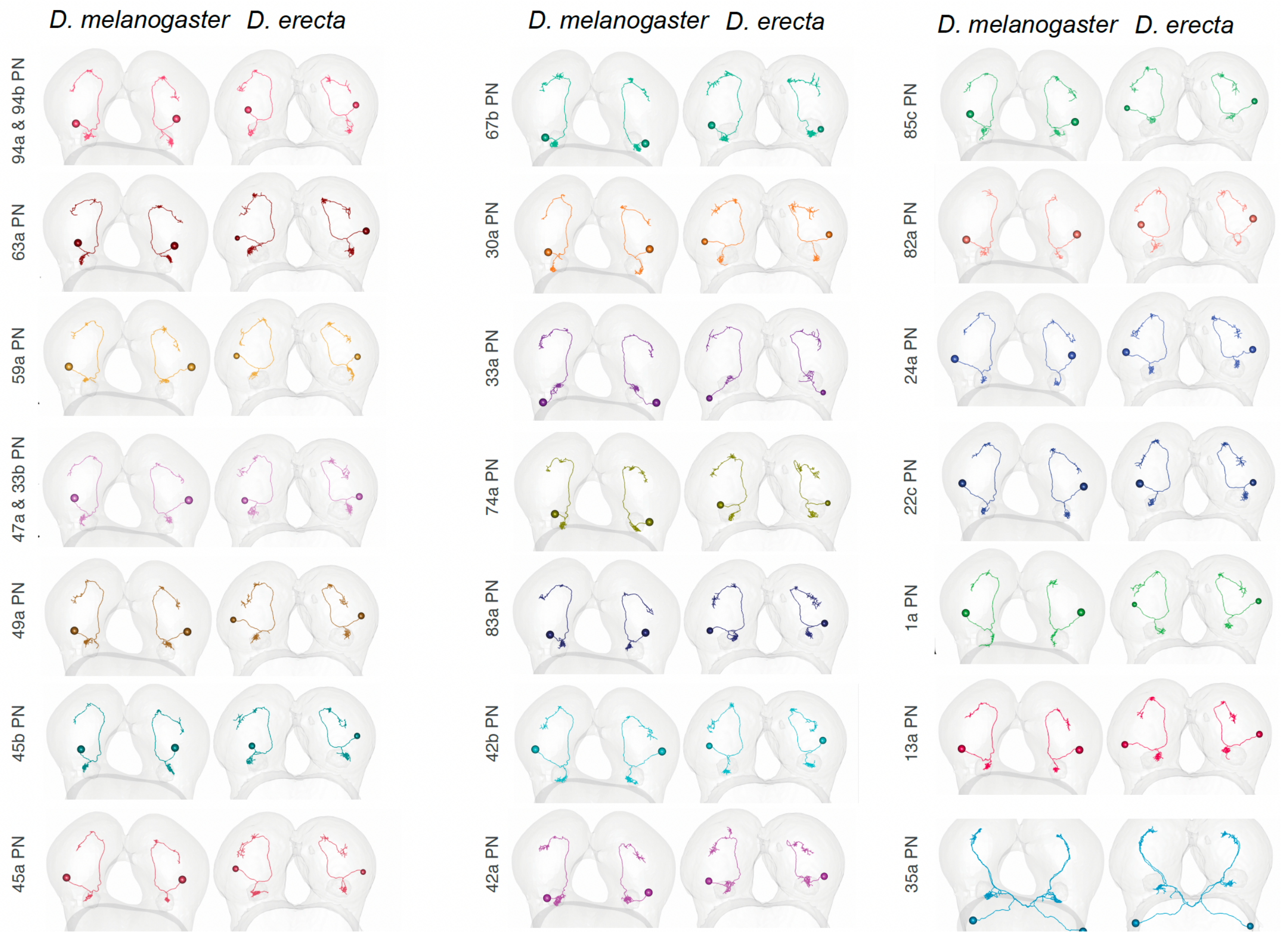

D. melanogaster
D. erecta
D. melanogaster
D. erecta
D. melanogaster
D. erecta
94a & 94b PN
63a PN
59a PN
47a & 33b PN
49a PN
45b PN
45a PN
67b PN
30a PN
33a PN
74a PN
83a PN
42b PN
42a PN
85c PN
82a PN
24a PN
22c PN
1a PN
13a PN
35a PN

### Figure S2

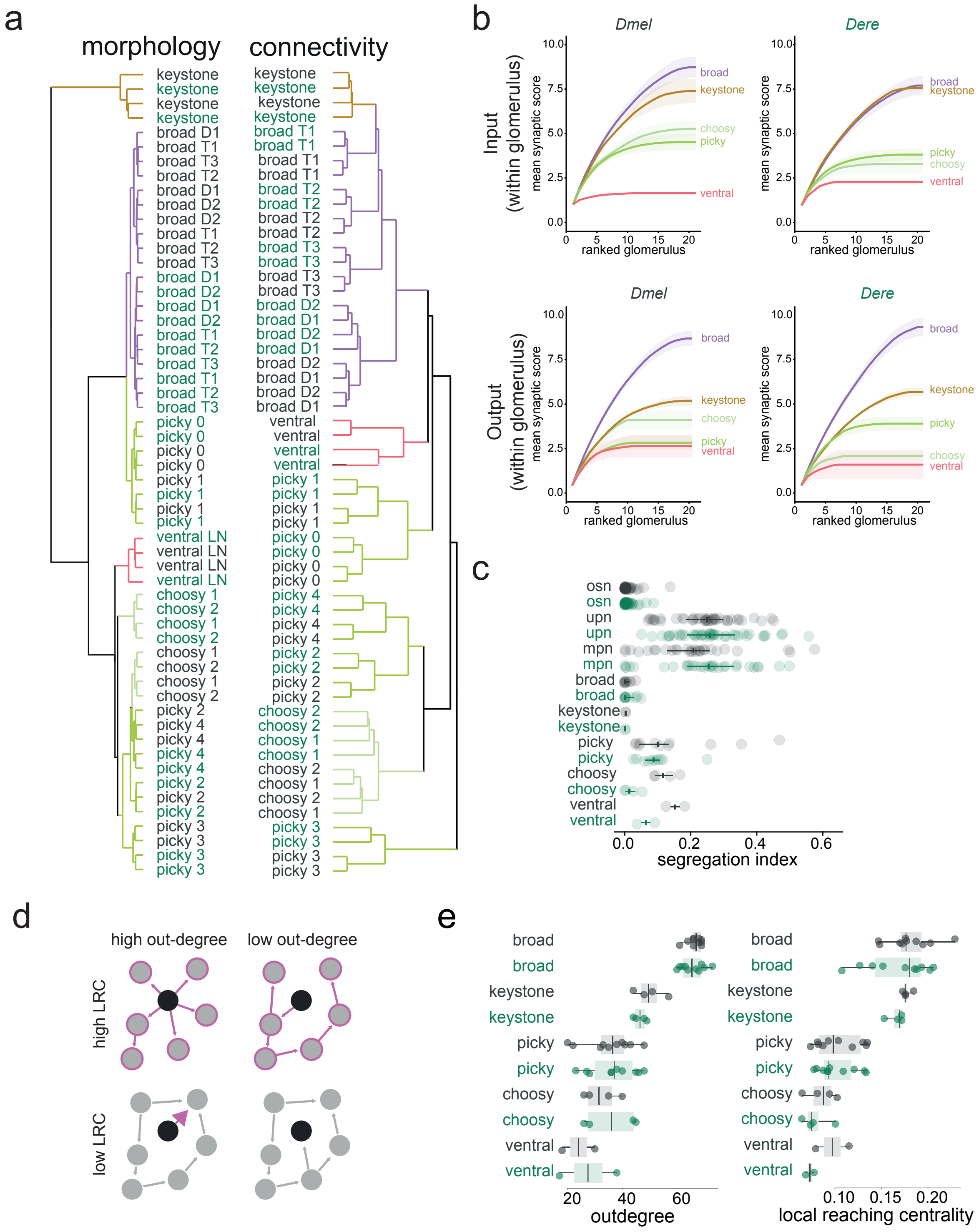

a
morphology
connectivity
keystone
broad D1
broad D2
broad T1
broad T2
broad T3
picky 0
picky 1
picky 2
picky 3
picky 4
ventral LN
ventral
choosy 1
choosy 2
b
Dmel
Dere
Input
(within glomerulus)
Output
(within glomerulus)
mean synaptic score
ranked glomerulus
broad
keystone
choosy
picky
ventral
c
osn
upn
mpn
broad
keystone
picky
choosy
ventral
segregation index
d
high out-degree
low out-degree
high LRC
low LRC
e
outdegree
local reaching centrality

### Figure S3

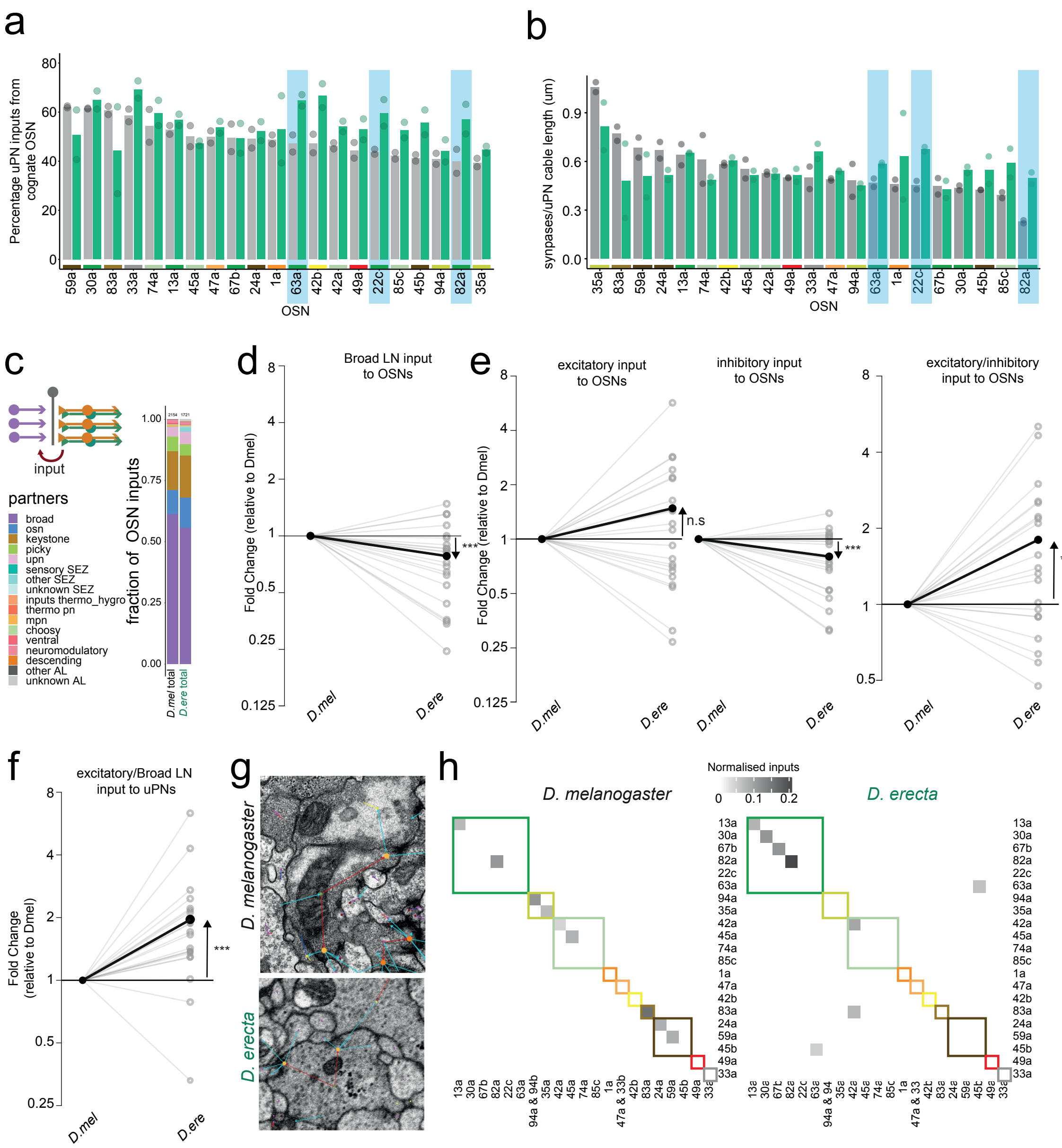

a
Percentage uPN inputs from cognate OSN
OSN
b
synapses/uPN cable length (um)
OSN
c
input
partners
broad
osn
keystone
picky
upn
sensory SEZ
other SEZ
unknown SEZ
inputs thermo_hygro
thermo pn
mpn
choosy
ventral
neuromodulatory
descending
other AL
unknown AL
fraction of OSN inputs
D.mel total
D.ere total
d
Broad LN input to OSNs
Fold Change (relative to Dmel)
D.mel
D.ere
e
excitatory input to OSNs
inhibitory input to OSNs
excitatory/inhibitory input to OSNs
n.s
f
excitatory/Broad LN input to uPNs
Fold Change (relative to Dmel)
g
D. melanogaster
D. erecta
h
Normalised inputs
D. melanogaster
D. erecta

### Figure S4

a

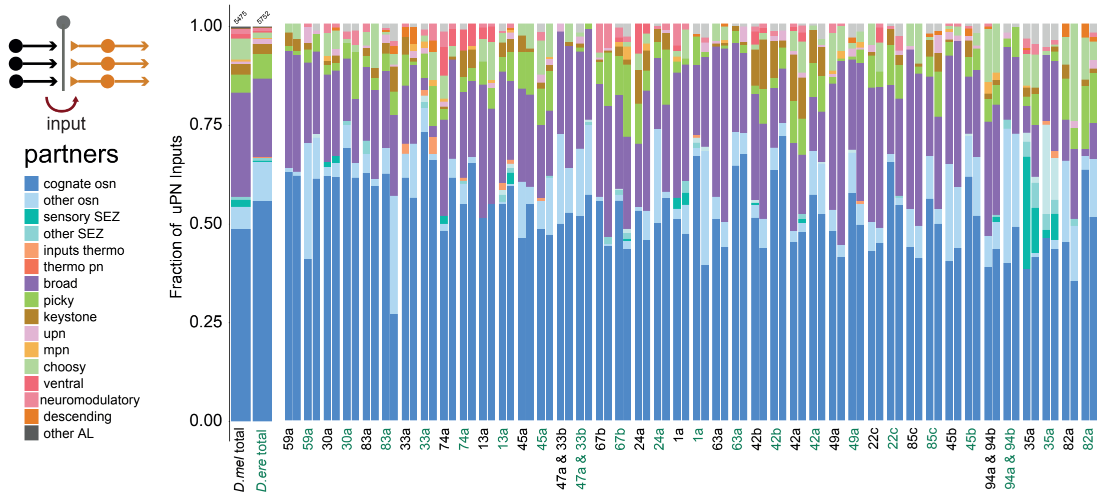
input
partners
cognate osn
other osn
sensory SEZ
other SEZ
inputs thermo
thermo pn
broad
picky
keystone
upn
mpn
choosy
ventral
neuromodulatory
descending
other AL
Fraction of uPN Inputs
1.00
0.75
0.50
0.25
0.00
D.mel total
D.ere total

b

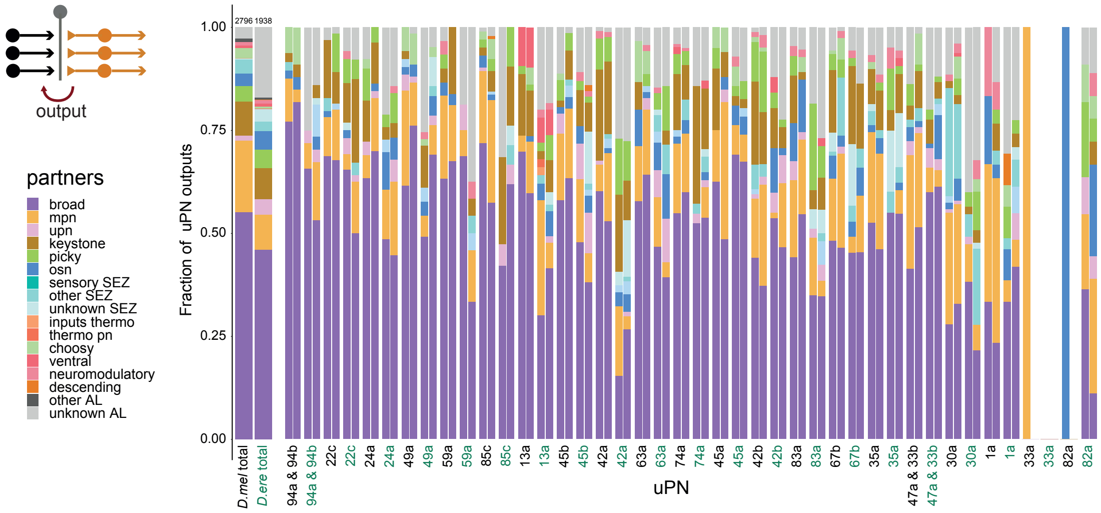
output
partners
broad
mpn
upn
keystone
picky
osn
sensory SEZ
other SEZ
unknown SEZ
inputs thermo
thermo pn
choosy
ventral
neuromodulatory
descending
other AL
unknown AL
Fraction of uPN outputs
1.00
0.75
0.50
0.25
0.00
2796
1938
D.mel total
D.ere total
uPN

Figure S5

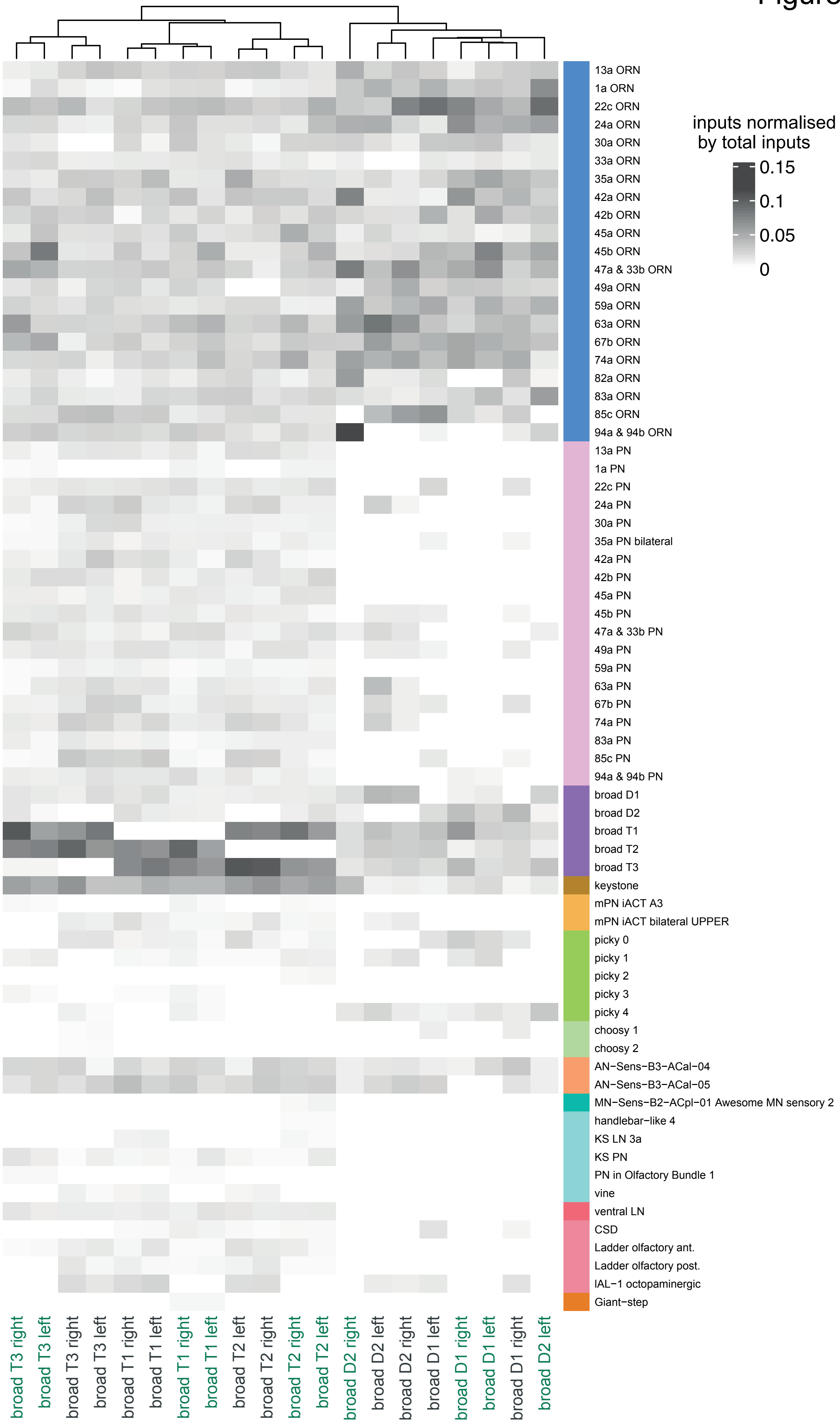
inputs normalised by total inputs
0.15
0.1
0.05
0
13a ORN
1a ORN
22c ORN
24a ORN
30a ORN
33a ORN
35a ORN
42a ORN
42b ORN
45a ORN
45b ORN
47a & 33b ORN
49a ORN
59a ORN
63a ORN
67b ORN
74a ORN
82a ORN
83a ORN
85c ORN
94a & 94b ORN
13a PN
1a PN
22c PN
24a PN
30a PN
35a PN bilateral
42a PN
42b PN
45a PN
45b PN
47a & 33b PN
49a PN
59a PN
63a PN
67b PN
74a PN
83a PN
85c PN
94a & 94b PN
broad D1
broad D2
broad T1
broad T2
broad T3
keystone
mPN iACT A3
mPN iACT bilateral UPPER
picky 0
picky 1
picky 2
picky 3
picky 4
choosy 1
choosy 2
AN−Sens−B3−ACal−04
AN−Sens−B3−ACal−05
MN−Sens−B2−ACpl−01 Awesome MN sensory 2
handlebar−like 4
KS LN 3a
KS PN
PN in Olfactory Bundle 1
vine
ventral LN
CSD
Ladder olfactory ant.
Ladder olfactory post.
IAL−1 octopaminergic
Giant−step
broad T3 right
broad T3 left
broad T3 right
broad T3 left
broad T1 right
broad T1 left
broad T1 right
broad T1 left
broad T2 left
broad T2 right
broad T2 right
broad T2 left
broad D2 right
broad D2 left
broad D2 right
broad D1 left
broad D1 right
broad D1 left
broad D1 right
broad D2 left

### Figure S6

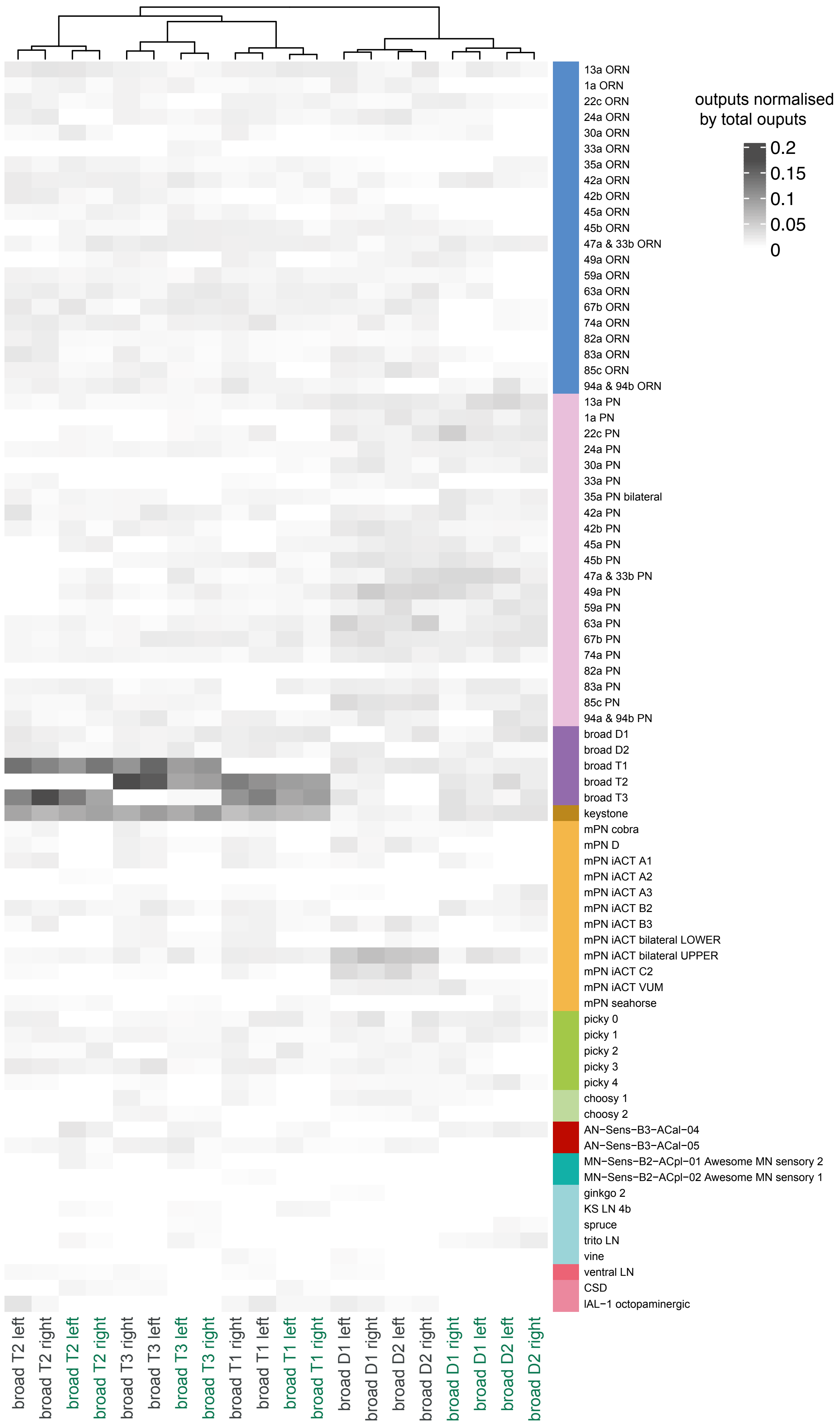
outputs normalised by total ouputs
0.2
0.15
0.1
0.05
0
13a ORN
1a ORN
22c ORN
24a ORN
30a ORN
33a ORN
35a ORN
42a ORN
42b ORN
45a ORN
45b ORN
47a & 33b ORN
49a ORN
59a ORN
63a ORN
67b ORN
74a ORN
82a ORN
83a ORN
85c ORN
94a & 94b ORN
13a PN
1a PN
22c PN
24a PN
30a PN
33a PN
35a PN bilateral
42a PN
42b PN
45a PN
45b PN
47a & 33b PN
49a PN
59a PN
63a PN
67b PN
74a PN
82a PN
83a PN
85c PN
94a & 94b PN
broad D1
broad D2
broad T1
broad T2
broad T3
keystone
mPN cobra
mPN D
mPN iACT A1
mPN iACT A2
mPN iACT A3
mPN iACT B2
mPN iACT B3
mPN iACT bilateral LOWER
mPN iACT bilateral UPPER
mPN iACT C2
mPN iACT VUM
mPN seahorse
picky 0
picky 1
picky 2
picky 3
picky 4
choosy 1
choosy 2
AN−Sens−B3−ACal−04
AN−Sens−B3−ACal−05
MN−Sens−B2−ACpl−01 Awesome MN sensory 2
MN−Sens−B2−ACpl−02 Awesome MN sensory 1
ginkgo 2
KS LN 4b
spruce
trito LN
vine
ventral LN
CSD
lAL−1 octopaminergic
broad T2 left
broad T2 right
broad T2 left
broad T2 right
broad T3 right
broad T3 left
broad T3 left
broad T3 right
broad T1 right
broad T1 left
broad T1 left
broad T1 right
broad D1 left
broad D1 right
broad D2 left
broad D2 right
broad D1 right
broad D1 left
broad D2 left
broad D2 right

### Figure S7

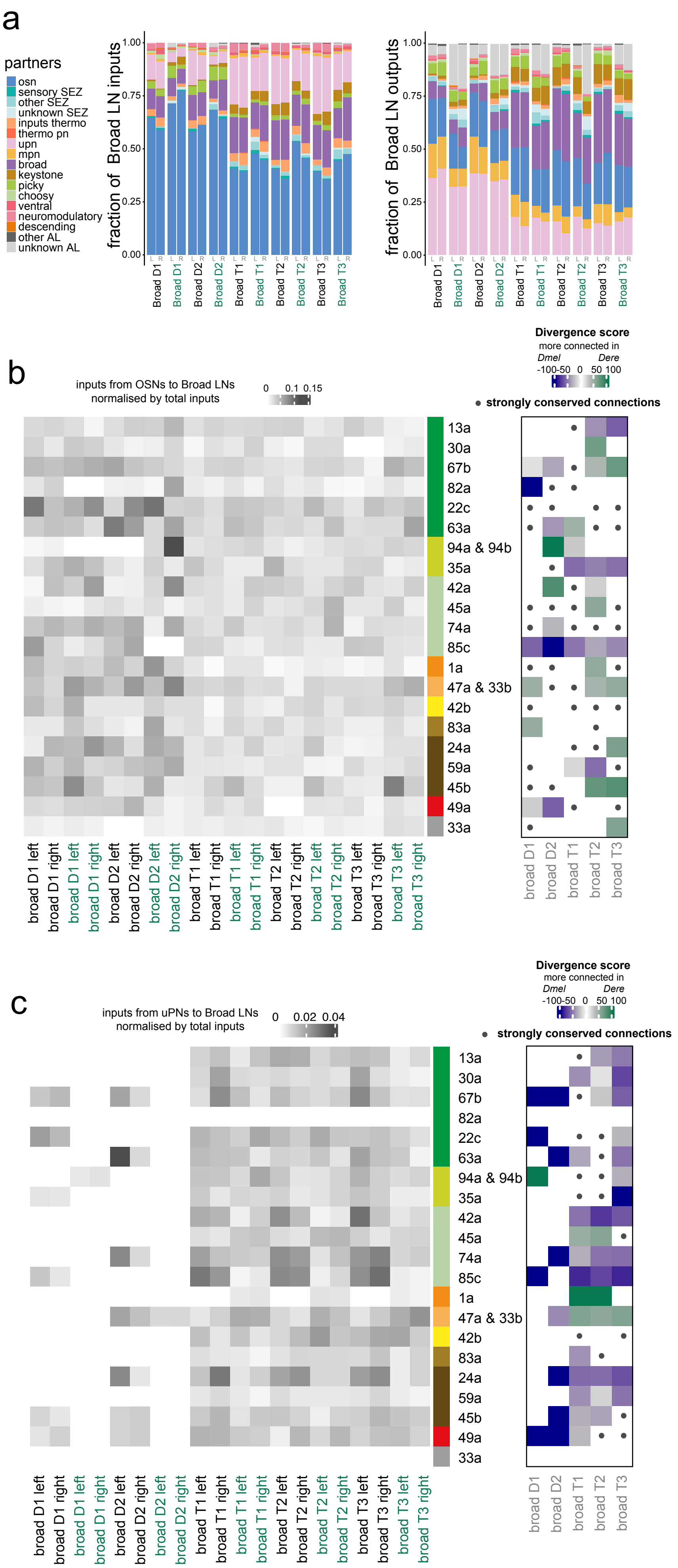
a
partners
osn
sensory SEZ
other SEZ
unknown SEZ
inputs thermo
thermo pn
upn
mpn
broad
keystone
picky
choosy
ventral
neuromodulatory
descending
other AL
unknown AL
fraction of Broad LN inputs
fraction of Broad LN outputs
0.00
0.25
0.50
0.75
1.00
Broad D1
Broad D2
Broad T1
Broad T2
Broad T3
b
inputs from OSNs to Broad LNs
normalised by total inputs
0
0.1
0.15
Divergence score
more connected in
Dmel
Dere
-100
-50
0
50
100
strongly conserved connections
13a
30a
67b
82a
22c
63a
94a & 94b
35a
42a
45a
74a
85c
1a
47a & 33b
42b
83a
24a
59a
45b
49a
33a
broad D1 left
broad D1 right
broad D2 left
broad D2 right
broad T1 left
broad T1 right
broad T2 left
broad T2 right
broad T3 left
broad T3 right
broad D1
broad D2
broad T1
broad T2
broad T3
c
inputs from uPNs to Broad LNs
normalised by total inputs
0
0.02
0.04

### Figure S8

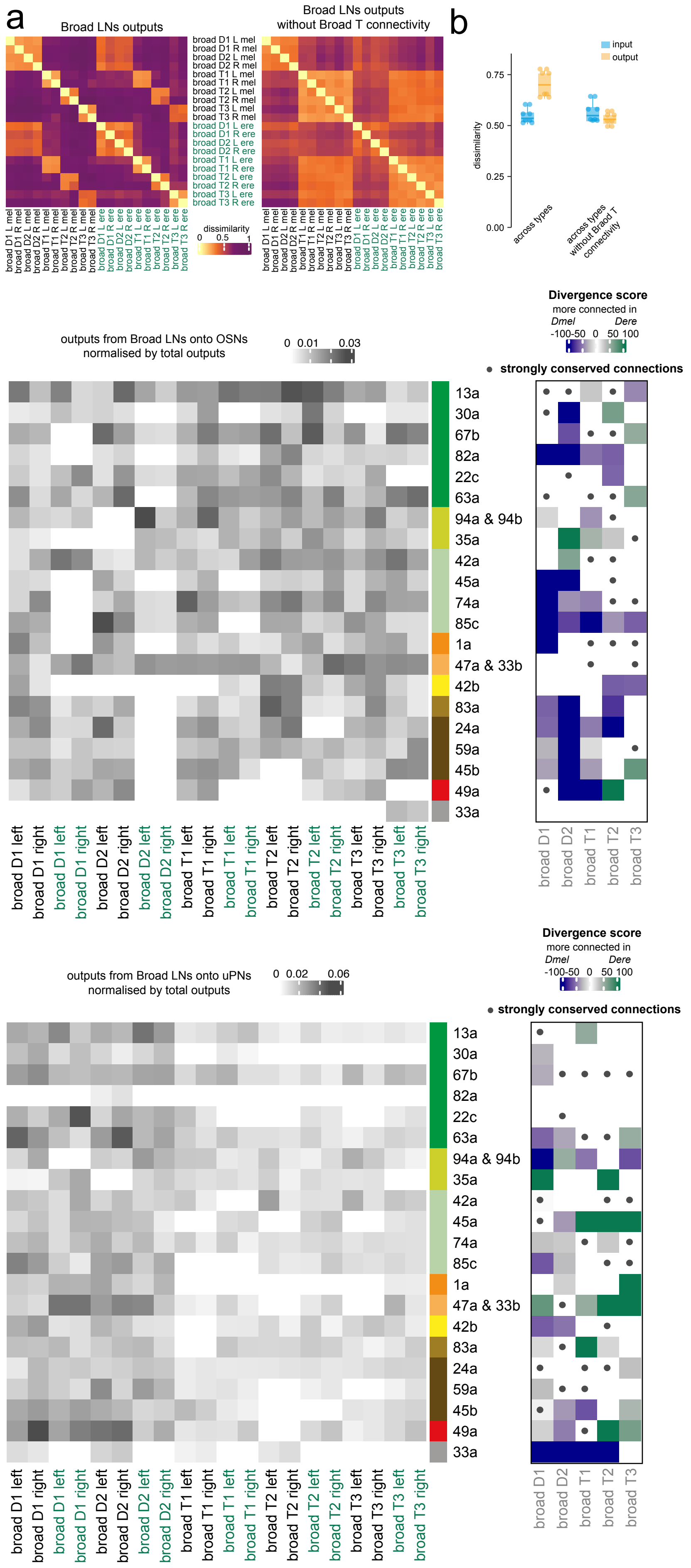
a
Broad LNs outputs
Broad LNs outputs without Broad T connectivity
broad D1 L mel
broad D1 R mel
broad D2 L mel
broad D2 R mel
broad T1 L mel
broad T1 R mel
broad T2 L mel
broad T2 R mel
broad T3 L mel
broad T3 R mel
broad D1 L ere
broad D1 R ere
broad D2 L ere
broad D2 R ere
broad T1 L ere
broad T1 R ere
broad T2 L ere
broad T2 R ere
broad T3 L ere
broad T3 R ere
dissimilarity
0 0.5 1
b
input
output
dissimilarity
0.00
0.25
0.50
0.75
across types
across types without Broad T connectivity
b
outputs from Broad LNs onto OSNs normalised by total outputs
0 0.01 0.03
Divergence score
more connected in
Dmel Dere
-100-50 0 50 100
strongly conserved connections
13a
30a
67b
82a
22c
63a
94a & 94b
35a
42a
45a
74a
85c
1a
47a & 33b
42b
83a
24a
59a
45b
49a
33a
broad D1 left
broad D1 right
broad D1 left
broad D1 right
broad D2 left
broad D2 right
broad D2 left
broad D2 right
broad T1 left
broad T1 right
broad T1 left
broad T1 right
broad T2 left
broad T2 right
broad T2 left
broad T2 right
broad T3 left
broad T3 right
broad T3 left
broad T3 right
broad D1
broad D2
broad T1
broad T2
broad T3
c
outputs from Broad LNs onto uPNs normalised by total outputs
0 0.02 0.06
Divergence score
more connected in
Dmel Dere
-100-50 0 50 100
strongly conserved connections

Figure S9

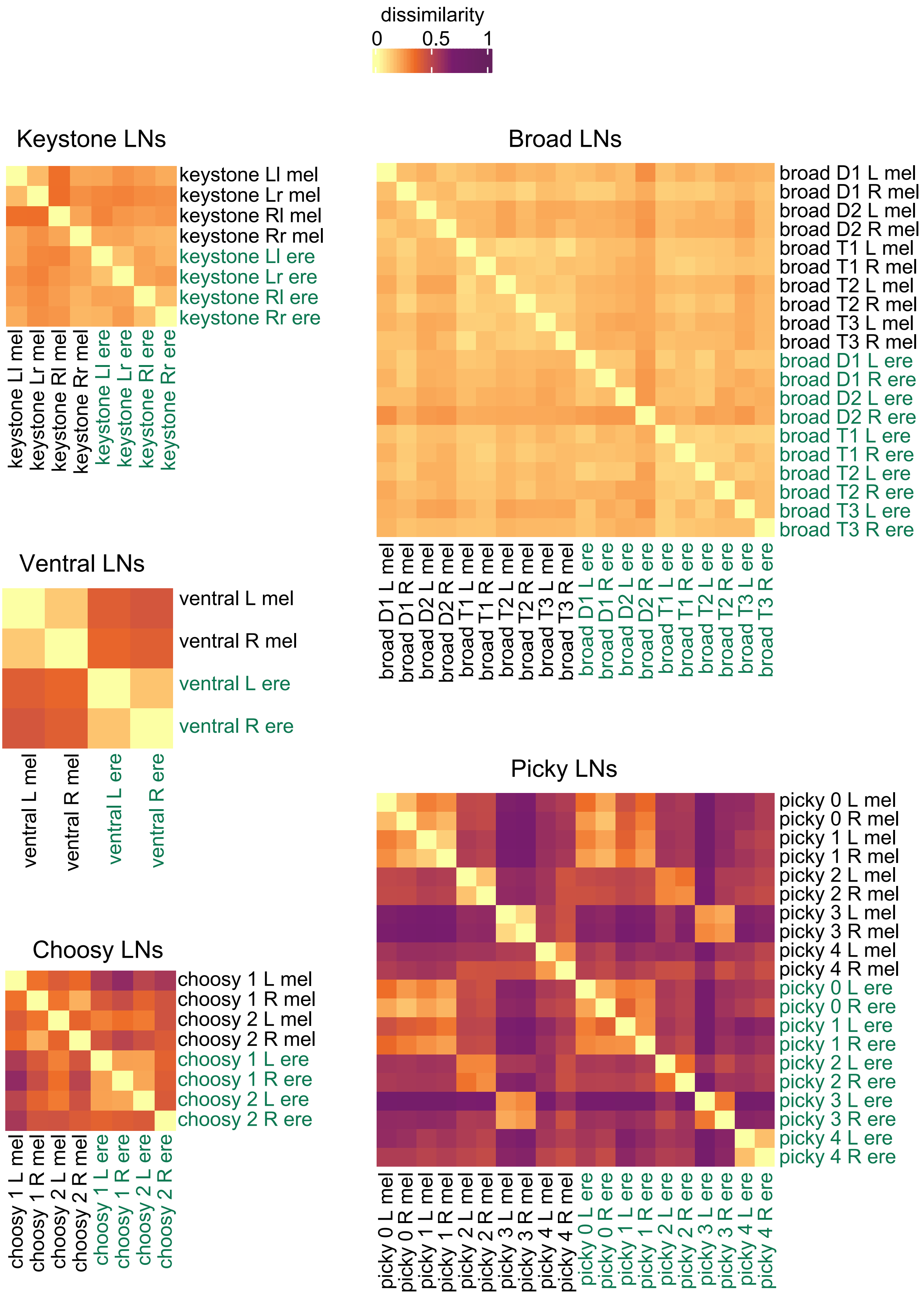
dissimilarity
0
0.5
1
Keystone LNs
keystone Ll mel
keystone Lr mel
keystone Rl mel
keystone Rr mel
keystone Ll ere
keystone Lr ere
keystone Rl ere
keystone Rr ere
Broad LNs
broad D1 L mel
broad D1 R mel
broad D2 L mel
broad D2 R mel
broad T1 L mel
broad T1 R mel
broad T2 L mel
broad T2 R mel
broad T3 L mel
broad T3 R mel
broad D1 L ere
broad D1 R ere
broad D2 L ere
broad D2 R ere
broad T1 L ere
broad T1 R ere
broad T2 L ere
broad T2 R ere
broad T3 L ere
broad T3 R ere
Ventral LNs
ventral L mel
ventral R mel
ventral L ere
ventral R ere
Picky LNs
picky 0 L mel
picky 0 R mel
picky 1 L mel
picky 1 R mel
picky 2 L mel
picky 2 R mel
picky 3 L mel
picky 3 R mel
picky 4 L mel
picky 4 R mel
picky 0 L ere
picky 0 R ere
picky 1 L ere
picky 1 R ere
picky 2 L ere
picky 2 R ere
picky 3 L ere
picky 3 R ere
picky 4 L ere
picky 4 R ere
Choosy LNs
choosy 1 L mel
choosy 1 R mel
choosy 2 L mel
choosy 2 R mel
choosy 1 L ere
choosy 1 R ere
choosy 2 L ere
choosy 2 R ere

Figure S10

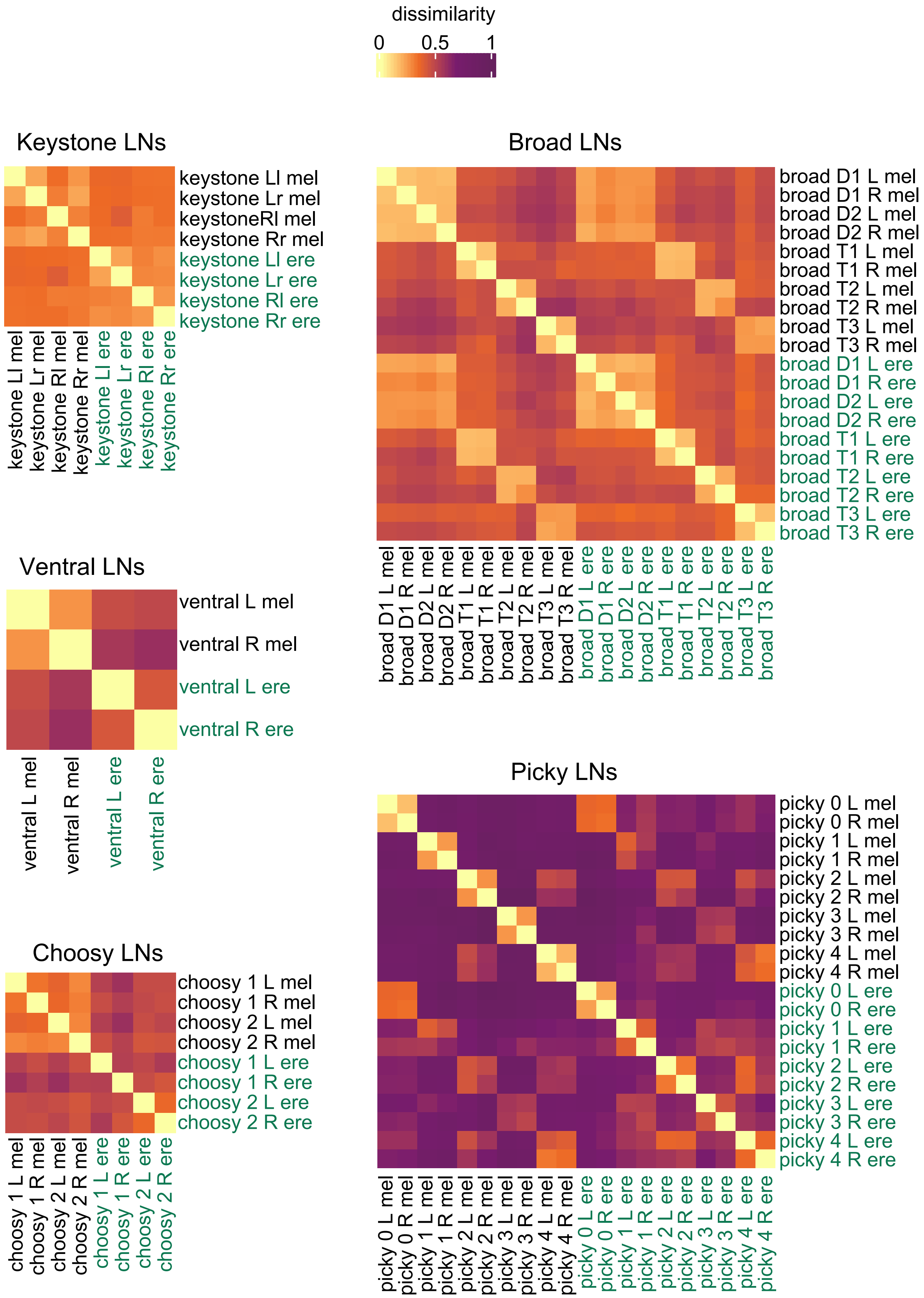
dissimilarity
0
0.5
1
Keystone LNs
keystone Ll mel
keystone Lr mel
keystoneRl mel
keystone Rr mel
keystone Ll ere
keystone Lr ere
keystone Rl ere
keystone Rr ere
Broad LNs
broad D1 L mel
broad D1 R mel
broad D2 L mel
broad D2 R mel
broad T1 L mel
broad T1 R mel
broad T2 L mel
broad T2 R mel
broad T3 L mel
broad T3 R mel
broad D1 L ere
broad D1 R ere
broad D2 L ere
broad D2 R ere
broad T1 L ere
broad T1 R ere
broad T2 L ere
broad T2 R ere
broad T3 L ere
broad T3 R ere
Ventral LNs
ventral L mel
ventral R mel
ventral L ere
ventral R ere
Picky LNs
picky 0 L mel
picky 0 R mel
picky 1 L mel
picky 1 R mel
picky 2 L mel
picky 2 R mel
picky 3 L mel
picky 3 R mel
picky 4 L mel
picky 4 R mel
picky 0 L ere
picky 0 R ere
picky 1 L ere
picky 1 R ere
picky 2 L ere
picky 2 R ere
picky 3 L ere
picky 3 R ere
picky 4 L ere
picky 4 R ere
Choosy LNs
choosy 1 L mel
choosy 1 R mel
choosy 2 L mel
choosy 2 R mel
choosy 1 L ere
choosy 1 R ere
choosy 2 L ere
choosy 2 R ere

### Figure S11

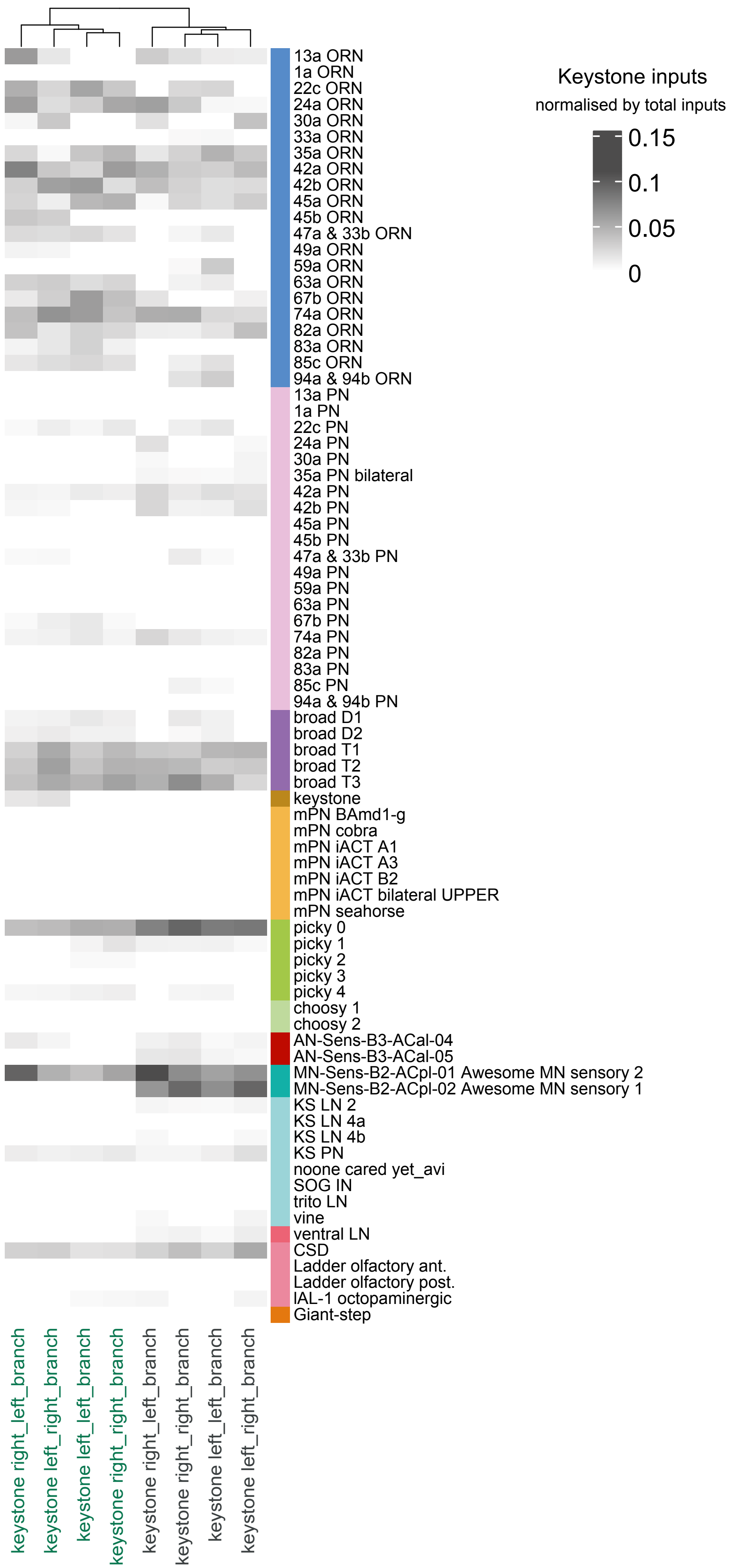
Keystone inputs
normalised by total inputs
0.15
0.1
0.05
0
13a ORN
1a ORN
22c ORN
24a ORN
30a ORN
33a ORN
35a ORN
42a ORN
42b ORN
45a ORN
45b ORN
47a & 33b ORN
49a ORN
59a ORN
63a ORN
67b ORN
74a ORN
82a ORN
83a ORN
85c ORN
94a & 94b ORN
13a PN
1a PN
22c PN
24a PN
30a PN
35a PN bilateral
42a PN
42b PN
45a PN
45b PN
47a & 33b PN
49a PN
59a PN
63a PN
67b PN
74a PN
82a PN
83a PN
85c PN
94a & 94b PN
broad D1
broad D2
broad T1
broad T2
broad T3
keystone
mPN BAmd1-g
mPN cobra
mPN iACT A1
mPN iACT A3
mPN iACT B2
mPN iACT bilateral UPPER
mPN seahorse
picky 0
picky 1
picky 2
picky 3
picky 4
choosy 1
choosy 2
AN-Sens-B3-ACal-04
AN-Sens-B3-ACal-05
MN-Sens-B2-ACpl-01 Awesome MN sensory 2
MN-Sens-B2-ACpl-02 Awesome MN sensory 1
KS LN 2
KS LN 4a
KS LN 4b
KS PN
noone cared yet_avi
SOG IN
trito LN
vine
ventral LN
CSD
Ladder olfactory ant.
Ladder olfactory post.
lAL-1 octopaminergic
Giant-step
keystone right_left_branch
keystone left_right_branch
keystone left_left_branch
keystone right_right_branch
keystone right_left_branch
keystone right_right_branch
keystone left_left_branch
keystone left_right_branch

### Figure S12

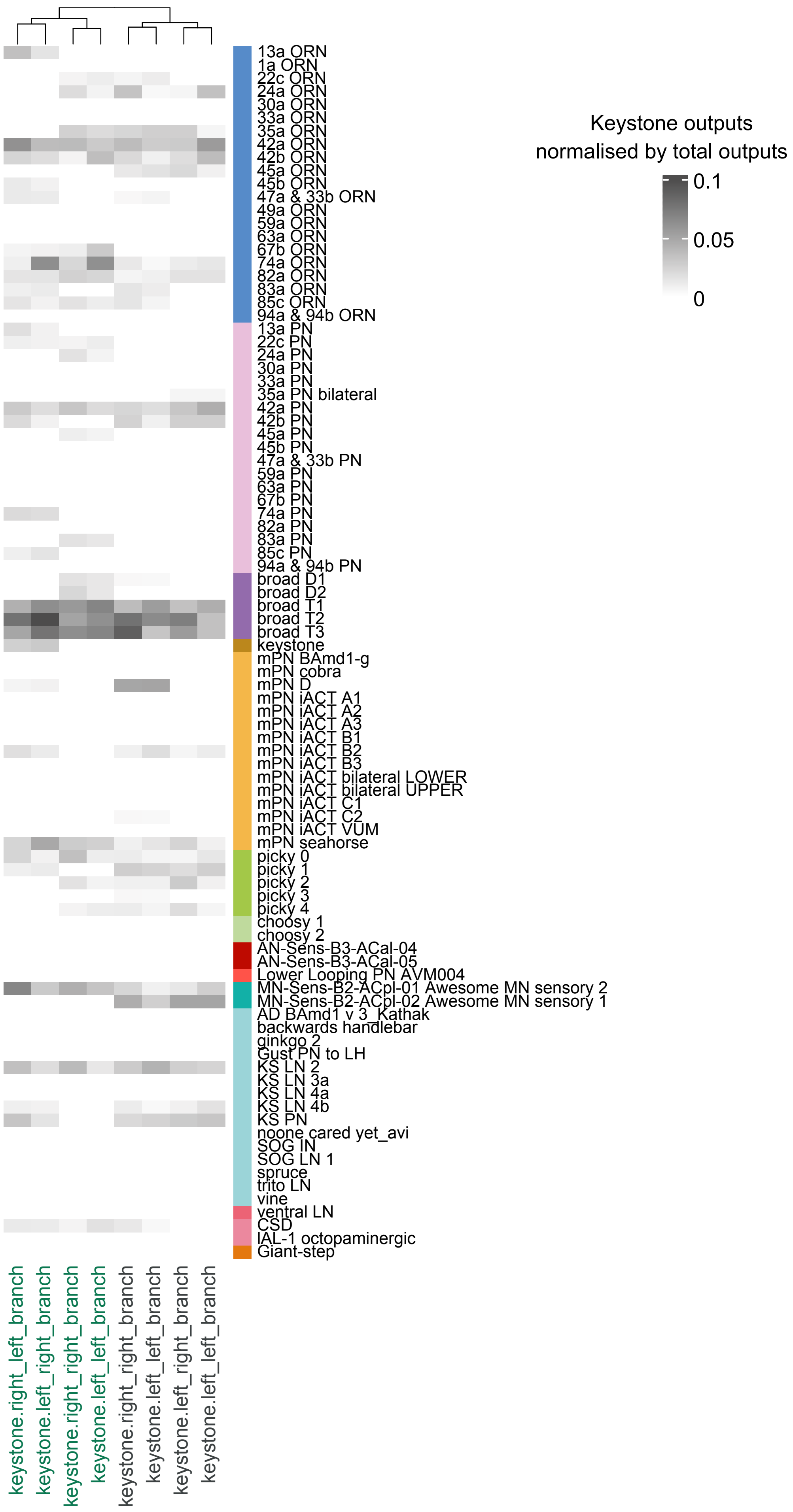
Keystone outputs
normalised by total outputs
0.1
0.05
0
13a ORN
1a ORN
22c ORN
24a ORN
30a ORN
33a ORN
35a ORN
42a ORN
42b ORN
45a ORN
45b ORN
47a & 33b ORN
49a ORN
59a ORN
63a ORN
67b ORN
74a ORN
82a ORN
83a ORN
85c ORN
94a & 94b ORN
13a PN
22c PN
24a PN
30a PN
33a PN
35a PN bilateral
42a PN
42b PN
45a PN
45b PN
47a & 33b PN
59a PN
63a PN
67b PN
74a PN
82a PN
83a PN
85c PN
94a & 94b PN
broad D1
broad D2
broad T1
broad T2
broad T3
keystone
mPN BAmd1-g
mPN cobra
mPN D
mPN iACT A1
mPN iACT A2
mPN iACT A3
mPN iACT B1
mPN iACT B2
mPN iACT B3
mPN iACT bilateral LOWER
mPN iACT bilateral UPPER
mPN iACT C1
mPN iACT C2
mPN iACT VUM
mPN seahorse
picky 0
picky 1
picky 2
picky 3
picky 4
choosy 1
choosy 2
AN-Sens-B3-ACal-04
AN-Sens-B3-ACal-05
Lower Looping PN AVM004
MN-Sens-B2-ACpl-01 Awesome MN sensory 2
MN-Sens-B2-ACpl-02 Awesome MN sensory 1
AD BAmd1 v 3_Kathak
backwards handlebar
ginkgo 2
Gust PN to LH
KS LN 2
KS LN 3a
KS LN 4a
KS LN 4b
KS PN
noone cared yet_avi
SOG IN
SOG LN 1
spruce
trito LN
vine
ventral LN
CSD
IAL-1 octopaminergic
Giant-step
keystone.right_left_branch
keystone.left_right_branch
keystone.right_right_branch
keystone.left_left_branch
keystone.right_right_branch
keystone.left_left_branch
keystone.left_right_branch
keystone.left_left_branch

### Figure S13

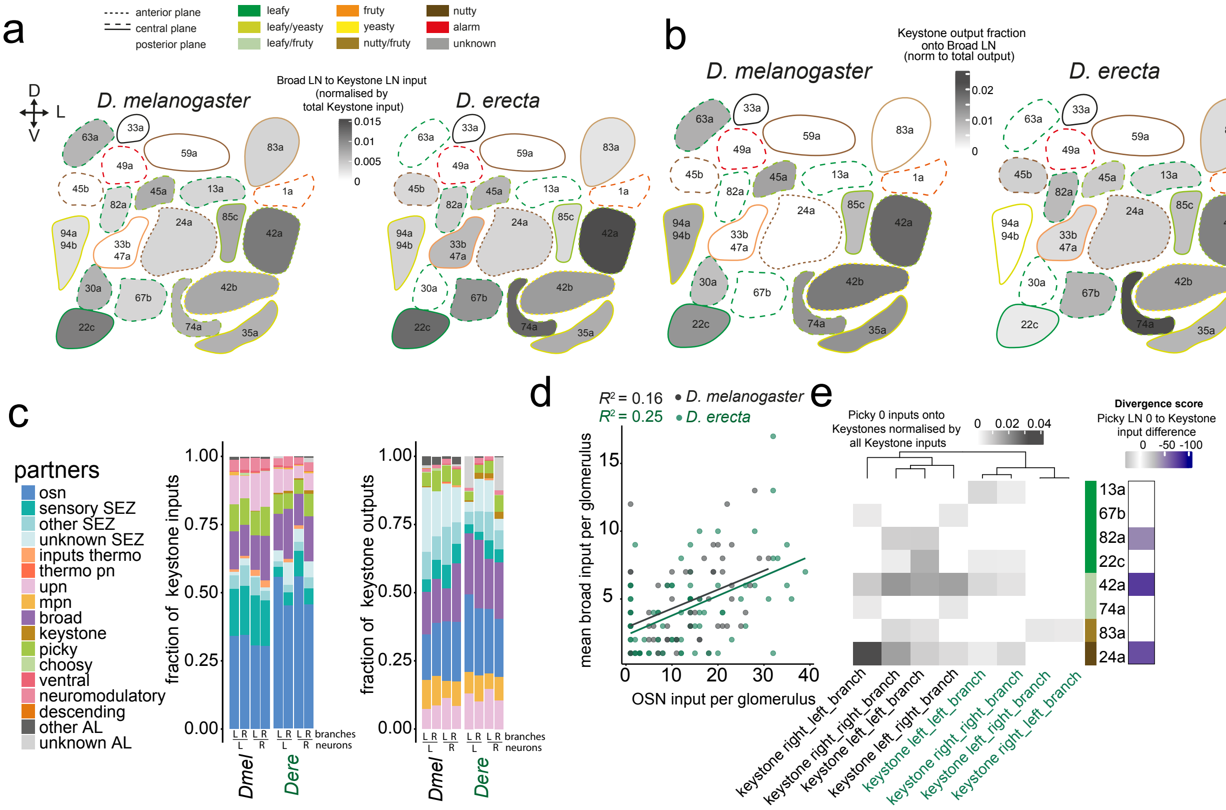
a
anterior plane
central plane
posterior plane
leafy
leafy/yeasty
leafy/fruty
fruty
yeasty
nutty/fruty
nutty
alarm
unknown
D
L
V
D. melanogaster
D. erecta
Broad LN to Keystone LN input
(normalised by
total Keystone input)
0.015
0.01
0.005
0
63a
33a
49a
59a
83a
45b
45a
13a
1a
82a
24a
85c
42a
94a
94b
33b
47a
30a
67b
42b
22c
74a
35a
b
Keystone output fraction
onto Broad LN
(norm to total output)
0.02
0.01
0
c
partners
osn
sensory SEZ
other SEZ
unknown SEZ
inputs thermo
thermo pn
upn
mpn
broad
keystone
picky
choosy
ventral
neuromodulatory
descending
other AL
unknown AL
fraction of keystone inputs
fraction of keystone outputs
1.00
0.75
0.50
0.25
0.00
L R L R L R L R branches
L R L R neurons
Dmel
Dere
d
$R^2$ = 0.16 • D. melanogaster
$R^2$ = 0.25 • D. erecta
mean broad input per glomerulus
OSN input per glomerulus
15
10
5
0
10
20
30
40
e
Picky 0 inputs onto
Keystones normalised by
all Keystone inputs
0
0.02
0.04
Divergence score
Picky LN 0 to Keystone
input difference
0
-50
-100
13a
67b
82a
22c
42a
74a
83a
24a
keystone right_left_branch
keystone right_right_branch
keystone left_left_branch
keystone left_right_branch
keystone left_left_branch
keystone right_right_branch
keystone left_right_branch
keystone right_left_branch

### Figure S14

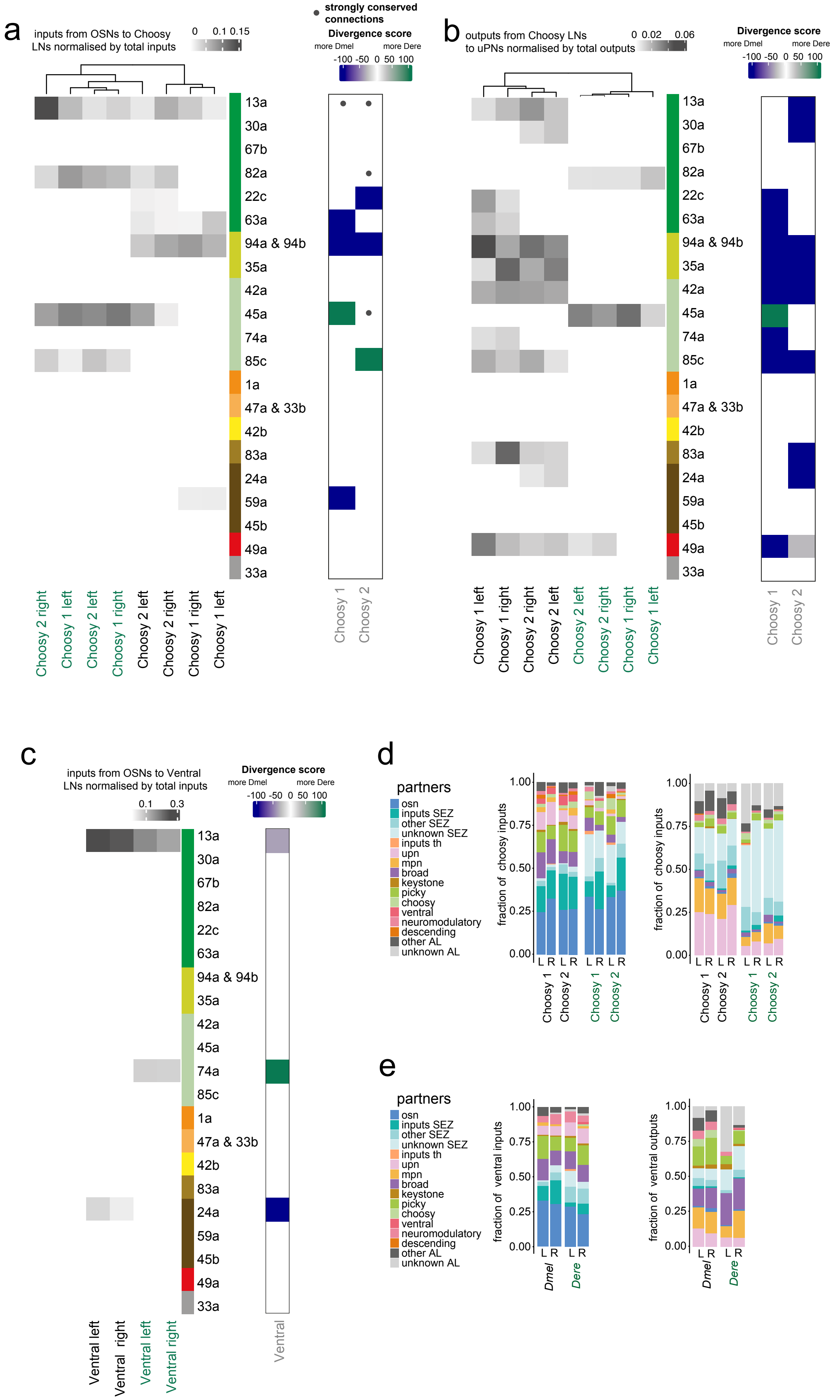

a
inputs from OSNs to Choosy LNs normalised by total inputs
0 0.1 0.15
strongly conserved connections
Divergence score
more Dmel
more Dere
-100 -50 0 50 100
13a
30a
67b
82a
22c
63a
94a & 94b
35a
42a
45a
74a
85c
1a
47a & 33b
42b
83a
24a
59a
45b
49a
33a
Choosy 2 right
Choosy 1 left
Choosy 2 left
Choosy 1 right
Choosy 2 left
Choosy 2 right
Choosy 1 right
Choosy 1 left
Choosy 1
Choosy 2
b
outputs from Choosy LNs to uPNs normalised by total outputs
0 0.02 0.06
Choosy 1 left
Choosy 1 right
Choosy 2 right
Choosy 2 left
Choosy 2 left
Choosy 2 right
Choosy 1 right
Choosy 1 left
c
inputs from OSNs to Ventral LNs normalised by total inputs
0.1 0.3
Ventral left
Ventral right
Ventral left
Ventral right
Ventral
d
partners
osn
inputs SEZ
other SEZ
unknown SEZ
inputs th
upn
mpn
broad
keystone
picky
choosy
ventral
neuromodulatory
descending
other AL
unknown AL
fraction of choosy inputs
fraction of choosy inputs
1.00
0.75
0.50
0.25
0.00
L R
Choosy 1
Choosy 2
e
fraction of ventral inputs
fraction of ventral outputs
Dmel
Dere

Figure S15

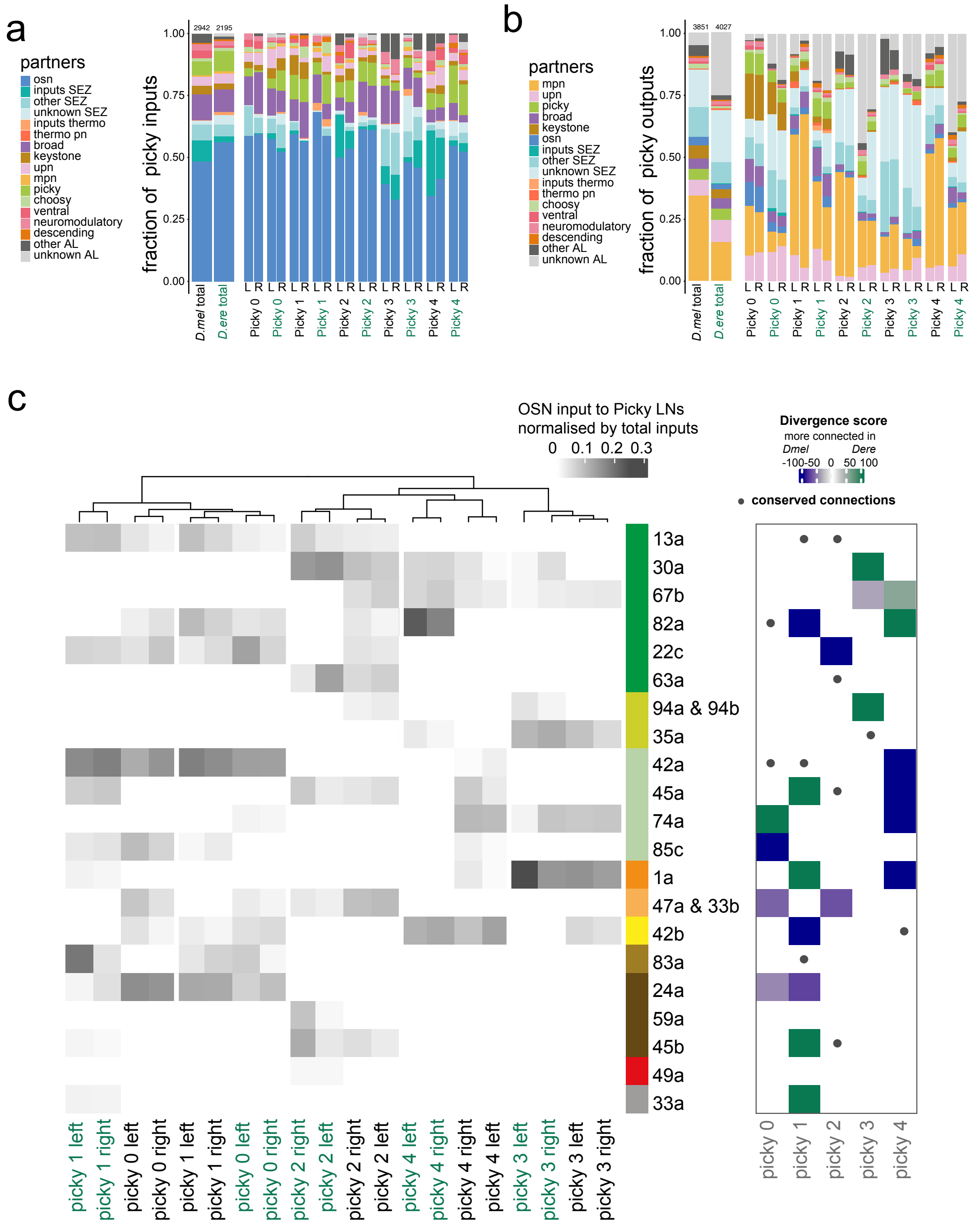
a
partners
osn
inputs SEZ
other SEZ
unknown SEZ
inputs thermo
thermo pn
broad
keystone
upn
mpn
picky
choosy
ventral
neuromodulatory
descending
other AL
unknown AL
fraction of picky inputs
1.00
0.75
0.50
0.25
0.00
2942
2195
D.mel total
D.ere total
Picky 0
Picky 0
Picky 1
Picky 1
Picky 2
Picky 2
Picky 3
Picky 3
Picky 4
Picky 4
L R
b
partners
mpn
upn
picky
broad
keystone
osn
inputs SEZ
other SEZ
unknown SEZ
inputs thermo
thermo pn
choosy
ventral
neuromodulatory
descending
other AL
unknown AL
fraction of picky outputs
3851
4027
c
OSN input to Picky LNs
normalised by total inputs
0 0.1 0.2 0.3
Divergence score
more connected in
Dmel Dere
-100-50 0 50 100
conserved connections
13a
30a
67b
82a
22c
63a
94a & 94b
35a
42a
45a
74a
85c
1a
47a & 33b
42b
83a
24a
59a
45b
49a
33a
picky 1 left
picky 1 right
picky 0 left
picky 0 right
picky 1 left
picky 1 right
picky 0 left
picky 0 right
picky 2 right
picky 2 left
picky 2 right
picky 2 left
picky 4 left
picky 4 right
picky 4 right
picky 4 left
picky 3 left
picky 3 right
picky 3 left
picky 3 right
picky 0
picky 1
picky 2
picky 3
picky 4

### Figure S16

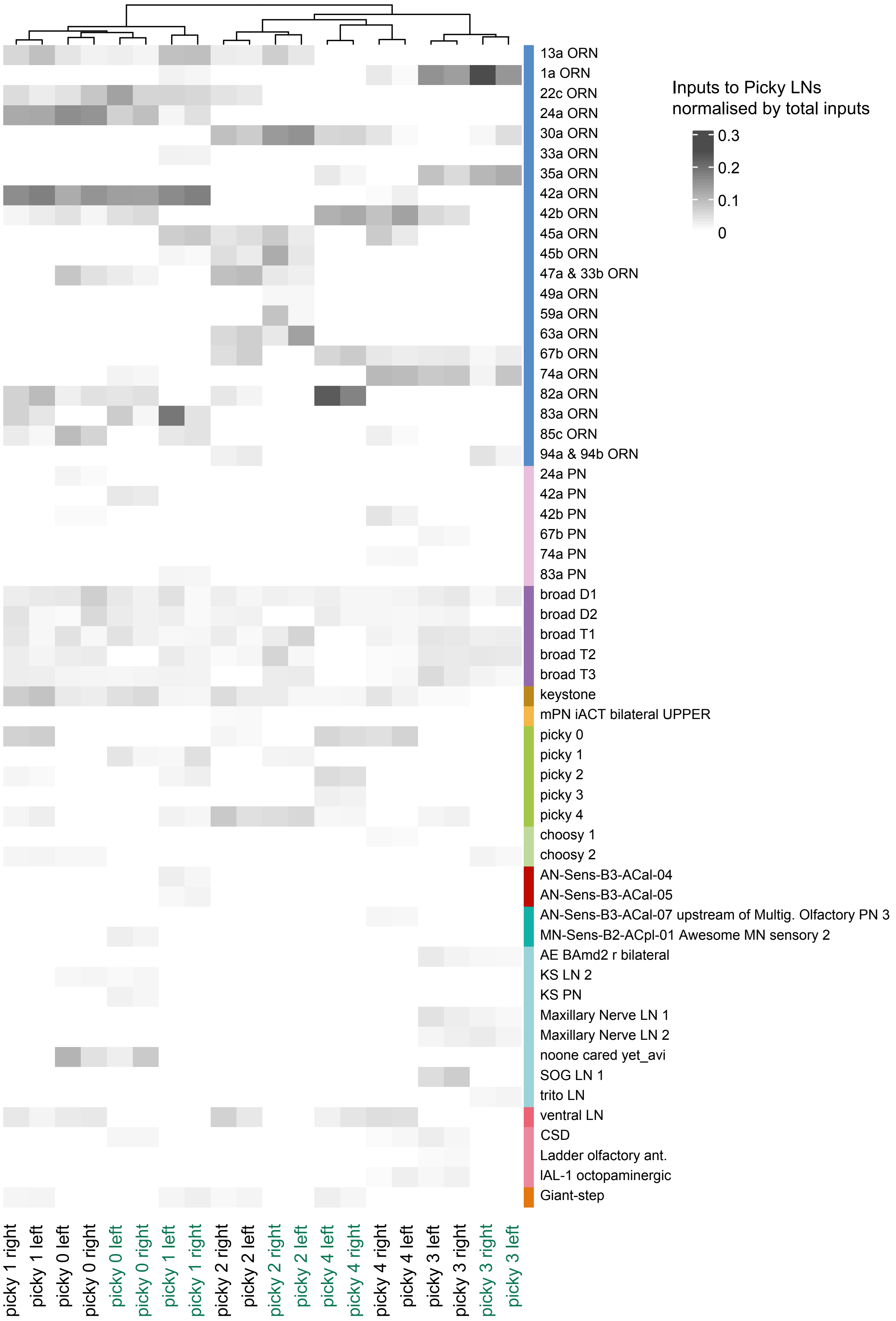
Inputs to Picky LNs
normalised by total inputs
0.3
0.2
0.1
0
13a ORN
1a ORN
22c ORN
24a ORN
30a ORN
33a ORN
35a ORN
42a ORN
42b ORN
45a ORN
45b ORN
47a & 33b ORN
49a ORN
59a ORN
63a ORN
67b ORN
74a ORN
82a ORN
83a ORN
85c ORN
94a & 94b ORN
24a PN
42a PN
42b PN
67b PN
74a PN
83a PN
broad D1
broad D2
broad T1
broad T2
broad T3
keystone
mPN iACT bilateral UPPER
picky 0
picky 1
picky 2
picky 3
picky 4
choosy 1
choosy 2
AN-Sens-B3-ACal-04
AN-Sens-B3-ACal-05
AN-Sens-B3-ACal-07 upstream of Multig. Olfactory PN 3
MN-Sens-B2-ACpl-01 Awesome MN sensory 2
AE BAmd2 r bilateral
KS LN 2
KS PN
Maxillary Nerve LN 1
Maxillary Nerve LN 2
noone cared yet_avi
SOG LN 1
trito LN
ventral LN
CSD
Ladder olfactory ant.
lAL-1 octopaminergic
Giant-step
picky 1 right
picky 1 left
picky 0 left
picky 0 right
picky 0 left
picky 0 right
picky 1 left
picky 1 right
picky 2 right
picky 2 left
picky 2 right
picky 2 left
picky 4 left
picky 4 right
picky 4 right
picky 4 left
picky 3 left
picky 3 right
picky 3 right
picky 3 left

### Figure S17

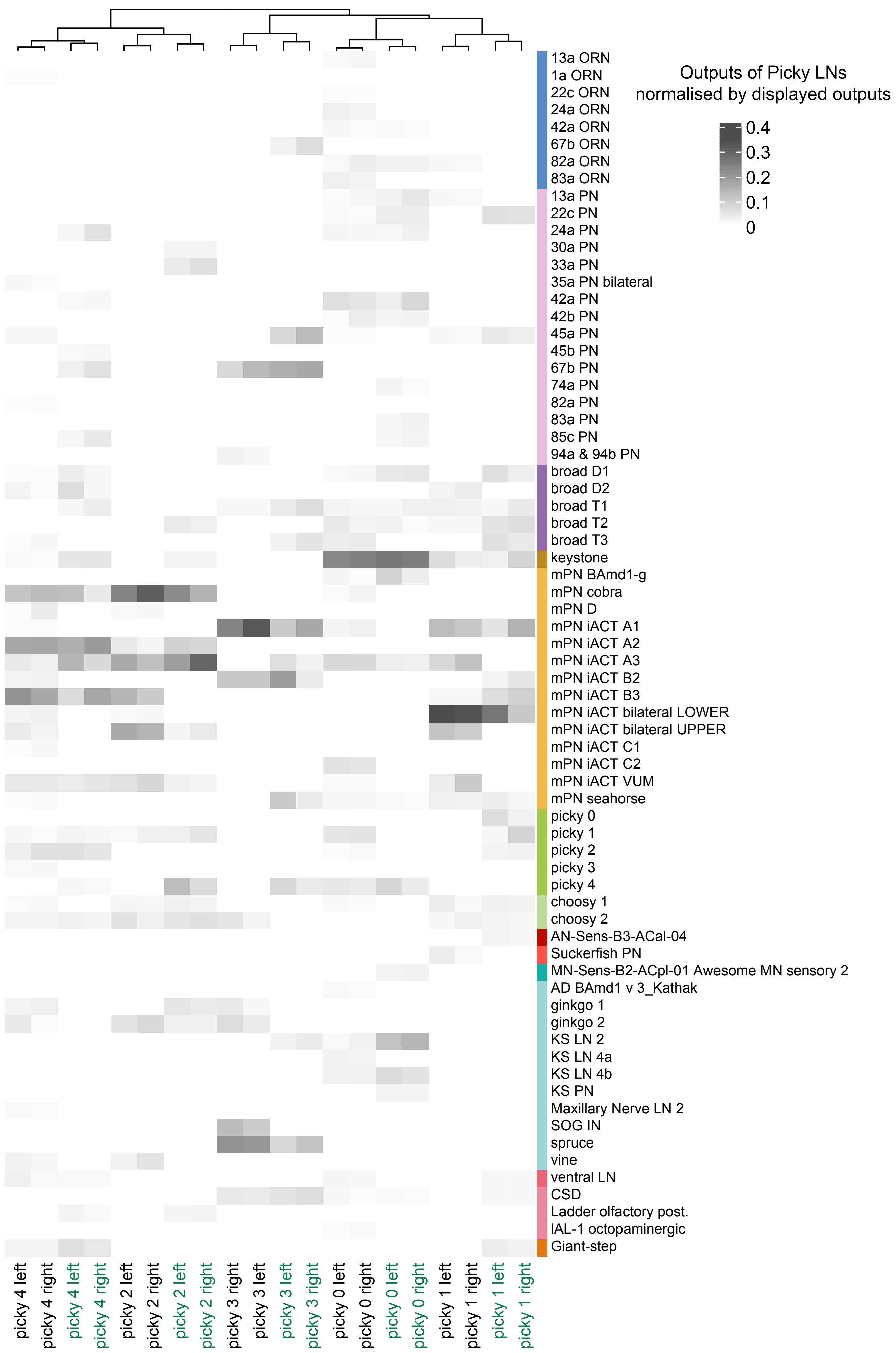

Outputs of Picky LNs
normalised by displayed outputs
0.4
0.3
0.2
0.1
0
13a ORN
1a ORN
22c ORN
24a ORN
42a ORN
67b ORN
82a ORN
83a ORN
13a PN
22c PN
24a PN
30a PN
33a PN
35a PN bilateral
42a PN
42b PN
45a PN
45b PN
67b PN
74a PN
82a PN
83a PN
85c PN
94a & 94b PN
broad D1
broad D2
broad T1
broad T2
broad T3
keystone
mPN BAmd1-g
mPN cobra
mPN D
mPN iACT A1
mPN iACT A2
mPN iACT A3
mPN iACT B2
mPN iACT B3
mPN iACT bilateral LOWER
mPN iACT bilateral UPPER
mPN iACT C1
mPN iACT C2
mPN iACT VUM
mPN seahorse
picky 0
picky 1
picky 2
picky 3
picky 4
choosy 1
choosy 2
AN-Sens-B3-ACal-04
Suckerfish PN
MN-Sens-B2-ACpl-01 Awesome MN sensory 2
AD BAmd1 v 3_Kathak
ginkgo 1
ginkgo 2
KS LN 2
KS LN 4a
KS LN 4b
KS PN
Maxillary Nerve LN 2
SOG IN
spruce
vine
ventral LN
CSD
Ladder olfactory post.
lAL-1 octopaminergic
Giant-step
picky 4 left
picky 4 right
picky 4 left
picky 4 right
picky 2 left
picky 2 right
picky 2 left
picky 2 right
picky 3 right
picky 3 left
picky 3 left
picky 3 right
picky 0 left
picky 0 right
picky 0 left
picky 0 right
picky 1 left
picky 1 right
picky 1 left
picky 1 right

### Figure S18

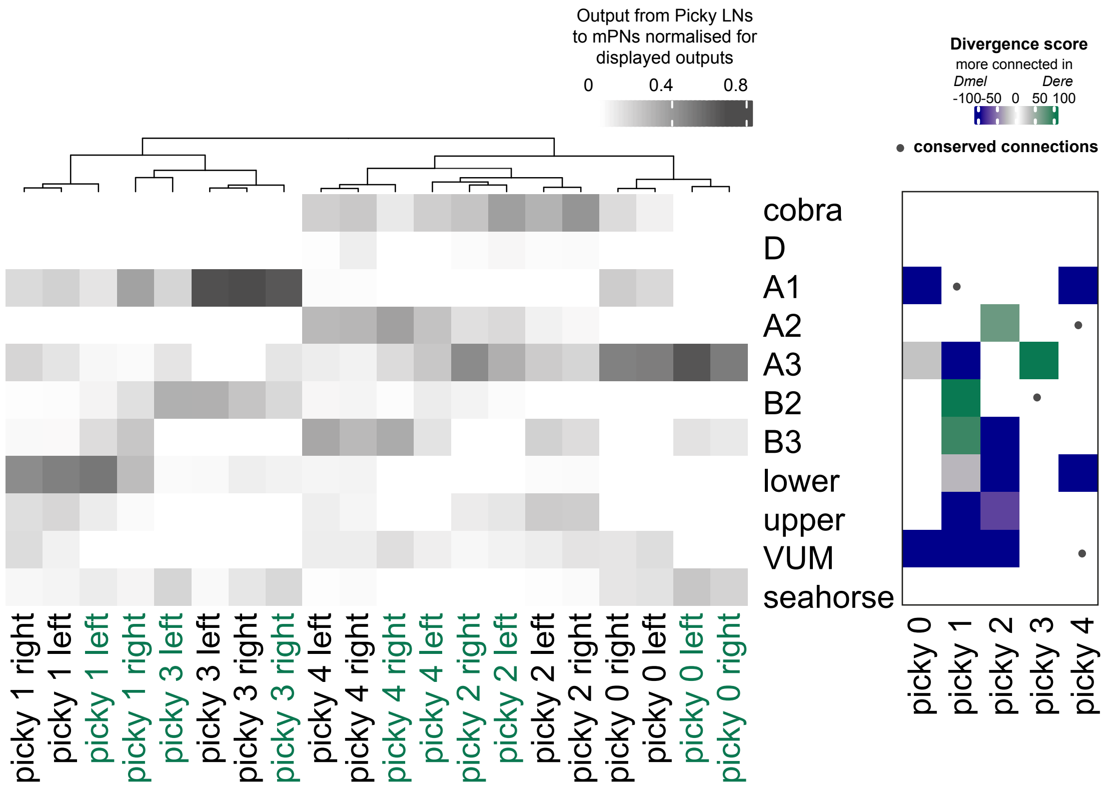

Output from Picky LNs to mPNs normalised for displayed outputs
0
0.4
0.8
Divergence score
more connected in
Dmel
Dere
-100
-50
0
50
100
conserved connections
cobra
D
A1
A2
A3
B2
B3
lower
upper
VUM
seahorse
picky 1 right
picky 1 left
picky 1 left
picky 1 right
picky 3 left
picky 3 left
picky 3 right
picky 3 right
picky 4 left
picky 4 right
picky 4 right
picky 4 left
picky 2 right
picky 2 left
picky 2 left
picky 2 right
picky 0 right
picky 0 left
picky 0 left
picky 0 right
picky 0
picky 1
picky 2
picky 3
picky 4

Figure S19

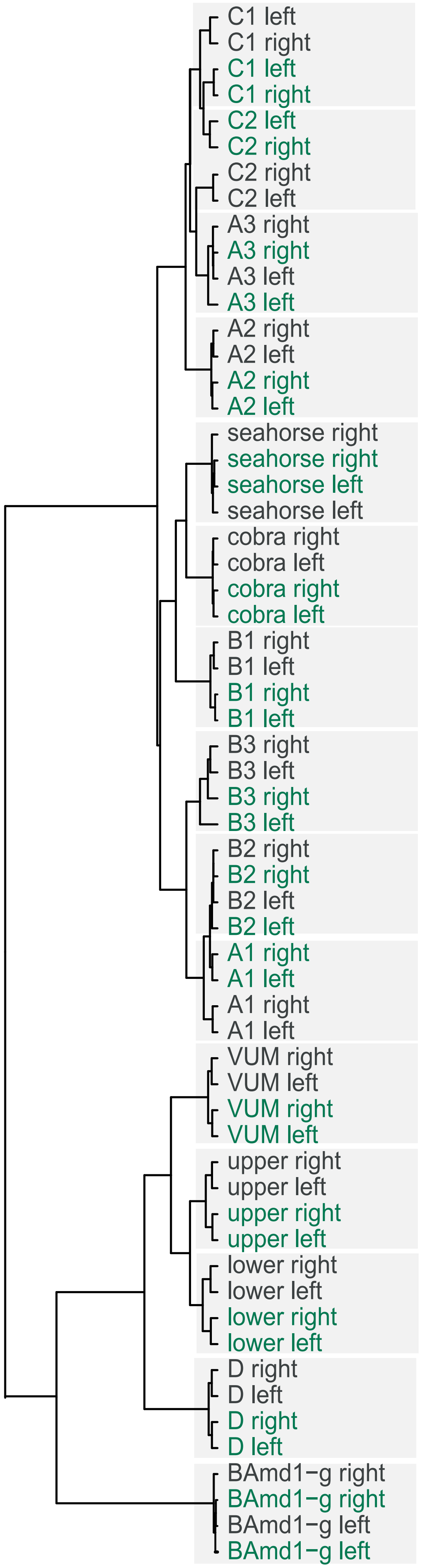
C1 left
C1 right
C1 left
C1 right
C2 left
C2 right
C2 right
C2 left
A3 right
A3 right
A3 left
A3 left
A2 right
A2 left
A2 right
A2 left
seahorse right
seahorse right
seahorse left
seahorse left
cobra right
cobra left
cobra right
cobra left
B1 right
B1 left
B1 right
B1 left
B3 right
B3 left
B3 right
B3 left
B2 right
B2 right
B2 left
B2 left
A1 right
A1 left
A1 right
A1 left
VUM right
VUM left
VUM right
VUM left
upper right
upper left
upper right
upper left
lower right
lower left
lower right
lower left
D right
D left
D right
D left
BAmd1−g right
BAmd1−g right
BAmd1−g left
BAmd1−g left

Figure S20

a

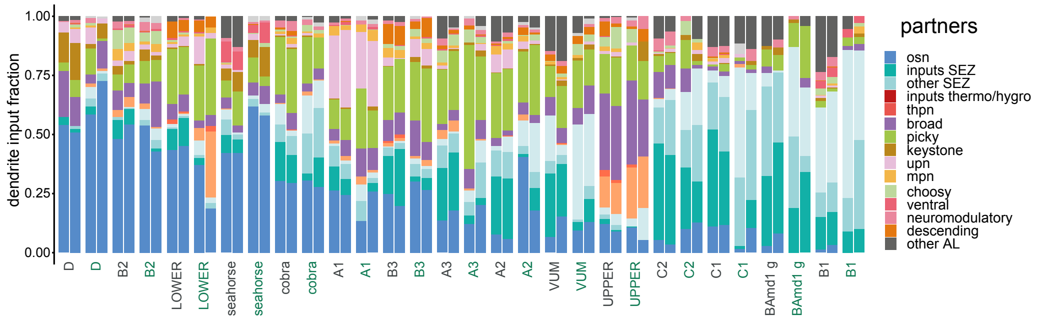


b

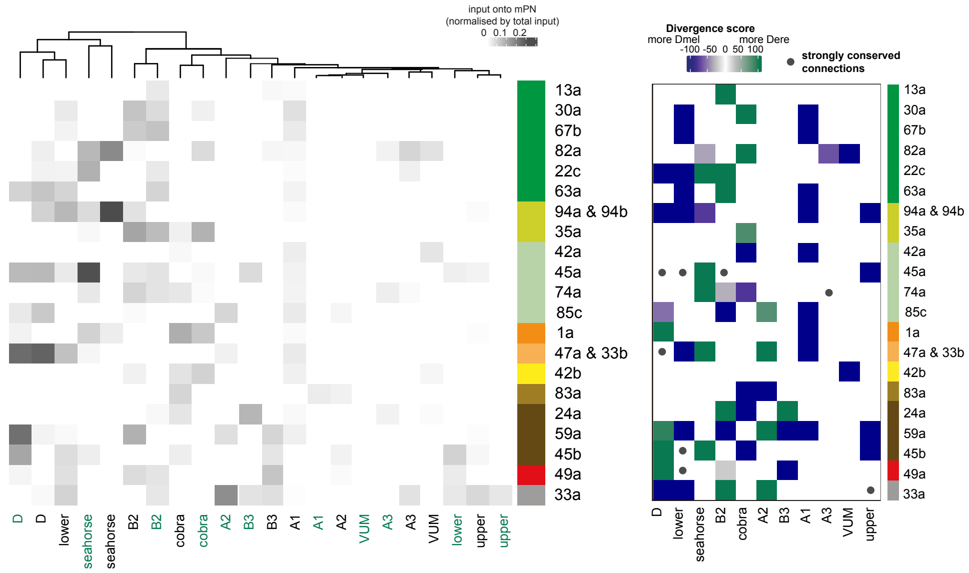

### Figure S21

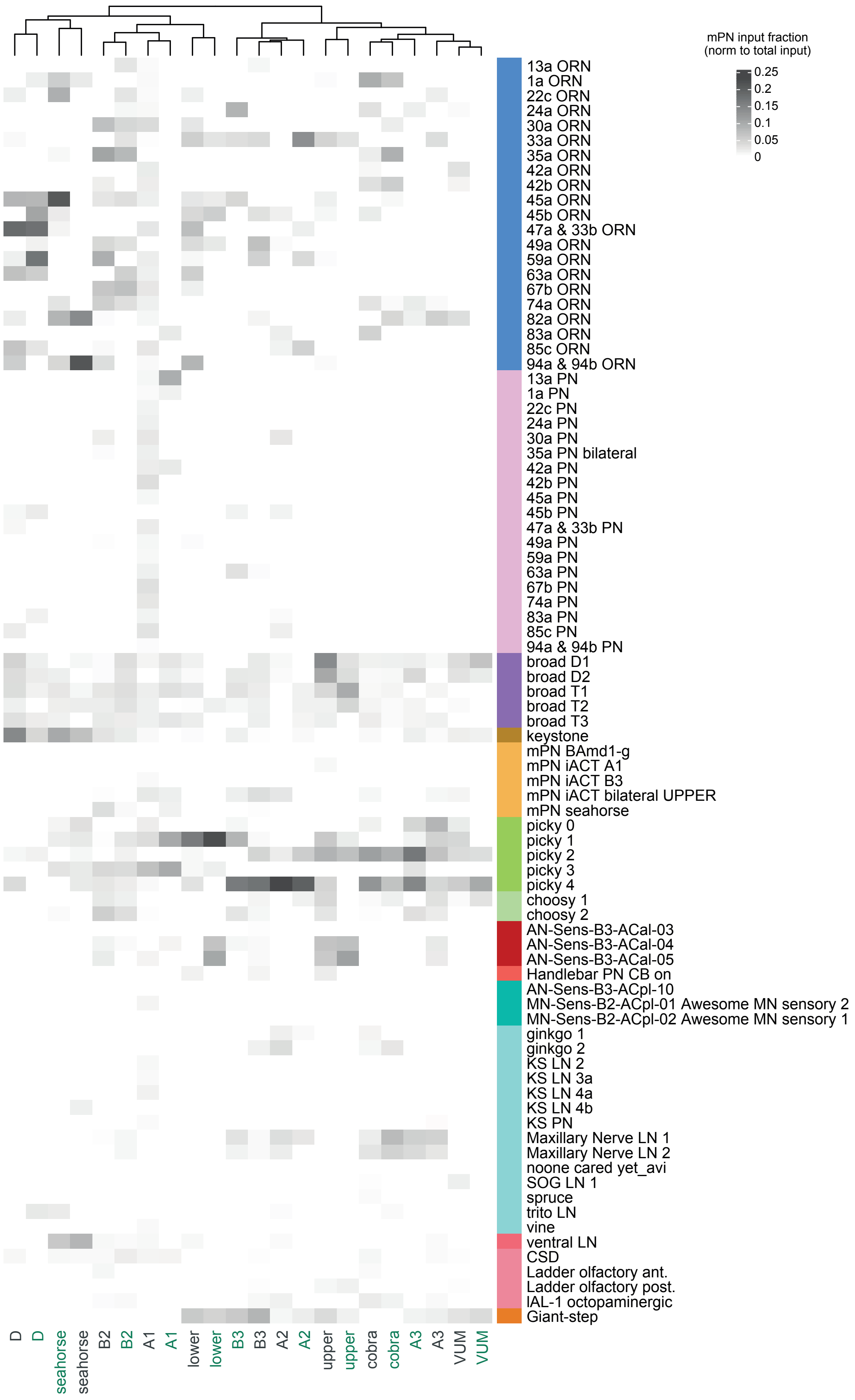
mPN input fraction
(norm to total input)
0.25
0.2
0.15
0.1
0.05
0
13a ORN
1a ORN
22c ORN
24a ORN
30a ORN
33a ORN
35a ORN
42a ORN
42b ORN
45a ORN
45b ORN
47a & 33b ORN
49a ORN
59a ORN
63a ORN
67b ORN
74a ORN
82a ORN
83a ORN
85c ORN
94a & 94b ORN
13a PN
1a PN
22c PN
24a PN
30a PN
35a PN bilateral
42a PN
42b PN
45a PN
45b PN
47a & 33b PN
49a PN
59a PN
63a PN
67b PN
74a PN
83a PN
85c PN
94a & 94b PN
broad D1
broad D2
broad T1
broad T2
broad T3
keystone
mPN BAmd1-g
mPN iACT A1
mPN iACT B3
mPN iACT bilateral UPPER
mPN seahorse
picky 0
picky 1
picky 2
picky 3
picky 4
choosy 1
choosy 2
AN-Sens-B3-ACal-03
AN-Sens-B3-ACal-04
AN-Sens-B3-ACal-05
Handlebar PN CB on
AN-Sens-B3-ACpl-10
MN-Sens-B2-ACpl-01 Awesome MN sensory 2
MN-Sens-B2-ACpl-02 Awesome MN sensory 1
ginkgo 1
ginkgo 2
KS LN 2
KS LN 3a
KS LN 4a
KS LN 4b
KS PN
Maxillary Nerve LN 1
Maxillary Nerve LN 2
noone cared yet_avi
SOG LN 1
spruce
trito LN
vine
ventral LN
CSD
Ladder olfactory ant.
Ladder olfactory post.
lAL-1 octopaminergic
Giant-step
D
D
seahorse
seahorse
B2
B2
A1
A1
lower
lower
B3
B3
A2
A2
upper
upper
cobra
cobra
A3
A3
VUM
VUM
